# Pan-genome and Haplotype Map of Cultivars and Their Wild Ancestors Provides Insights into Selective Evolution of Cassava (*Manihot esculenta* Crantz)

**DOI:** 10.1101/2023.07.02.546475

**Authors:** Zhiqiang Xia, Zhenglin Du, Xincheng Zhou, Sirong Jiang, Tingting Zhu, Le Wang, Fei Chen, Luiz Carvalho, Meiling Zou, Luis Augusto Becerra López-Lavalle, Xiaofei Zhang, Liangye Xu, Zhenyu Wang, Meili Chen, Binxiao Feng, Shujuan Wang, Mengtao Li, Yuanchao Li, Haiyan Wang, Shisheng Liu, Yuting Bao, Long Zhao, Chenji Zhang, Jianjia Xiao, Fenguang Guo, Xu Shen, Cheng Lu, Fei Qiao, Henan Ceballos, Huabin Yan, Huaifang Zhang, Shuang He, Wenmin Zhao, Yinglang Wan, Yinhua Chen, Dongye Huang, Kaimian Li, Bin Liu, Ming Peng, Weixiong Zhang, Birger Muller, Xin Chen, Ming Cheng Luo, Jingfa Xiao, Wenquan Wang

**Affiliations:** Hainan University, Haikou 570228, China; National Key Laboratory of Biotechnology and Breeding of Tropical Crops; National Genomics Data Center, Beijing Institute of Genomics, Chinese Academy of Sciences and China National Center for Bioinformation, Beijing 100101, China; Institute of Tropical Biosciences and Biotechnology, Chinese Academy of Tropical Agricultural Sciences (CATAS), Haikou 571101, China; Department of Plant Sciences, University of California, Davis, California 95616, USA; Tropical Crop Genetic Resources Institute, Chinese Academy of Tropical Agricultural Sciences (CATAS), Haikou 571700, China; Brazilian Enterprise for Agricultural Research (EMBRAPA), Genetic Resources and Biotechnology, Brasilia 70770, Brazil; International Center for Tropical Agriculture (CIAT), Cali 6713, Colombia; Institute of Nanfan & Seed Industry, Guangdong Academy of Sciences, Guangzhou, 510316, China; State Key Laboratory for Conservation and Utilization of Subtropical Agro-bioresources/College of Life Science and Technology, Guangxi University, Nanning 530004, China; The Academy of Agriculture and Forestry Sciences, Qinghai University, Xining 810016, China; Economic Crop Institute, Guangxi Academy of Agricultural Sciences, Nanning, China; Department of Health Technology and Informatics, Department of Computing, The Hong Kong Polytechnic University, Hong Kong, China; Plant Biochemistry, Department of Plant and Environmental Sciences, University of Copenhagen, DK-1871 Frederiksberg C, Denmark

**Keywords:** Genome, Pan-genome, Evolution, Structure Variation, Selective sweeping, Photosynthesis, Storage root formation, Cyanide metabolism, Cassava

## Abstract

Cassava is the most important starch sources, a tropical model crop. We constructed nearly T2T genomes of cultivar AM560, wild ancestors FLA4047 and W14, pan-genome of 24 representatives and a clarified evolutionary tree with 486 accessions. Comparison of SVs and SNVs between the ancestors and cultivated cassavas revealed predominant expansion, contraction of genes and gene families. Significantly selective sweeping occurred in the cassava genomes in 122 footprints with 1,519 candidate domestication genes. We identified selective mutations in *MeCSK* and *MeFNR3* promoting photoreaction associated with *MeNADP-ME* of C_4_ assimilation in modern cassava. Co-evolved retardation of floral primordia and initiation of storage roots arose from *MeCOL5* mutants with altered bindings to *MeFT1, MeFT2* and *MeTFL2. MebHLHs* evolved to regulate the biosynthesis, transport and endogenous remobilization of cyanogenic glucosides, with new functionalities of *MeMATE1, MeGTR* in selected sweet cassava. These findings enhanced comprehensive knowledge and database on the evolution and breeding of cassava.

**HIGHLIGHTS:**

1. Three nearly T2T cassava genomes of cultivar AM560 and its wild ancestors FLA4047 and W14.
2. A species-level cassava panSV haplotype map across 346,322 structural variations over 31,362 gene families and 96,032,008 SNPs and InDels variations globally and a clarified evolutionary tree with 486 accessions.
3. Selective mutations in *MeCSK* and *MeFNR3* promoted photoreaction associated with *MeNADP-ME* of C_4_ assimilation shaped the C_3_-C_4_ intermediate photosynthesis of modern cassava.
4. Coevolution of floral primordia contrary to initiated storage root is pivotal for the domestication of cassava, and arose from *MeCOL5* mutants altered the binding with *MeFT1, MeFT2* (SP6A), and *MeTFL2*.

## INTRODUCTION

Cassava (*Manihot esculenta* Crantz) is a staple food crop belonging to the spurge family (*Euphorbiaceae*) and known for producing numerous large storage tubers enriched in starch. Characterized as tolerance to high light, heat, drought and barren soil, it will be an ideal future crop to combat global climate change. Compared with other major food crops, cassava is attractive due to its tremendous production potential, likely reaching a fresh root yield of 45-60 tons/ha on average or 75-90 tons/ha under favorable conditions. Paleontological records of starch grains and palynology indicated that the origin and domestication of cassava took place 7500-9000 years ago in the Southwest Amazon (Allem, 1994; Aorega *et al*., 2006; Isendahl, 2011), a region regarded as a center of early Amazonian civilization and agriculture fostering domestication of squash, beans, and guava. Cassava fossils from the Late Holocene excavated from Mayan ruins in Meso-America demonstrate that cassava is one of the earliest crops domesticated in South America with an important role in American civilization (Bartlett, *et al*., 1969; Gibbons,1990).

Ninety-eight wild species of cassava have been identified predominantly being distributed in Brazil and the surrounding regions. Among these *Manihot esculenta* ssp. *flabellifolia* (FLA) is a small climbing shrub distributed at the boundary between the tropical rainforest and savanna in the Amazon Basin. It is is considered the most recent ancestor of cultivated cassava (Allem, 1999; Carvalho *et al*., 2013). Evidence from structural variations of a single-copy gene (G3pdh) suggested, that cassava was originally domesticated in the Acre, Rondonia, and Matogrosso states in southern Brazil, where an abundant number of ancestral species and landraces are presently found (Olsen *et al*., 1999). However, it is viable to postulate that cassava domestication occurred in dryland, which is pivotal for storage root formation and promotion of C_4_ photosynthesis. Recently, evidence of cassava producing storage roots in regions near the Andes highland already 10,350 years ago, has been provided based on ^14^C isotope measurements (Lombardo *et al*., 2020).

Cultivated cassava is capable of producing a large quantity of biomass even under harsh conditions of drought, high-intensity sunlight, and hyperthermia (Carvalho *et al*., 2013). These favorable agronomical traits of cultivated cassava are derived from their wild ancestors by adaptation to environmental challenges and artificial selection (Olsen and Wendel, 2013; Alonge *et al*., 2020) as manifested in the expansion of genes and gene families that contribute to the traits. The genomic expansion and shrinkage processes that happened in the course of evolution can now be assessed using genomic technologies, e.g. by sequencing the genomes and constructing a pan-genome of the wild cassava species and cultivars and by re-sequencing and mining alleles associated with domesticated traits (Liu *et al*., 2020; Zhang *et al*., 2020; Varshney *et al*., 2021).

We previously reported the genome sequences of cassava cultivar KU50 and the wild ancestor W14 (Wang *et al*., 2014). Subsequently, the genome sequences of cultivar AM560 (Bredeson *et al*., 2015), SC205 (Hu *et al*., 2021) and TME204 (Qi *et al*., 2022) were obtained using PacBio sequencing and improved Hi-C sequencing. These genomic resources have made it possible to gain insight into the evolution of favorable traits. For example, by comparing the genomes of the wild ancestor W14 and the cultivar KU50, we detected variations of genes and gene families involved in starch accumulation, biosynthesis of the cell wall and of natural products (specialized metabolites)such as cyanogenic glucosides, demonstrating the rigor of genomic technologies (Wang *et al*., 2014). A total of 203 selective sweeping regions including 427 genes has also been reported, many of which are involved in sugar and starch metabolism and stress tolerance (Ramu *et al*., 2017). However, no species-level analysis has been performed so far to compare wild ancestors and modern varieties of cassava or seldom done in other economic crops.

Domesticated cassava possesses C_3_-C_4_ intermediate photosynthesis with a significantly higher photosynthetic carbon fixation rate and a more efficient responses to light intensity compared with its wild relatives (EI-Sharkawy, 1987; Bräutigam *et al*., 2016; CALATAYUD *et al*., 2002). For example, pyruvate orthophosphate dikinase (*PPDK*), phosphoenolpyruvate carboxykinase (*PEPC*), and NADP-dependent malic enzyme (*NADP-ME*), which are representatives of C_4_ carbon assimilation ([Mercado](https://pubmed.ncbi.nlm.nih.gov/?term=Mercado+MA&cauthor_id=35231181) and [Stude](https://pubmed.ncbi.nlm.nih.gov/?term=Studer+AJ&cauthor_id=35231181), 2022) are all expressed abundantly in functional leaves of cultivated cassava (Sharkawy,2004; our unpublished data). The evolution of these gene families from wild ancestors to cultivated cassava is unknown, and the characteristics of photochemical reactions in C_4_ species have not yet been fully explained (Munekage, 2016; Mercado and Stude, 2022). The knowledge will be crucial for understanding the photosynthetic mechanisms in cassava and other plants (Schlüter *et al*. 2020).

The coevolution of flowering and storage root formation is another key event in the speciation of modern cassava. Ancestral species blossom early and produce a large number of fruits and seeds, but carries tiny storage roots. Cultivars typically have delayed flowering and produce few fruits, but develop large storage roots and use nutrient stems as seeds for reproduction. Such drastic and essential changes transferred nutrition reservoirs from fruits and seeds to stems and storage roots consequentially promoted starch accumulation (Carvalho *et al*., 2013). The florigen (*FT*) gene family in plants plays an essential role in plant flowering and vegetative organ formation. Members of the family, *FT1* and *FT2* are expressed in specific companion cells in leaves induced by environmental signals, and are transported remotely to the apical meristem or the tips of adventitious roots in the form of mRNA or protein to initiate development of floral primordiaa or tubers (Corbesier, 2007; Navarro,2011; Hoang, 2020). A third member, *TFL2* is a flowering suppressant and essential for enlargement of vegetative organs. In cassava, there are ten members of the *FT* family (Adeyemo *et al*. 2019), but no clear structural variations between wild ancestor to cultivars have been identified except for *MeFT2 and MeTFL2*. Interestingly, a sequence deletion has been found in a *CONSTANS* gene (*MeCOL5*), which is regarded as a co-regulator of *FTs* (Freytes *et al*, 2021; Zierer *et al*., 2021). The identity of the swich controlling the balance between flowering or storage root development thus remains elusiveness.

Cyanogenic glucosides are a class of natural products which initially evolved in Pteridophytes ∼320 million years ago and are present in many higher plants (Zagrobelny *et al*, 2008; Gleadow and Møller, 2014). Cyanogenic glucosides serve as remobilizable storage forms of reduced nitrogen (Picmanova et al, 2015; Bjarnholt et al, 2018), but are mostly known for their function as defense compounds against herbivores and pests (Tattersall *et al*, 2001; Zagrobelny and Møller, 2011). Cassava contains two cyanogenic glucosides linamarin and lotaustralin. When used as a food source, cassava leaves and tubers need to be processed to avoid exposure to toxic hydrogen cyanide upon consumption. In cassava, linamarin and lotaustralin are synthesized mainly in the leaves and transported to the storage roots through the phloem (Jørgensen *et al*., 2005). The content of cyanogenic glucosides in storage roots has been decreased during the domestication process. The genomic resources within the population of wild and cultivated cassava represent a route to understand and further augment the domestication process.

In this study, we established complete genome sequences of two wild ancestral cassava sub-species FLA4047 and W14 as well as of the cultivar AM560 and pan genome of 24 selected genotypes using a combination of PAC-Bio, BioNano, and Nanopore sequencing technologies. We also re-sequenced a total of 486 varieties, landraces, and wild relatives to build a Pan SV genome and an evolutionary tree of cassava. The large collection of genome sequencing data from a variety of cassava species enabled resolution of a large number of SVs, SNPs, and InDels, outlining the complexity, diversity, and evolutionary history of the cassava genome. Using selective sweeping analysis, we identified the majority of SV and SNP mutations associated with domestication events. Based on the complete reference genomes, pan-genome, and genome variations in about 500 accessions, we unraveled the evolutionary trajectory from progenitor to cultivars and revealed new knowledge on several key domestication events in cassava e. g. in evolution of C_3_-C_4_ photosynthesis, coevolution of flowering and storage root formation, and in cyanogenic glucoside biosynthesis which are essential for cassava and of key relevance to domestication of other potential tropical crop plants and support the validity of the predictions by biological experiments.

## RESULTS

### Complete Genomes of Cultivar AM560 and Wild Ancestors FLA4047 and W14

A selfing S3 clone AM560 with low heterozygosity as a representative cultivated cassava (*Manihot esculenta*), an accession of wild ancestor FLA4047 (*Manihot esculenta* ssp. *flabellifolia*) and another wild relative W14 were selected for genome sequencing (**Fig.S1**). Their estimated genome sizes were 720Mb, 720Mb, and 830Mb, respectively (**Fig.S3, Table S1**,). The basic frameworks of the genomes were constructed by integrating multiple next- and third-generation sequencing technologies and *de novo* assembly (**Fig.S4, Table S2**). Two cells of nanopore Ultra-long reads at ∼170X depth for each genome were used to assemble into contigs using Nextdenovo (**Fig.S4**) and resulted in three nearly completely sequenced genomes. The genome of cultivar AM560 was assembled into a 645 Mb reference genome with no gaps in 16 of the 18 chromosomes, three gaps on Chr3 and 2 gaps on Chr 18. The FLA4047 genome was assembled into 672Mb in size with 1 to 9 gaps in 15 of the 18 chromosomes. The genome of W14 was assembled into 709 Mb in size without gaps in 10 of the 18 chromosomes and 1 to 10 gaps in the remaining 8 chromosomes with a total of 33 gaps. The scaffold N50 of the three genomes ranged from 35 to 40 Mb. The completeness and accuracy of the genome assemblies of AM560, FLA4047, and W14 were assessed by BUSCOs to reach 97.7%, 96.9%, and 97.5%, respectively (**Table 1**,**Table S3**). The sequencing and assembly results showed high integrity and accuracy of the genome assemblies. A high percentage of repeats appeared in the cassava genomes, ranging from 63.7% to 67.9%. The most abundant type of repeat is Gypsy, a type of long terminal repeat (LTR), which accounts for more than 55% of all repeats. After masking and excluding repetitive sequences, coding genes in the three genomes were annotated using homologous protein comparison and gene prediction, and long noncoding RNAs (lncRNAs) were identified using RNA-sequencing data. In total, 30,571, 33,913, and 32,406 protein-coding genes and 12,253, 12,683, and 8,687 lncRNA genes were annotated in the AM560, FLA4047, and W14 genomes, respectively (**Table 1**). All non coding RNAs were accuratedly predicated in **Table S4**.

**Table 1.** Statistics for Assembly and Annotation of the Genomes of AM560, FLA4047, and W14.

| Chr ID | AM560 |  | FLA4047 |  | W14 |  |
| --- | --- | --- | --- | --- | --- | --- |
|  | No. of gaps | Length (bp) | No. of Contigs | Length (bp) | No. of Contigs | Length (bp) |
| Chr1 | 0 | 43,337,869 | 9 | 54,294,512 | 0 | 46,595,211 |
| Chr2 | 0 | 39,653,371 | 1 | 40,267,192 | 0 | 40,768,185 |
| Chr3 | 3 | 33,784,460 | 0 | 39,425,326 | 4 | 42,268,827 |
| Chr4 | 0 | 36,239,750 | 2 | 35,416,046 | 0 | 35,710,473 |
| Chr5 | 0 | 33,752,819 | 7 | 47,532,515 | 5 | 40,270,531 |
| Chr6 | 0 | 31,220,583 | 6 | 33,926,122 | 4 | 35,748,825 |
| Chr7 | 0 | 35,869,022 | 4 | 35,975,965 | 0 | 42,243,325 |
| Chr8 | 0 | 42,184,545 | 4 | 40,590,130 | 4 | 44,818,185 |
| Chr9 | 0 | 39,139,566 | 2 | 37,853,125 | 0 | 42,675,627 |
| Chr10 | 0 | 32,276,928 | 1 | 36,384,499 | 0 | 32,699,422 |
| Chr11 | 0 | 33,666,177 | 0 | 33,978,540 | 10 | 47,379,937 |
| Chr12 | 0 | 38,689,200 | 0 | 32,432,365 | 0 | 43,693,216 |
| Chr13 | 0 | 37,027,919 | 7 | 35,451,318 | 0 | 38,835,027 |
| Chr14 | 0 | 29,478,005 | 5 | 33,542,330 | 0 | 36,926,023 |
| Chr15 | 0 | 33,649,108 | 3 | 30,579,525 | 0 | 32,946,348 |
| Chr16 | 0 | 34,124,650 | 2 | 34,960,364 | 4 | 42,388,355 |
| Chr17 | 0 | 36,189,752 | 5 | 34,881,022 | 1 | 32,988,721 |
| Chr18 | 2 | 35,115,907 | 2 | 32,667,266 | 1 | 30,941,161 |
| No. of contigs |  | 21 |  | 91 |  | 51 |
| Contig N50 (Mb) |  | 35,869,022 |  | 19,815,018 |  | 32,946,348 |
| Scaffold N50 (Mb) |  | 36,121,210 |  | 35,451,318 |  | 40,768,185 |
| Total Length of assembly |  | 645,399,631 |  | 672,347,788 |  | 709,897,399 |
| Complete BUSCOs (%) |  | 97.7 |  | 96.9 |  | 97.5 |
| Repeats (%) |  | 65.70 |  | 63.73 |  | 67.86 |
| Protein Coding genes |  | 30,571 |  | 33,913 |  | 32,406 |
| LncRNA number |  | 8,687 |  | 12,253 |  | 12,683 |

### Evolution and Phylogeny of Cassava Ancestral and Cultivated Subspecies

The synteny of chromosome pairs across the three cassava genomes was assessed using single-copy homologous (SCH) genes (**Fig. 1A**). There exist 8,903 SCH genes between AM560 and A4047, 10,314 SCH genes between AM560 and W14, and 10,173 SCH genes between W14 and A4047. High synteny between wild and cultivar species is observed, with the minor interruptions being assigned to the identified low numbers of inter-chromosome rearrangements, translocations and inversions (**Fig. 1A; Fig. S7-8**). A comparison of the three cassava genomes identified 16,796 conserved genes as well as 646 unique genes for W14, 157 for A4047, and 819 for AM560 (**Fig. S9**). A GO enrichment analysis of the unique genes of wild and cultivated varieties showed that these were significantly enrichedwith functions assigned as catalytic activities and metabolic processes. The evolutionary relationship of the three cassava varieties and a dozen species in *Euphorbiaceae* and outgroups was estimated by the synonymous substitution rates (Ks) of single-copy ortholog genes (**Fig. 1C**). The speciation of three *Euphorbiaceae* species, hevea, jatropha, and castor bean, occurred around 21, 23 and 25 MYA, respectively. The W14, representative species of *M. esculenta* differentiated from them at about 20 million years ago (MYA). FLA4047 and cultivated AM560 were separated from W14 about 0.2 and 0.02 MYAs, respectively. FLA4047 was the closest wild cassava to the cultivated variety AM560. (**Fig.1D**).

**Fig. 1.**
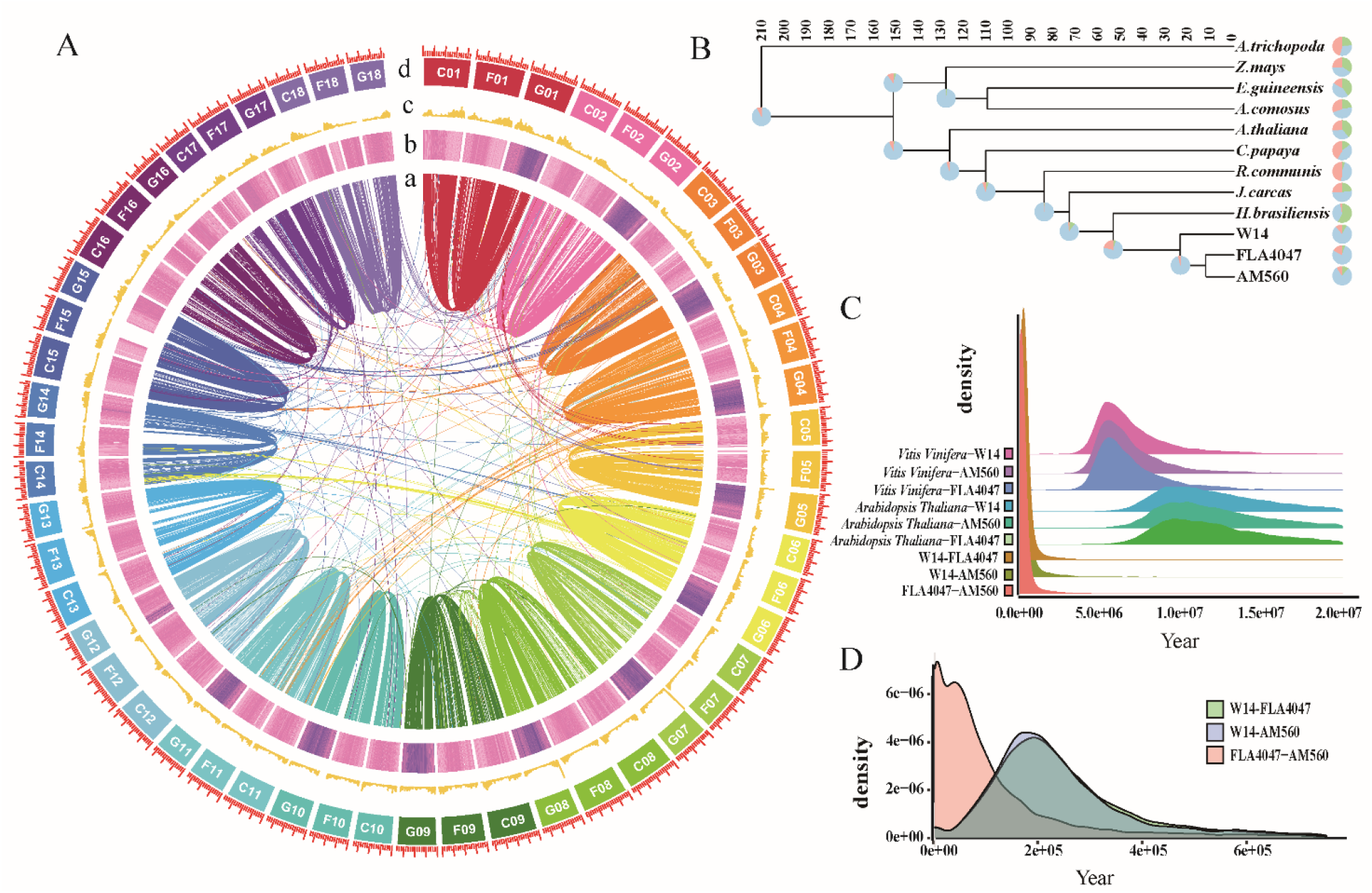
Structural Comparison of The Three Cassava Genomes|. **A**. Chromosomal synteny of genome AM560, FLA4047, and W14. (a) Chromosome karyotypes. (b) Gene density. (c) TEs. (d) Chromosomal synteny among the three genomes. C01-18,AM560;F01-18,FLA4047;G01-18,W14; **B**. Sequential evolutionary relationships between cassava and rubber tree, Jatropha, castor bean, papaya, pineapple, oil palm, maize, and *Arabidopsis thaliana*. As shown, FLA4047 is the nearest wild relative of cultivated cassava. **C**. Estimated evolutionary time of cassava in comparison with grape and *Arabidopsis thaliana*. Concretely, the species or subspecies differentiation from W14 to FLA4047, W14 to AM560, and FLA4047 to AM560 rank in about 0.20, 0.18, and 0.02 MYA, respectively. **D**. Distribution of Ks values among W14, FLA4047 and AM560.

### Pan SV Genome of 24 Representative Genotypes in Cassava

Of the 486 accessions, 21 were chosen for pangenome construction. The selection included 20 cultivars from different continents and the wild species GLZyn (*Manihot glazovii*). Each accession was subjected to ONT (Oxford Nanopore Technologies) long-read sequencing to generate a minimum of 50x genome coverage. This enabled high-quality genome assembly with a contig N50 of 5-10 Mb and total genome sizes between 638-843Mb (**Table S5**) and prediction of 30,245 to 41,950 protein-coding genes (**Table S6**). Following inclusion of the available three complete reference genomes, a pan genome with 24 representive cassava genotypes was constructed (**Fig. 2**). The integrated orthologous genes can be classified into 31,362 families with about 47,000 gene models (**Supplementary Table S17**). The number of gene families increased with the number of genomes and was saturated at 22 genomes (**Fig. 2A**), suggesting that the pan genome of 24 accessions provides a proper representation of the genomes of the cassava species. Overall, 6,503 gene families (∼21% of all gene family) appeared in all genomes and were assigned as core genes. Similarly, 4,794 families (15%) were softcore gene families, 19,417 families (∼62%) were dispensable gene families. A total of 652 gene families were found only in single genome and were assigned as private genes. The latter family accounted for ∼ 2% of all gene families (**Fig. 2 B, Table S7**). A KEGG pathway analysis revealed that the core genes were enriched primarily in metabolic pathways, spliceosome and proteasome (**Fig. S13a**), the softcore genes enriched in mismatch repair, non-homologous end-joining and RNA degradation (**Fig. S13b**), and the dispensable genes enriched in RNA degradation, plant-pathogen interaction and ribosome biogenesis in eukaryotes (**Fig. S13c**). The private genes were mainly ascribed into pentose and gluconate conversions, valine, leucine and isoleucine degradation, and butanoate metabolism (**Fig. S13d**).

**Fig. 2.**
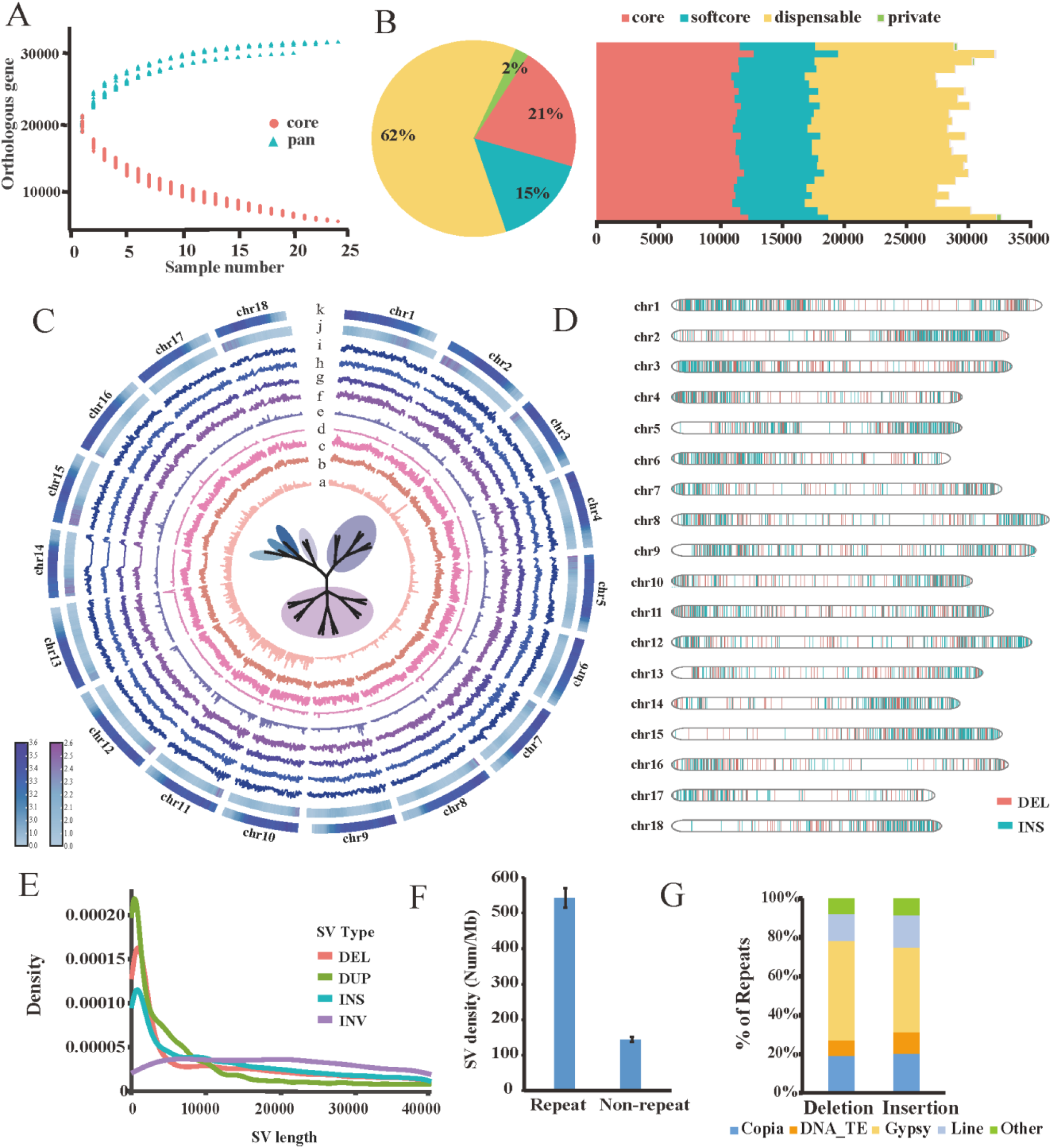
Pan SV-genome of 24 Cassava Accessions |. **A**. Variation of gene families in the pan and core genome. **B**. Proportions of core, softcore, dispensable and private gene families. **C**. Pan SV genome in 18 chromosomes. (a) Inversions. (b) Duplications. (c) Translocations. (d) Deletions. (e) Insertions. (f) SNPs distribution in genome of wild accessions (GLZyn, W14, FLA4047). (g) SNPs distribution in ME population1 (IB005 and ZM9781). (h) SNPs in ME population2 (BC006, BC008, Fuxuan01, HB60, ITBB04 and Mianbao). (i) SNPs in ME population3 (Baxi-1, GR4, IB003, NZ199, SC205, SC5, SC6068, SC8, SC9, Taiyin01, Wenchang red and Yinni). (j) Gene model distribution. (k) Repeats distribution in 18 chromosomes. **D**. Chromosomal distribution of INSs and DELs co-evolved in more than 15 accessions referenced to wild ancestor W14. **E**. Histogram of SV sizes for deletions (DEL), duplications (DUP), insertions (INS), and Inversions (INS). The TRA type is the breakpoint of the swap, so no length is drawn SV in different varieties. **F**. SVs distributed in repeats and in non-repeat sequences based on InDels at least 50 bp in size. **G**. Stacked bar graph showing the proportional distribution of Repeat Types in SVs, including all annotated insertions and deletions. TE accounts for more than half of the SVs among the insertions as well as and deletions.

Taking the AM560 genome as a reference, the structural variants were extracted from 21 cultivars and three wild accessions using Sniffles software, respectively (**Fig. 2C**). A total of 346,322 SV events occurred in cultivated varieties, among which Deletions (DEL) and Insertions (INS) accounted for 51.01% and 23.32% of the total SVs with scattered distribution across 18 chromosomes (**Fig.2D**), followed by Translocations (TRA, 18.33%) and Duplications (DUP, 5.5%) (**Fig. S11, Table S8**). The DEL, DUP and INS were relatively short, typically ranging from 50-200bp, although some were up to 40Kb (**Fig. 2E**). A large proportion (78.7%) of the SVs appeared in repeat regions (**Fig. 2F**), with the bulk being DEL and INS variants such as Gypsy and Copia (**Fig. 2G**). the number of INS ranged from 3915 to 7878, with a minimum sum of INS size being 95.30 Mb and the maximum sum of INS size being 161.42 Mb. The average insertion size was 18.7 Kb (**Fig. S12**). This indicates that selection within repetitive sequences has played a major role in the evolution of the cassava genomes.?

### Gene Expansion and Contraction between FLA to ME

A systematic comparison of gene models of the three wild cassava ancestors (FLA) and 21 cultivars (ME) revealed that the wild ancestral genomes harbored 382 homologous gene families with a total of 1,015 members, which were lost to varying degrees in the modern cultivated varieties. The genomes of the modern cultivated varieties contained 6,812 gene families involving 34,241 genes not present in the cassava ancestors (**Supplementary Table S17**). This demonstrates that a large number of new genes were acquired in the course of natural selection and artificial domestication. A comparative GO pathway analysis between wild ancestors and cultivated cassava varieties showed substantial differences in the gene content related to metabolic pathways. In the wild progenitors, the genes were mainly related to environmental adaptations like biosynthesis of natural products, photosynthesis, cysteine and methionine metabolism. In comparison, the genes and gene families expanded in cultivated cassava were involved in biosynthesis of natural products, starch and sucrose metabolism, amino sugar and nucleotide sugar metabolism, protein processing in endoplasmic reticulum, and RNA transport. In addition, a MAPK signaling pathway, ABC transporters and fatty acid degradation were also enriched in the cultivars (**Fig. S14, Table S17**).

### Copy Number Variations (CNVs)

We compared the copy number variations (CNVs) of the genes and gene families between the 3 wild ancestors and the 21 cultivars. CNVs were frequent within photosynthesis, starch and sugar metabolism, hormone metabolism and signaling, and within some of the transcription factor families. In photosynthesis, 13 gene families in 30 KEGG pathways in the photosynthetic light reaction were expanded to different degrees in the cultivars. On the contrary, 12 contracted gene families were found in the ATP synthase subunits (*atpH* (K02113), *atpE*(K02114)) and in photosystem I P700 Apoprotein A1 (*psaA* (K02689)) etc. (**Fig. S15-16, Table S18**). In sucrose and starch metabolism, a large number of CNVs were found in 17 of 31 gene families involved. Gene expansion significantly happened in alpha-1,4 glucan phosphorylase L-2 isozyme (K00688), alpha-glucosidase-like (*STP-1*, K01187), sucrose synthase (K00695) etc. The remaining 14 gene families showed some shrinkage in cultivated species (**Fig.S17-18; Supplementary Table S18**). The increased CNVs in sucrose and starch metabolism may be the driving factors of the high biomass and starch accumulation in the storage roots in cultivated cassava.

We analyzed the CNVs between the three wild and 21 cassava cultivars within a total of 42 gene families in nine plant hormone biosynthesis and signal pathways. Several significant gene expansion events were found in basal regulation, indol acetic acid, cytokinin, salicylic acid and the ethylene biosynthetic pathways. Other gene families were contracted to different degrees (**Fig. S20, Supplementary Table S18**). The contractions observed in the major transcription factor families were remarkable. The 58 families of transcription factors present in the analyzed cassava genomes showed overall gene contraction and also tended to be subjected to a decrease gene copy numbers in the course of domestication from wild ancestor to cultivars. The thirteen transcription factor families minimized with between 4 and 26 copies include *bHLH, MYB, ERF, NAC, C2H2, WRKY, bZIP05, HD-Zip*.. Only *HSF* showed a slight expansion in gene number. The cumulative cultivated/wild CNV ratio was 0.91 for all 58 TFs (**Fig. S20, Supplementary Table S18**).

### Phylogeny and genetic variation from wild ancestors to cultivated cassava

Based on 30X Illumina short reads resequencing data of 291 accessions along with the published 195 accessions (Ramu *et al*, 2017), we obtained a rich dataset on the genetic variation in 486 different cassava genotypes covering 35 wild relatives (FLA and GLA) and 451 cultivated cassava varieties. The domesticated varieties were collected in Central and South America, Africa, Southeast Asia and China and consist of landraces and improved hybrids (**Fig. 3A, Supplementary Table S15**). A total of 96,032,008 convincing SNP and InDel variations were obtained, with an average of 131.4 base mutations per kb sequence. Of these, 95,581,122 (56.00%) were located in intergenic regions, 2,571,999 (1.507%) in exonic and 10,929,118 (6.40%) in intronic regions (**Supplementary Table S16**).

**Fig.3.**
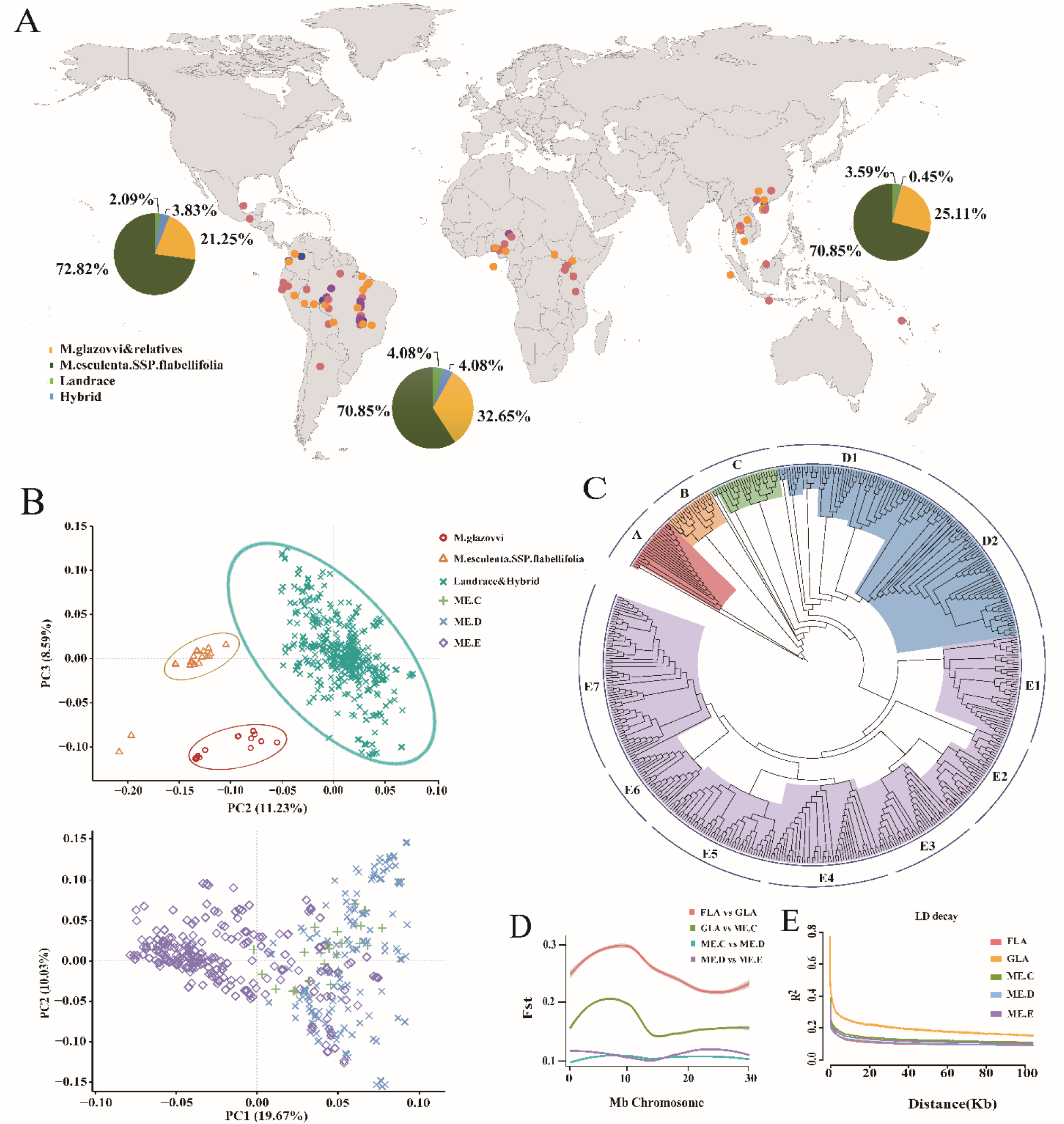
Phylogenetic tree and population domestication revealed by comparison among 486 landraces and cultivars to wild relatives in cassava. **A**. Geographic distribution of cassava accessions; **B**. Principal Component Analysis (PCA) of accessions showing clear differentiation among GLZs and its relatives, FLAs and cultivated clones (MEs) (n=482) (upper panel). PCA of cultivated cassava clones reveals little differentiation between the ME C, ME D and ME E clades (n=450)(lower panel); **C**. Phylogenetic tree and population structure of 486 accessions: All accessions were ascribed into 5 clades, Group A: GLZ and its relatives; Group B: FLA, genotypes in the wild ancestor sub-species; Group C: Landrace and hybrids probably rooted around Ecuador; Group D: with D1 and D2 sub-groups covering landraces and cultivars distributed in Central Brazil where cassava originated. Group E: E1-E3 are accessions mostly derived from Southern Brazil and include cold tolerant accessions; E4-E7 are landraces and hybrids collected in east and west Africa. **D**. The Fst value between GLZs and FLAs is higher than between FLAs and ME C, and even lower between ME C and ME D orME E. **E**. LD decays took place in GLAs, FLAs, ME C,ME D and ME E groups.

A total of 6,021,758 filtered SNPs were retained for construction of a phylogeny of cassava (**Supplementary Table S16**). At K=5, the whole population was divided into five groups (**Fig.3B**) : Group A consists of nineteen *M. glazovii* accessions including Tree Cassava, GLZyn, GLZb, W14 and hybrids with cultivated cassava. The W14, reported in the present study, own the background of GLZ, but PCA cluster analyses imply it is a hybrid with a cultivated species. This is supported by its production of storage roots low in starch (**Fig.3C**) and identifies GLZ as one of the few wild cassava species that exchanged genes with cultivated cassava in the course of natural evolution (Bredeson *et al*., 2016). Group B includes 22 accessions of FLA cassava. The accessions were obtained from several Brazilian states on the southern edge of the Amazon. The positioning of FLA in the phylogeny further augments the conclusion presented based on the genomic sequences that FLA is the closest progenitor of cassava. Group C is the domesticated type closest to FLA, and includes 43 landraces distributed in Ecuador and their hybrid offsprings with cultivars. Ecuador was previously considered the oldest area of human activities and species domestication in America (Isendahl, 2011). Group D is cultivars with two subgroups: D1 consist of 68 varieties, which are landraces and hybrid progeny geographically distributed in central and western Brazil, adjacent to the cassava domestication center (Olsen, *et al*., 1999). D2 consists of 85 genotypes, with sugary cassava, SC5, SC8 and their derived hybrids named the K series. These genotypes, seem to have their origin in the Amazon close to the reported domestication center. Group E covers 248 genotypes and are divided into seven subgroups E1-E7 according to their genetic distance: E1 and E2 contain a total of 56 cultivated clones derived from Southern Brazil, and includes the relatively cold tolerant clones SC124, F201, CH16. E3 contains several specific varieties from the Parana state in Brazil, as well as many edible clones from Asian and Africa countries, assigned to the geographic subtype of southern Brazil; E4 consists of landraces and hybrids from Uganda in Eastern Africa, and a few African collections introduced from Colombia. E5 to E7, 99 represent accessions of all landraces and their hybrid progenies collected from western to eastern Africa. It is indicated that genetic domestication occurred since cassava came into Africa about 450 years ago. PCA analysis show that GLA, FLA and ME hold independent evolutionary positions. Inside ME, only ME-E appears to reveal a domesticated trend (**Fig.3C)**. The genetic differentiation index (Fst) between the phylogenetic clades is highest between GLA and FLA (0.3), less between GLA and ME-C (0.164), and very low between Me-C, Me-D and Me-E (**Fig. 3D, Supplementary Table S16**). The genetic variation inside each population was low (π were 2.46E-05 to 0.0024), the lowest in GLZ, followed by FLA and higher in cultivated populations. The linkage disequilibrium(LD) decay is very slow except for GLA (**Fig.3E**).

### Selective Sweeping in Domestication of Cassava Genome

We analyzed the selective sweeps represented in genomes of 198 cultivars and 28 accessions of the wild ancestor (FLA) using ROD (reduction of diversity), Fst (population genetic differentiation index), and the combination of the two indices (ROD-Fst) with all diversity SNPs and InDels (**Table S9**). For all 18 chromosomes, a sequence sliding window of 100 kb with 10 Kb increments was applied followed by calculation of the ROD and Fst values in each window. We identified the selective sweep regions using the significance criteria 5%, 2%, and 1%. The statistical results revealed a total of 18,006 windows, where the diversity decreased in cultivars compared to the FLA. Within the top 5% ROD value, we identified 162 loci, covering 49.3 Mb over 2150 genes. Only 28 loci were found within the top 2% of ROD, covering 16.4 Mb over 390 genes. Within the top 1% of ROD, 27 regions were identified covering a total of 9.3 Mb and spanning 174 genes (**Supplementary Table S19**). Analyses based on the Fst parameter and the top 5%, 2%, and 1% significance criteria identified 271, 150, and 67 loci, accounting for 60.7 Mb, 27.9 Mb, and 13.2 Mb of the genome and including 2931, 1412, and 652 genes, respectively (**Fig. 3 E; Supplementary Table S20**). The cultivars had larger π values of gene sequence diversity compared with the wild relatives. These differences might relate to the preferred region in cassava domestication.

Integrated index of ROD and Fst helped identify 122 selective sweeping regions with 1519 genes within the top 5% (**Supplementary Table S21**). These genes were distributed across all 18 chromosomes with several hot spots (**Fig. S22**). KEGG and GO annotations showed their involvement in metabolic pathways, including biosynthesis of cofactors (32 genes), plant hormone signal transduction (26 genes), starch and sucrose metabolism (24 genes), virus related (29 genes) and ribosome composition (22 genes) (**Fig.S23**). Several genes with interesting functionalities were uncovered. *MeTFL2* (Manes.C.13G010700) is associated with flowering inhibition and root tuber development in cassava. Chloroplast Censor kinase (*MeCSK*, Manes.C.13G024500) regulates the expression levels of the D1 and D2 proteins in photosystem II, and are functionally linked to the efficient adaptation of cassava to high light conditions. *MeTIR* (Manes.02G147400) is an auxin regulator, as confirmed by population association analysis (Hu *et al*., 2021). We further examined the distribution of 9 types of pivot transposons, including *hAT*-MITE, *mutator*-MITE, *stowaway*-MITE, and SINEs in the promoter region (2,000 bp upstream of ATG) of genes annotated in all 198 cultivars and 28 wild relatives using ITIS software (**Supplementary Table S22**). For most of the cultivars (185/198), a 1053 bp *hAT-*MITE transposable element appeared 7 kb upstream of the coding region of *TFL2* (Manes.C.13G010700). In contrast, no transposon was found in the promoters of *TFL2* in 26/28 wild ancestors. An 83 bp SINE retrotransposon element was inserted in the promoter regions of *CSK* in many cultivars (167/198), but not in any wild relatives except one (**Supplementary S23**).

### Photosynthesis evolution in cultivated cassava

The domesticated cassava is known for C3-C4 intermediate photosynthesis and is characterized by high *Me*PEPC and *Me*PPDK activities resulting in a high net photosynthetic rate (Pn). In contrast, most wild species are C3 type (Calatayud *et al*., 2002). A whole-genome comparison between 298 varieties (ME) and 28 wild ancestor species (FLA) discovered selective mutations in several essential genes in photochemical conversion and carbon fixation pathways. Cultivars (e.g. KU50) were more light tolerant than the wild ancestors (FLA4047, W14), reflected by a significantly increased Pn response to PAM from 500 to 2500 μmol/m^2^s light intensity in plants of DAP120 (**Fig.4A**), while the ATP and NADPH outputs of KU50 were 0.82,This was 1.44 fold higher than FLA4047 and 1.29, 1.53 folds than W14, respectively (**Fig.4B,Supplementary Table S26**). Leaves ultrastructure and *in situ* expression of *MePEPC* in KU50 and W14 showed that bundle sheath (BS) cells surrounding xylem formed the Kranz-like structure in KU50 but not in W14. Green fluorescent protein GFP *in situ* hybridization shown that *MePEPC* expressed in mesophyll cell (MC) outside of the BS cells (**Fig.4 C**). Transcriptome analyses between three ancestors and 19 cultivars revealed differences in the expression pattern of a few critical genes functioning within the photosynthetic light reactions and carbon assimilation (**Fig.4 D**; **Fig.S22**). Interestingly, *MeCSK* (chloroplast receptor kinase) in photosystem II and *MeFNR3* in the electron transfer chain were up-regulated in relation to wild-type. *MeCSK* is a regulator of chloroplast genes encoded proteins of PSII as D1 and D2 responsible for light strength. In carbon fixation, only C4 genes *MePEPC, MePPDK*, and *MeNADP-Me* were significantly up-regulated in cultivars (**Fig.4D, Supplementary Table S26**). Quantitative RT-PCR confirmed that 11 genes for the light reaction, including *MeCSK, MeCytb6f, MePC, MeFNR2, MeFNR3*, and *MeFd*, were more abundantly expressed in cultivars than in wild progenitors, in particular, the *MepsbA* and *MepsbD*, the chloroplast genes encode D1 and D2 proteins consist of PSII unit didMoreover, the expression of three C4 genes (*MePEPC,MePPDK* and *MeNADP-Me*) in cultivated clones was confirmed (**Fig. 4E, Fig. S26**). Comparing the upstream 2-4 Kb promoter sequences of these photosynthesis genes across 198 cultivated clones and 28 FLA wild species (**Supplementary Table S23**) identified InDel and single-base mutations in the promoter regions of *MeCSK* (13G022900), *MeLHCB2*.*1* (01G175700), and *MeFNR*3 (02G10500). The promoter regions of *MeNADP-ME* (11G034000) associated with Carbon fixation showed varying degrees of deletion and structural variation. These structural variations affected the binding of transcriptional regulatory factors and thus regulated transcription (**Fig.4F-G, Fig. S24**). Based on these results, we hypothesize a model of light adaptation and evolution of C4 photosynthesis in cassava: The evolutionary modification in the promoters of *MeCSK* and *MeFNR3* promote regeneration of D1 and D2 proteins, stabilize PSII structure, and subsequently maintaine ATP and NADPH production; Meanwhile, the elevated expression of *MePPDK, MePEPC*, and *MeNADP-ME*, which are C4 genes with structure variations, resulted in increased biomass accumulation in cultivated cassava (**Fig.4H**).

**Fig.4.**
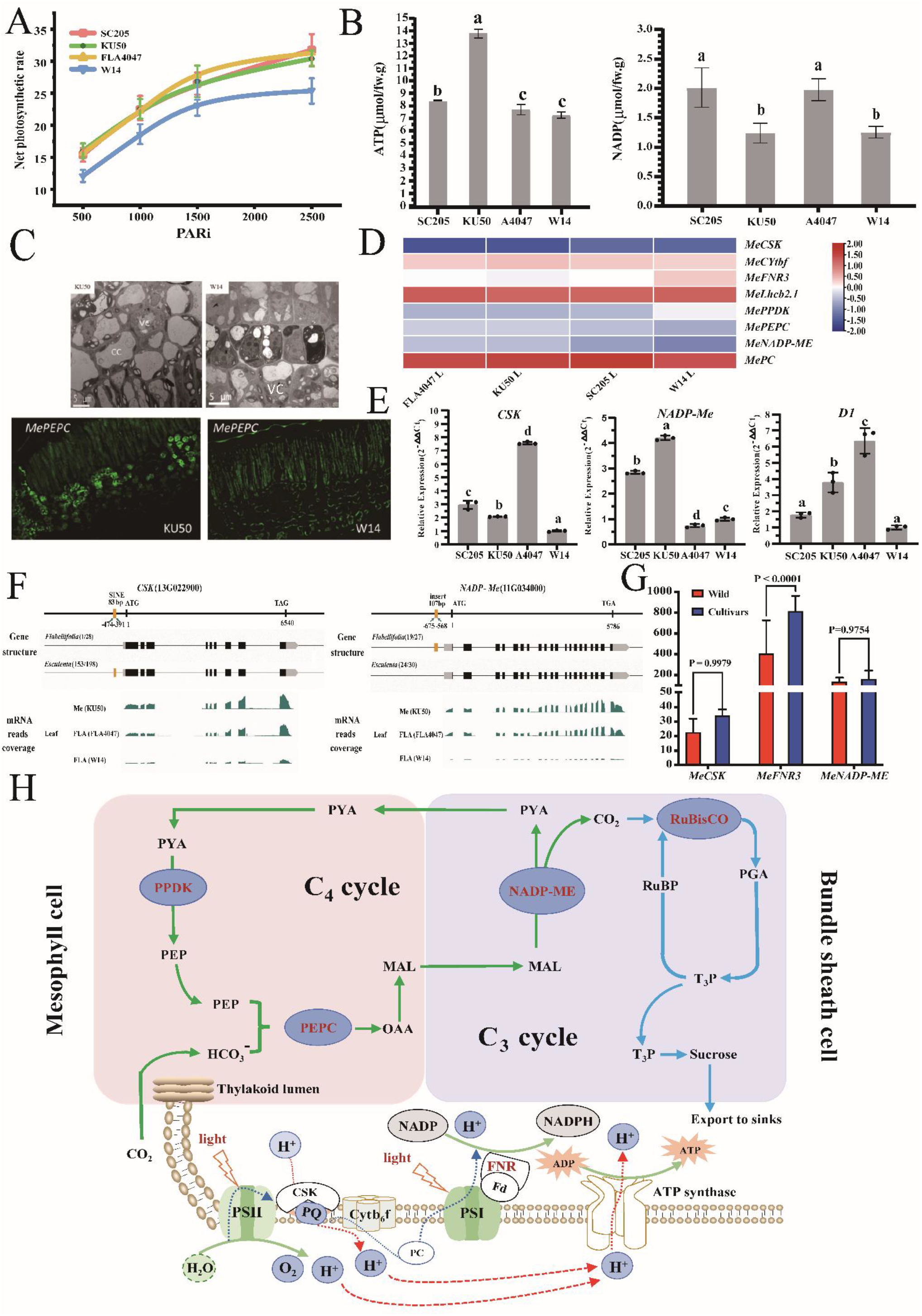
Domestication of High Light Adaptation and C_4_ Assimilation in Cassava. **A**. The Pn in functional leaf under light intensity (PARi value) from 500 to 2500 μmol/m^2^s, of cultivar KU50 much higher than wild ancestor FLA4047 and W14 with plants of DAP 120; **B**. ATP and NADPH productivity was slight higher in functional leaf of KU50 than in the FLA4047 and W14 under field condition. **C**. Leaves ultrastructure and *in situ* expression of *MePEPC* in KU50 and W14. The BS BLM:spell out or definecells surrounding xylem formed the Kranz-like structure in KU50 but not in W14; Green fluorescent protein GFP *in situ* hybridization shown that *MePEPC* expressed in mesophyll cell outside of the BS cells. **D**. Hotmap of expressive profiling of 8 genes important in light response and carbon assimilation in functional leaf 22 cultivated and wild clones: Genes in photoreaction *MeCSK, MeCytb6f, MeFNR2* and *MeFNR3* of cultivars generally increased, and C4 carbon fixation gene *MePPDK, MePEPC* and *MeNADP-ME* of cultivars up-regulated than the wild.BLM: the color differences on the panels are not clear **E**. Quantitative RT-PCR of *MeCSK, MepsbA* coding D1 protein and *MePPDK-ME* confirmed the significantly high expression in cultivar KU50, SC205 than in the wild FLA4047 and W14. **F**. Upstream structural evolution and expression differences of *MeCSK* and *MeNADP-ME* promoters between cultivars and FLA. **G**. Significant higher expression of *MeCSK* and *MeNADP-ME* in cultivated varieties(n=19)than wild accessions(n=9)counted by RNAseq. **H**. The hypothesis model of highlight adaptation and evolution of C4 photosynthesis in cassava: The evolutionary modification in promoter of *MeCSK* and *MeFNR3* promotes regeneration of D1 and D2 protein and stabilizes PSII structure then guaranteed ATP and NADPH yielding; Meanwhile, increasing expression of *MePPDK, MePEPC* and *MeNADP-ME*, the only C4 genes for somehow structure variations resulted enlarged biomass accumulation in cultivated cassava than their wild ancestors.

### Coevolution of Flowering and Storage Root Development

Storage root formation in cultivated cassava is an adaptive event important for starch yield production and is accompanied by delayed flowering. However, it is unknown if there exists a synergistic relationship between flowering time and storage root development. Wild ancestors FLA4047 and W14 start flowering around 35-40 days after seedlings emerge and produce many flowers and fruits without swollen roots. Most cultivars, such as KU50 and SC205, starts to produce storage roots 35-40 days after rooting from stem cuttings (DAP50-55), and flower around DAP300 or later, with little or no fruit and seed set (**Fig.S1**). We compared the endogenous hormone levels in floral primordial (FP) tissue of FLA4047 and W14 plants with that in stem apical meristem (SAM) and initial storage roots of KU50 and SC205 plants. The IAA, JA, ABA, and cytokinin trans-zeatin in FP of FLA4047 and W14 were 2-4 folds higher than that in SAM of KU50 and SC205 (**Fig. 5A**). Conversely, the IAA in developing storage roots of KU50 and SC205 was 2-6 folds greater than that in FLA4047. The JA, ABA, and trans-zeatin were 2-4 folds higher in cultivars than in the wild accessions (**Fig.5B**). Interestingly, the four hormones were up-regulated both in primordial tissues and initial storage roots.

**Fig.5.**
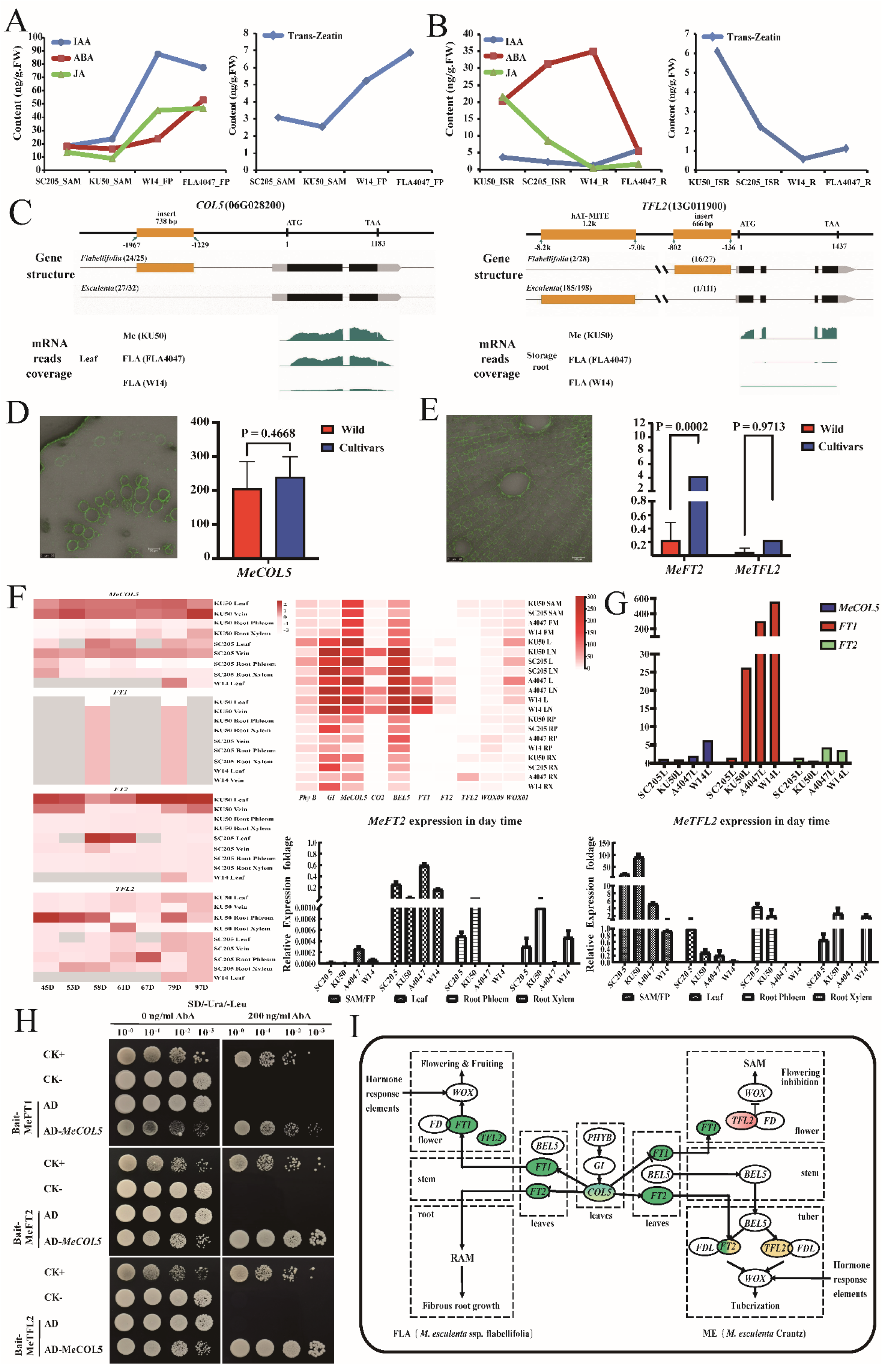
Coevolution of Flowering and Storage Root Differentiation. **A** The fold differences in the phytohormonal content of IAA, ABA, JA and Trans-zeatin in the floral primordial (FP) tissue of FLA4047 and W14 in comparison to their content in the stem apical meristem (SAM) of SC205 and KU50. **B** IAA, JA, ABA and Trans-zeatin significantly increased in the initial storage roots of cultivars SC205 and KU50 but not in the wild ancestors FLA4047 and W14. **C** Structural Variations (SVs) in the upstream sequence of *MeCOL5* and *MeTFL2* coincided with expression difference in storage roots of cassava. **D** *MeCOL5 in situ* expression in vascular bundle of SR, and it higher expressed in cultivars (n=19)than wild accessions(n=9)in statistics. **E** *MeTFL2 in situ* expression in parenchyma cells of SR, and its significantly higher expression in SC205 and KU50 than in FLAs. **F** The different expression profiling of *MeFT1, MeFT2*(SP6A), *MeCOL5* and *MeTFL2* in leaves and storage root of cultivars and wild ancestors in the initial process of FP and SR from DAP 45 to 97 (left); RNAseq data at the DAP60, *MeFT1, MeFT2* was preferentially expressed in leaves of wild FLA4047 and W14, but no expressed in FP, xylem and phloem of storage roots (right). **G** Real time**-**PCR validated that *MeCOL5, MeFT1* and *MeFT2* is predominantly expressed in leaves of ancestor FLAs with flower, and *MeFT2, MeTFL2* was preferentially expressed in storage root of cultivars. **H** Y1H assays identified that MeCOL5 interacted with the promoter of MeFT1, MeFT2 and MeTFL2. The transformed yeast cells were plated on SD/-U-L medium with 0, and 200 ng ml^−1^ AbA. CK+ representative positive control pAbAi-p53+pGADT7-p53; CK-representative negative control pAbAi+pGADT7. **I** The hypothesis model of coevolution flowering and storage root differentiation.

We compared the sequence variations in the upstream 2-4 kb promoter and gene body of 43 genes (families) involved in flowering and storage root development in wild and cultivated cassava. Noticeable structural variations are detected in the promoter regions of photoreceptor *MePHYB* (11G052800), cell cycle regulator *MeGI* (05G043800), florigen and its related gene *MeFT2* (SP6A, 13G000800), *Me14-3-3* (02G146400), *MeEFL* (04G083700) and *MeMADS* (05G187800). A 738 bp deletion occurred in the promoter of *MeCOL5* (06G028200) in most (27/32) cultivars and landraces but not in FLA accessions (24/25)(**Fig.5C**). Transposon *hAT*-MITE was inserted in the promoter of *MeTFL2* (13G011900) in 185 of the 198 cultivars, and a 666bp deletion occurred in almost cultivars (110/111) (**Fig.5C, Supplementary Table S23**). *MeCOL5* and *MeTFL2* expressed more abundantly in cultivars than in wild accessions, based on RNA-seq and *in situ* gene profiling on parenchyma cells of the primary root microtubule (**Fig.5D-E**). RNA-seq for plants at different developmental stages showed that *MeCOL5* and *MeFT2* were preferentially expressed in leaves and veins but expressed with decreased abundance in storage roots of cultivars SC205 and KU50. These two genes were expressed earlier at DAP45 in KU50 and later at DAP58 in SC205, coinciding with its storage root (SR) initiation. The florigen gene *MeFT1* (12G001600) had decreased expression in all plants we profiled. However, *MeTFL2* exhibited a higher expression in storage roots but had no detectable expression in the leaves of cultivars. Concretely, *MeTFL2* had an increased expression in SR xylem at DAP61 of KU50 and phloem at DAP67 of SC205 (**Fig.5F left**). RNA-seq on the flower primordia and storage root at DAP60 further demonstrated that *MeCOL5* preferentially expressed in leaves of cultivars and wild accessions associated with *GI, CO2* (15G148400), and *BEL5* (09G045600). *MeFT1* was preferentially expressed in leaves of wild species FLA4047 and W14 but not cultivars, while *MeFT2* was synchronously expressed at a low level in the leaves of wild and cultivated varieties (**Fig.5F right**). Real-time Q-PCR confirmed that *MeCOL5, MeFT1*, and *MeFT2* were more significantly expressed in leaves of FLA4047 and W14 than in cultivar KU50 and SC205 at DAP60, the flowering/storage root formation stage. *MeFT2* and *MeTFL2* in initiating storage roots of cultivars were significantly higher expressed than wild species (**Fig.5G**). *MeFT2* and *MeTFL2* might be important signal molecules and regulatory genes during storage root formation. Yeast single hybridization experiments illustrated that *MeCOL5* interacted with *MeFT2, MeFT1*, and *MeTFL2* (**Fig.5H**). Accordingly, we propose a model for the coevolution of flowering time and initial storage root formation in cassava. The structural variation of *MeCOL5* selectively regulates the expression of *MeFT1*/*MeFT2*, which is preferentially expressed in wild ancestors, initiating floral primordia differentiation and reproductive development. *MeFT2* was predominantly expressed in leaves of cultivars and transported to cambium cells binding with *MeTFL2* and other functional proteins to initiate storage root development (**Fig.5I**). The detailed regulatory mechanism needs additional experimental work to be understood and made ready for practical uses in cassava breeding.

### Evolution of Cyanogenic Glycoside Metabolism in Cassava

Two cyanogenic glucosides, linamarin and lotaustralin, are known to be biosynthesized and accumulate in leaves, storage roots, and other organs of cassava and to act as protectants against herbivores and pests (Tattersall *et al*., 2001; Gleadow and Møller, 2014; Ogbonna *et al*., 2021). Sweet cassava with low hydrogen cyanide potential is available for human and livestock consumption, but how this domestication has been achieved is not known. We used the genome sequences of wild and cultivated cassava to reconstruct the selection history embedded in the promoters of critical genes in cyanogenic glucoside biosynthesis, transport and remobilization metabolism. Our results indicate that altered transcriptional regulation, with the basic helix-loop-helix transcription factor *bHLH2* acting as the essential regulon, drove the domestication of cultivated sweet cassava. Evaluation of cyanogenic glucoside content in storage roots of 30 cultivars and wild sub-species analyzed at DAP120 showed an evident diversity (**Supplementary Table S24**). We compared the hydrogen cyanide potential in storage roots and leaves of wild ancestors W14 and FLA4047 with six cultivars. In storage roots, the hydrogen cyanide potential ranged from 70.89 to 397.89 μg/g, 1.4 to 5.6 folds higher in the wild species than in the cultivars. The hydrogen cyanide potential in leaves ranged from 379 to 1326 μg/g, 1.51 to 3.49 folds higher in the wild than in the cultivars (**Fig. 6A**). This result indicates that a reduced content of cyanogenic glucosides has been a selected target in the course of modern cultivated cassavas from a wild ancestor. RNA-seq data revealed that genes for hydrogen cyanide detoxification (including *MeHNL10-like, MeHNL24-like*, and *MeB-CASa*) and cyanogenic glucoside transport (including *MeGTR, MeCGTR1, MeMATE1*, and *MeH*^*+*^*-ATPase*) are down-regulated in the storage roots of sweet cultivars, whereas *MeHNL24-like, MeMATE1*, and the genes *MeCYP79D2 and MeUGT85K5* involved in cyanogenic glucoside biosynthesis are down regulated in the leaves of cultivars (**Supplementary Table S25**). The specific expression in roots and leaves of these genes is highlighted: *MeMATE1* is only expressed in storage roots and *MeGTR* mainly in leaves(**Fig. 6B**). Q-PCR further confirmed that *MeCYP79D1, MeCYP79D2, MeMATE1, MeGTR*, and *MeHNL10-like* are expressed at significantly lower levels in the pericarpium, stele of storage roots and leaves of sweet cassava KU50, SC16 and SC205 than that of FLA4047 and W14 (**Fig.6C,Fig. S28**).

**Fig.6.**
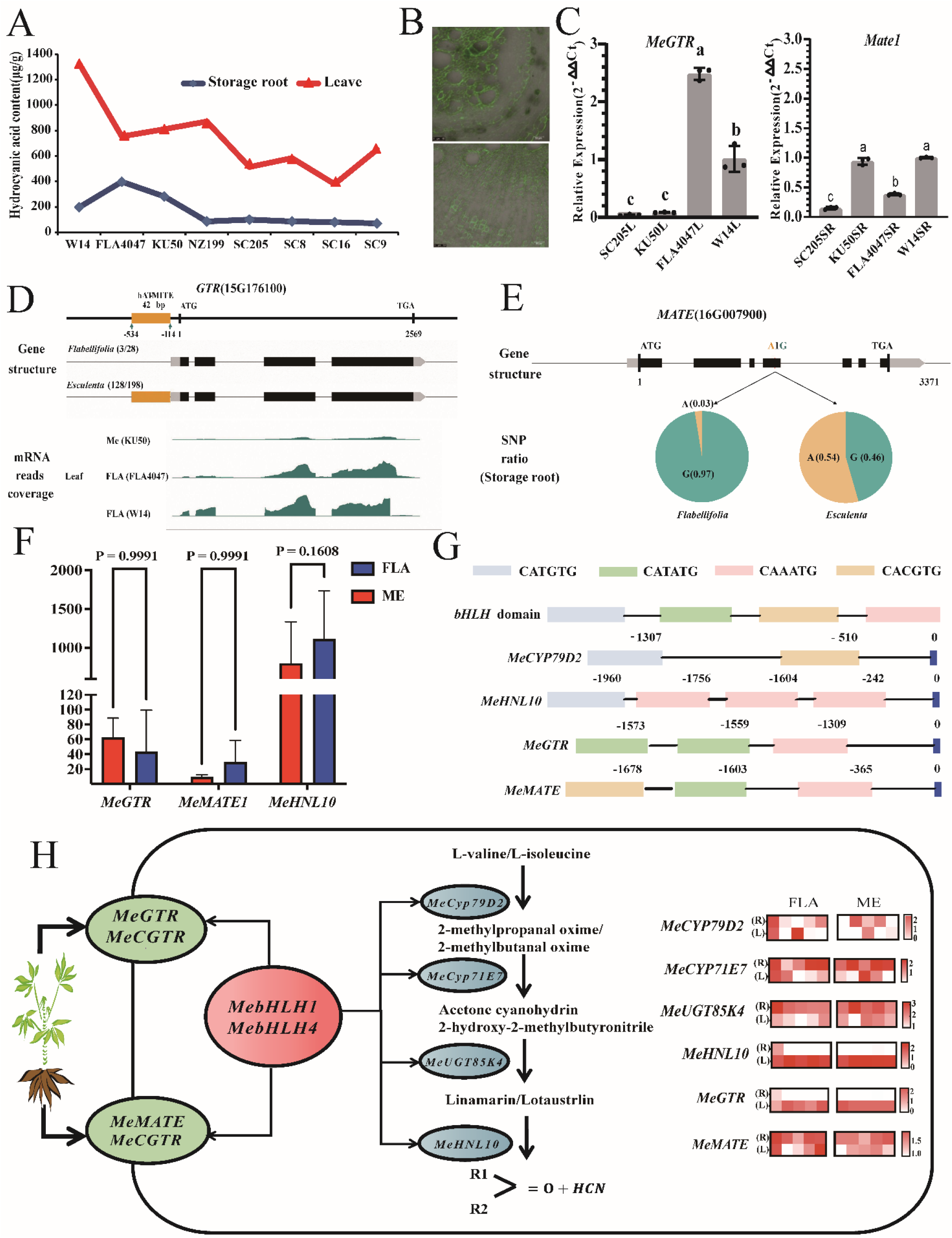
Domestication of Genes for Synthesis, Transport and Turn-over of Cyanogenic Glycosides in Cassava. **A**. Cyanogenic glucoside content in storage root and leaves of wild ancestors FLA4047 and W14 are much higher than in 5 cultivars except for KU50. **B**. *In situ* RT-PCR showed that *MeGTR* is preferentially expressed in mesophyll cells around vascular bundle of leaves, while *MeMATE1* is preferentially expressed in the secondary xylem of storage roots. **C**. qRT-PCR verified that expression of *MeGTR* in leaves (L) and *MeMATE1* in storage roots (R) of cultivars KU50 and SC205 are lower than in wild FLA4047 and W14. D. A 420 bp insertion in the promoter of *MeGTR* for most cultivars (128/298) but not in the wild accessions (3/28) may be the reason for the transcript decrease in the cultivars. **E**. A single nucleotide substitution in the *cds* of *MeMATE1* causing a G→ S substitution is present in more than half (54%) of the cultivars and rare (3%) in in the wild accessions and may reduce transport efficiency. **F**. The RNA-Seq reads of *MeGTR* is lower in leaves of cultivars compared with the wild accessions but the difference is not significant. The transcript level of *MeHNL10* is much lower in cultivars compared with the wild accessions; *MeMATE1* is significantly lower in storage roots of cultivars than in the wild accessions. P values are calculated with two-sided Student’s t-test. Numbers of accessions: n = 3 (wild accessions), n = 12 (cultivars). **G**. More than one binding domain of *bHLHs* are found in the promoter regions of *MeGTR, MeMATE, MeCYP79D2* and *MeHNL10*; **H**. DAP-Seq experiment confirmed the interaction between *MeCYP79D2* and *bHLH2* protein. **I**. Based on this set of transcriptional data, we hypothesize that the molecular evolution of sweet cassava with a low content of cyanogenic glucosides has been achieved by lower expression of genes for biosynthesis, transport and bioactivation of cyanogenic glucosides at lower levels in cassava cultivars in comparison to wild casssava. Predominant structural variations in the promoter of *MeGTR* and in *cds* of *MeMATE1* are the likely cause of the decreased content of cyanogenic glucosides in leaves and storage roots of the modern cultivars. The *MebHLH2* transcriptional factor is a key regulator of biosynthesis, transport and bioactivation of cyanogenic glucosides in cassava.

Comparison of the sequences of 12 genes with a differential expression between the wild and cultivar populations showed that only *MeMATE1, MeCGTR1*, and *MeCYP71E7* harbored nucleotide mutations causing a single to three amino acid changes in the encoded proteins. However, significant structural variations are apparent in the promoter regions of *MeGTR* and *MeMATE1*. These include a 420 bp insertion in the promoter of *MeGTR* for most cultivars (128/298) but not in the wild accessions (3/28) may be the reason for the transcript decrease in the cultivars (**Fig.6D**). A single nucleotide substitution in the *cds* of *MeMATE1* causing a G→ S substitution is present in more than half (54%) of the cultivars and rare (3%) in the wild accessions and may reduce transport efficiency(**Fig. 6E, Supplementary Table S23**). RNA-Seq data show that *MeGTR* and *MeHNL10* show reduced expression in leaves of cultivars than in wild accessions. The expression of *MeMATE1* is significantly lower in storage roots of cultivars than in the wild accessions (**Fig.6F**). The structural evolution could change the *cis* elements and modify the binding efficacy of transcription factors, resulting in significant alterations in the gene expression patterns. Analysis of *cis* elements in the promoters of the genes involved in cyanogenic glucoside biosynthesis and detoxification identified many motifs, including CATGTG, CAAATG, and CACGTG, corresponding to the binding domains of *bHLH* transcription factors (**Fig.6G**). Based on the data obtained, we present an evolution model for cyanogenic glucoside biosynthesis and detoxification in cassava (**Fig.6I**). Cassava cyanogenic glucosides were preferably synthesized in phloem parenchyma cells in leaves and storage roots, transported into phloem, and accumulated or remobilized in younger tissues. The genes for cyanogenic glucoside biosynthesis, transport, and remobilization in edible sweet cassava have been selectively modified, particularly *MeMATE1* and *MeGTR*, resulting in the decrease of cyanogenic glycoside content in leaves and storage roots. Transcription factor *bHLH2* is shown to be an integrated upstream regulator in cyanogenic glucoside metabolism.

## DISCUSSION

### Evolution and Domestication adaptive in Cassava

For the first time, we released three nearly complete reference genomes of cultivar and ancestor of cassava a tropical model crop (Wang et al., 2014; Bredeson et al., 2015; Hu et al., 2021; Qi et al., 2022). Based on pan-genome of 24 representative accessions and the re-sequencing data of 484 diverse collections, we produced an integrated Pan SV Cassava map covering 346,322 SVs, 96,032,008 SNVs distributed in 18 pairs of chromosomes, that accounted for 13.4% of the genome. The genetic variation was significantly higher than Bamboo (Zhao et al, 2021), Lychee (Hu et al, 2022) and Grape (Magris et al, 2021). The evolutionary tree was reconstructed with 484 cassava accessions which confirmed the FLA is the original species and the domestication pathways of cultivated cassava (Olsen et al., 1999). According to the evolutionary analysis, the cultivated cassava differentiated from FLA at about 0.02MYA years, while the progenitor subspecies FLA diverged from the outgroup GLA was about 0.18MYA. The estimation is consistent with the local agriculture origin explored with carbon isotope analysis in general. With large-scale forest island samples in highland nearby, it was convinced that Manioc (*Manihot* sp.) found at about 10,350 years ago, and Quash (*Cucurbita* sp.) at about 10,250 years (Lombardo et al., 2020). Relevant data link to China’s national data center of tropical crop genome database (https://ngdc.cncb.ac.cn/tcod/home).

Parallel evolution of C_4_ enzymes and photochemical reactions enhanced photosynthetic productivity in cassava. ^14^C isotope experiments already proved that cultivated cassava has a C_4_ enzyme system (Cock et al., 1987). We further found that the leaves of cassava had a similar Kranz anatomy through electron microscopy and *in situ* hybridization, and it could act as a CO2 concentration pump. *MeRuBisCO* predominantly expressed in vascular sheath cells and *MePEPC* was more expressed in peripheral mesophyll cells, as in C_4_ maize (Llorens et al., 2017). Different from the wild ancestor, *MePPDK, MePEPC* and *MeNADP-ME* in functional leaves of cultivars were enhanced expressed, and *cis* element mutation only found in the promoter region of *MeNADP-ME*, not in other two genes. However, systemic structural variation occurred in the promoter region of *MeCSK* and *MeFNR1* in the photochemical response pathway, and their expression amount significantly higher in cultivars than in the wild subspecies correspondingly. Chloroplast receptor kinase (*CSK*) might be a key signal switch, It has a ferrithionein structure that balances plastiquinone (PQ) electron transfer and coordinates the expression of multiple chloroplast genes of Photosystem II toward to regulation of highlight adaptation (Puthiyaveetil *et al,*2008, 2012, 2013; Iskander M. Ibrahim, 2020). SV mutation analysis revealed CNV expansion took place in 13 photosynthesis-related gene families, and *MeCSK* and genes for glucose metabolism were highly selected. The latest review of C_4_ photosynthesis suggests new ideas for C_4_ plant improvement, such as increasing the proportion and the size of BS cells and the number of chloroplasts in BS cells (Cui H, 2021). Cultivated cassava owned C_4_ enzyme system and kranz anatomy could be used as a typical material for C_4_ improvement. It is extremely valuable in theory to decipher the high light adaptation and heavy biomass accumulation utilizing cassava with C_3_-C_4_ intermediate photosynthesis.

Structural variation of *MeCOL5* and *MeTFL2* essentially driving storage root formation in cassava. Storage root formation is the most profound change from wild ancestor species to cassava varieties. Along with flowering delay or degradation of reproductive growth, more rely on vegetative stems than seeds. In biology, the center of growth and photosynthates accumulation changed from seeds to underground storage roots. By comparing the *cds* and promoter sequences of 54 genes involved in FP and storage root development between FLA and cultivars, we found that only *MeCOL5* and *MeTFL2* had systematic structure evolution, and *MeCOL5* was induced expression in leaves under appropriate light length, temperature signal and inner sugar level. Yeast one hybrid demonstrated that *MeCOL5* selectively interacted with the florigen *MeFT1* and *MeFT2 (MeSP6A)*, produced isogenous mobile mRNA and signal protein in potato (Navarro et al, 2011). *MeCOL5* preferentially initiated expression in cultivar leaves of *MeFT2*, whom was transported into roots as a signal molecule, combined with *MeTFL2* to initiate storage root development. In wild ancestral species, *MeCOL5* promoted expression of *MeFT1* and regulated floral primordium development. Interestingly, the hormone signals after storage root or flower primordium initiated are remarkably similar. (Kuznetsova et al., 2020; Hoang et al, 2020). *CONSTANS (CO)* and its homologous genes regulate flowering in Arabidopsis, rice and potato (Mariana et al.,2006; Gonzalez-schain et al., 2012), *FT1* and *FT2* are synthesized in leaves and transported to floral primordia over long distances (Laurent et al., 2007; Daniela et al., 2020) and potato tuber (Navarro et al.,2011) have been widely reported. Both *FT1* and *FT2* were regulated by several upstream binding proteins and transcription factors, as photoreceptors, *CONSTANS* and *FD* (Zhu et al., 2020) and linked to sugar leakage (Sweet11), activation of auxin and CK (POTH, BEL5) (Navarro et al., 2011; Hoang et al., 2020). It has been proved that CO-FT system integrates exogenous light and temperature signals as well as endogenous sugar and hormone signals to become the center of flowering regulation (Freytes et al, 2021). In addition, potato root tuber development is inhibited by high temperature (Lehretz et al., 2019), while cassava is obviously different. If and how was it transported of *MeSP6A*, and what binding proteins in storage root development of cassava are still needed to be further uncovered. It is reported that *TFL2* is usually recruited by the bZIP transcription factor FD to regulate flower organ differentiation and cell growth (Silvio *et al*.,1999; Daniela *et al*., 2020).

### Understanding cyanide evolution in cassava

Presently, the synthesis, transport and catabolic pathways of cyanogenic glucosides and involved genes have been clarified in cassava. But how they are regulated by environmental factors and endogenous sugar and hormone signals remains unknown. In a structural comparison of all 22 genes for cyanide metabolism between clones with high cyanide include of wild ancestors and varieties with low cyanide, it was found that the promoter region of *MeGTR*, a cyanide transporter had been selected, which resulted in down-regulated expression in leaves specifically. The SNP mutant altered an amino acid of *MeMATE1* and caused down regulation of its cyanide transport capacity that will probably be one of typical events in domestication of sweet cassava. This is consistent with that a single nucleotide variation in *MATE* alter cyanoside content of cassava (Ogbonna et al., 2021). In addition, *MeCYP79D1*, key to synthesis and *MeHNL10-like* response to decomposition cyanide were significantly down-regulated in both leaves and storage roots of low cyanide cultivars, but no systematic variation was observed in gene body sequences. However, we found *bHLH* binding motifs in the promoter regions of major cyanoside metabolism genes. Yeast one hybrid experiment confirmed *MebHLH2* interacted with *MeCYP79D2*, that indicated *MebHLH2* could participate the metabolic regulation of cyanide synthesis, transport and decomposition. This seems to be parallel evolution with the bHLH transcription factor minimizing cyanide in sweet almond (Sanchez-Perez et al., 2019). These findings provide an important basis for gene editing to improve sweet varieties of cassava and other crops (McMahon et al., 2022).

## Supporting information

STAR+METHODS

## STAR ★ METHODS

## ACKNOWLEDGMENTS

We thank Hongbin Zhang for academic and language revision; Robert Henry,Peter Kulakow, Cankui Zhang, Zhangjun Fei, Peng Zhang and Weimin Tian for comments and suggestions for the article; Yimin Bao for data deposition. Funding: The research has been supported by grants from the National Key R&D Program of China (2018YFD1000501), the National Natural Science Fundation of China-CG joint fundation (3181101517) and The startup funds for the double first-class disciplines of crop science in Hainan University (RZ2100003362).

## AUTHOR CONTRIBUTIONS

Z.X., Z.D. and X. Z. chiefed the genome sequencing, all data analysis and jointly conceived the study. W.W., M. L. and J.X. designed the research and wrote the article. S. J., T. Z., L. W., M. C., F. C., H. Z, W. Z., S. H. and Y. W analyzed the data. L. C., L. A. B. H. C., H. Y. and X. Z. donor part of genotypes and transcriptome data. M. Z., L. X., Z. W., B. F., S. W, M. L., Y. L, H. W, S. L, Y. B., L. Z, C. Z., J. X, F. G, X. S., C. L and F. Q performed the experiments organized by X. C. Y. C, D. H, K. L, B. L. and M. P. supervised the experiments. W. Z., H. Z. and B. M. contributed to refined the key points and language in English. All authors discussed the results and took part in writing the manuscript.

