## Supplementary material for "Pan-genome and Haplotype Map of Cultivars and Their Wild Ancestors Provides Insights into Selective Evolution of Cassava (*Manihot esculenta* Crantz)": STAR+METHODS

**KEY RESOURCES TABLE**

| REAGENT or RESOURCE | | SOURCE | IDENTIFIER |
| --- | --- | --- | --- |
| Deposited data | | | |
| *M.esculenta* Crantz AM560 genome assembly | This study | | GWHBHDV00000000 |
| *M.esculenta* SSP*. flabellifolia* FLA4047 genome assembly | This study | | GWHBHDU00000000 |
| *M.esculenta* SSP*. flabellifolia* W14 genome assembly | This study | | GWHBHCP00000000 |
| Pan genome assemblies for 21 selective genotypes: | This study | | Following: |
| Manes.BX genomes | This study | | GWHBHCQ00000000 |
| Manes.S25 genomes | This study | | GWHBHCR00000000 |
| Manes.GLZyn genomes | This study | | GWHBHCS00000000 |
| Manes.TY genomes | This study | | GWHBHCT00000000 |
| Manes.NZ genomes | This study | | GWHBHDC00000000 |
| Manes.IT4 genomes | This study | | GWHBHDD00000000 |
| Manes.GR4 genomes | This study | | GWHBHDE00000000 |
| Manes.MB genomes | This study | | GWHBHDF00000000 |
| Manes.SC9 genomes | This study | | GWHBHDG00000000 |
| Manes.IB5 genomes | This study | | GWHBHDH00000000 |
| Manes.BC6 genomes | This study | | GWHBHDI00000000 |
| Manes.BC8 genomes | This study | | GWHBHDJ00000000 |
| Manes.ZM genomes | This study | | GWHBHDK00000000 |
| Manes.IB3 genomes | This study | | GWHBHDL00000000 |
| Manes.YNZ genomes | This study | | GWHBHDM00000000 |
| Manes.WCH genomes | This study | | GWHBHDN00000000 |
| Manes.SC8 genomes | This study | | GWHBHDO00000000 |
| Manes.S68 genomes | This study | | GWHBHDP00000000 |
| Manes.SC5 genomes | This study | | GWHBHDQ00000000 |
| Manes.FX genomes | This study | | GWHBHDR00000000 |
| Manes.HB genomes | This study | | GWHBHDS00000000 |
| 30X Resequencing data for 290 accessions of MEs, FLAs and GLAs | This study | |  |
| RNAseq data for 118 plant organs of cassava and its wild relatives | This study | | CRA008070 (https://ngdc.cncb.ac.cn/gsub/submit/gsa/subCRA012245) |
| Other 196 accessions with re-sequencing data down loaded | Punna Ramu *et al*, 2017 | | https://doi.org/10.1038/ng.3845 |
| AM560 JGIv7.1 genome assembly | Bredeson et al., 2016 | | https://phytozome-next.jgi.doe.gov/Mesculenta |
| Supplemental Information and Supplemental Table supporting the manuscript | This study | |  |
| Biological Samples |  | |  |
| *Manihot esculenta* Crantz AM560 | This study | | CIAT |
| *Manihot esculenta* SSP. *flabellifolia* FLA4047 | This study | | Germplasm and Biotechnology, EMBRAPA, Brazil |
| *Manihot esculenta* SSP. *flabellifolia* W14 | This study | | https://DOI: [10.1038/ncomms6110](https://doi.org/10.1038/ncomms6110) |
| Other 290 accessions include of FLA and ME used for Pan genome and re-sequencing | This study | | Supplementary Table: S Table1 |
| The 118 plant samples used for RNA-seq data collection | This study | | Supplemental information Table S9 |
| Experimental Models: Organisms/Strains |  | |  |
| Software and Algorithms |  | |  |
| Sniffles | Sedlazeck et al., 2018a | | https://github.com/fritzsedlazeck/Sniffles |
| BUSCO | Sima˜ o et al., 2015 | | https://busco.ezlab.org/ |
| DiffMatrix | Hyunjoo Song et al.,2012 | | https://github.com/mbrazeau/diffmatrix |
| NextDenovo | https://github.com/Nextomics/NextDenovo | | https://github.com/Nextomics/NextDenovo |
| Smartdenovo | Liu H et al.,2021 | | https://github.com/ruanjue/smartdenovo |
| BWA v0.7.12 | Li H, 2013 | | https://github.com/lh3/bwa |
| SAMtools | Li, Heng et al., 2013 | | http://samtools.sourceforge.net |
| VCFtools | Danecek, Petr et al., 2011 | | https://vcftools.github.io/license.html |
| GCTA | Yang, Jian et al., 2011 | | http://gump.qimr.edu.au/gcta |
| PopLDdecay | Zhang, Chi et al.,2019 | | https://github.com/BGI-shenzhen/PopLDdecay |
| BLAST | Altschul, S F et al., 1990 | | https://blast.ncbi.nlm.nih.gov/Blast.cgi |
| Bionano Solve v3.3 | https://bionanogenomics.com/support-page/data-analysis-documentation/ | | https://bionanogenomics.com/support/software-downloads/ |
| Stitch.pl | Shelton et al., 2015 | | https://github.com/i5K-KINBRE-script-share/Irys-scaffolding/tree/master/KSU_bioinfo_lab/stitch |
| RepeatMasker with Repbase | Bao et al., 2015 | | http://repeatmasker.org |
| RepeatProteinMask | Bao et al., 2015 | | https://github.com/rmhubley/RepeatMasker/blob/master/RepeatProteinMask |
| RepeatModeler2 | Flynn et al., 2020 | | http://www.repeatmasker.org/RepeatModeler/ |
| Exonerate | Slater, Guy St C, and Ewan Birney, 2005 | | https://www.animalgenome.org/bioinfo/resources/manuals/exonerate/index.html |
| genBlastA | She, Rong et al. , 2009 | | <http://genome.sfu.ca/projects/genBlastA/> |
| Augustus | Stanke, Mario et al., 2008 | | http://augustus.gobics.de/ |
| GlimmerHMM | Majoros, W H et al., 2004 | | http://ccb.jhu.edu/software/glimmerhmm/ |
| SNAP | Korf, Ian, 2004 | | https://github.com/KorfLab/SNAP |
| Genemark.HMM | L. S., et al., 2005 | | http://exon.gatech.edu/GeneMark |
| IsoSeq | Di Tommaso, Paolo et al., 2017 | | https://github.com/nf-core/isoseq/ |
| LoRDEC | Salmela, Leena, and Eric Rivals.,2014 | | https://www.lirmm.fr/~rivals/lordec/ |
| BLAT+CD-hit-est | Kent WJ, 2002 | | http://genome.ucsc.edu |
| Infernal | E. P. Nawrocki and S. R. Eddy, 2013 | | http://eddylab.org/infernal/ |
| tRNAscan-SE | Lowe TM, Eddy SR, 1997 | | https://www.psc.edu/resources/software/trnascan-se/ |
| RNAmmer | Lagesen, Karin et al. , 2007 | | http://cbs.dtu.dk/services/RNAmmer/ |
| EVM | Haas et al., 2008 | | https://evidencemodeler.github.io/ |
| PASA | Haas et al. ,2008 | | https://github.com/PASApipeline/ |
| Blast2GO | Conesa, Ana et al. , 2005 | | https://www.blast2go.com/ |
| KEGG | Kanehisa, M, and S Goto. ,2000 | | http://www. genome.ad.jp/kegg/ |
| nr/nt | <ftp://ftp.ncbi.nlm.nih.gov/blast/db/FASTA> | | <ftp://ftp.ncbi.nlm.nih.gov/blast/db/FASTA> |
| InterPro | Mulder, Nicola, and Rolf Apweiler., 2007 | | https://interpro-documentation.readthedocs.io/en/latest/citing.html |
| FALCON | Chin, Chen-Shan et al. , 2016 | | https://pb-falcon.readthedocs.io/en/latest/ |

**RESOURCE AVAILABILITY**

**Lead Contact**

Further information and requests for resources and reagents should be directed to Wenquan Wang.

**Materials Availability**

This study did not generate new unique reagents.

**Data and code availability**

All sequencing data generated in this study have been deposited at the Genome Sequence Archive (GSA, <https://ngdc.cncb.ac.cn/gsa> ) under BioProject PRJNA557253. Github repositories for software presented in this work are listed as follows: https://github.com/malonge/DupCheck, https://github.com/mkirsche/Jasmine, https://github.com/srividya22/geneLift, https://github.com/malonge/CallIntrogressions.

**EXPERIMENTAL MODEL AND SUBJECT DETAILS**

Plants of 290 accessions in *Manihot esculenta* Crantz, its wild ancestor subspecies *Manihot esculenta* SSP. *flabellifolia* and relative species *Manihot glaziovii* growing in Chengmai county , Hainan island, Southern China. We sampled leave and storage roots for DNA sequencing and RNA-seq where there are excellent sunlight and rainfall conditions available for cassava growth and development.

**METHOD DETAILS**

**Genome size measurement by flow cytometry**

The genome size of 1C=0.72 ±0.02 Gb for the cultivated cassava, *M. esculenta* with accession KU50，the same as its nearest ancestor FLA, *M.esculenta* SSP. *flabellifolia* with accession FLA4047. While the genome size of relative species GLA, *M. glaziovii* with accessions GLAyn and GLAb is 1C=0.73±0.01 Gb, Differently, the W14, originally ascribed into *M. esculenta* SSP. *flabellifolia* with genome size 1C=0.83±0.01. This estimated results slightly larger than the previously reported (**Wang et al., 2014**). Thus, genome size estimations by flow cytometry were performed according to Dolezel et al. (2007). For that, roughly 0.5 cm^2^ of young leaf tissue was chopped with a sharp razorblade in a Petri dish together with appropriate amounts of leaf tissue of the internal reference standard rice with accession [Nipponbare](javascript:;)（1C=0.43Gb）and maize with accession B73（1C=2.30Gb）. Using the ‘CyStain PI Absolute P’ nuclei extraction and staining kit (Sysmex-Partec). The resulting nuclei suspension was filtered through a 50-mm filter (CellTrics,) and measured on a CyFlow Space flow cytometer (FAC Scalibur). The absolute DNA content (pg/2C) was calculated based on the values of the G1 peak means and the corresponding genome size (Mb/1C).

**Library preparations and sequencing**

***DNA isolation***

High-molecular-weight DNA was isolated from 1.5 g of material with a QIAGEN® Genomic DNA Extraction Kit (Cat#13323 Qiagen). Quality was assessed with a NanoDrop™ One UV-Vis spectrophotometer (Thermo Fisher Scientific, USA), and quantity was measured accurately with a Qubit® 3.0 Fluorometer (Invitrogen, USA).

***Whole-genome shotgun sequencing (WGS)***

Genomic DNA from W14, FLA4047 and AM560 were deep-sequenced with an Illumina HiSeq 2500 in 150-bp paired-end mode. Also, DNA samples from 290 accessions were sequenced with an Illumina HiSeq 2500 reached to 30X coverage each sample.

***PacBio library construction and sequencing***

PacBio libraries for W14 with the insert size of >10 kb were pre-pared and then sequenced with P6-C4 reagent kits on PacBio RSII (Pacific Biosciences, Menlo Park, CA), and a total of 18G clean reads were obtained for W14 with sequencing depth of 25X.

***ONT library construction and sequencing***

DNA Long libraries of AM560 and FLA4047 with gDNA size-selected (>20kb) and used a LSK108 ligation Kit (Cat#SQK-LSK108, Oxford) were constructed and accurately detected by Qubit® 3.0 Fluorometer. The DNA library was added to a flow cell, which was transferred to Nanopore GridION X5 (Oxford Nanopore Technologies, UK) for real-time single molecule sequencing. The ultra-long DNA libraries for AM560 (2 cells), FLA4047 (2 cells) and W14 (2 Cells) genomes were constructed respectively used for complete genome sequences assembly. For each ultra-long Nanopore library approximately 8-10 hg of gDNA was size-selected (>50 kb) with SageHLS HMW library system (Sage Science，USA)，and processed using the Ligation sequencing 1D kit (SQK-LSK109, Oxford Nanopore Technologies，UK) according the manufacturer s instructions. Menwhile, long DNA libraries have been constructed with >50 kb for 21 accessions and assemblied independently used for pan-genome analysis. About 800ng DNA libraries were constructed and sequenced on the Promethion (Oxford Nanopore Technologies, UK) at the Genome Center of Grandomics (Wuhan, China).

***BioNano***

The ultra high-molecular-weight (uHMW) DNA was isolated from young leaves of *Manihot esculenta* genotype W14 and AM560 by the Amplicon Express (Pullman, WA, USA). The nicked DNA molecules were labeled and stained according to the instructions of the IrysPrep Reagent Kit (Bionano Genomics, San Diego, CA, USA) for NLRS method as described in detail in Luo et al. (2017). The DNA sample was loaded onto the nanochannel array of an IrysChip (Bionano Genomics, San Diego, CA, USA) and was automatically imaged by the Irys system (Bionano Genomics, San Diego, CA, USA). For DLS method, the uHMW DNA molecules were labeled with the DLE-1 enzyme (Bionano Genomics, San Diego, CA), and were stained according to the instructions of the Bionano Prep™ Direct Label and Stain (DLS) Kit (Bionano Genomics, San Diego, CA).

***RNAseq***

Total RNA was isolated from 118 different tissues as leaves, storage roots, and flower buds of cassava and its wild relatives FLAs and GLAs. Poly-A RNA was enriched from 1 mg total RNA using the NEBNext® Poly(A) mRNA Magnetic Isolation Module. The quality and quantity were further assessed using NanoDrop 2000C and Agilent 2100 platforms. For illumina sequencing, 3 mg RNA from each sample were used to cDNA library construction using the NEBNext Ultra RNA library Prep Kit. The PCR products obtained were purified (AMPure XP system) and library quality was assessed on the Agilent Bioanalyzer 2100 system. Library preparations were sequenced on illumina HiSeq 2500 or HiSeq X ten systems according to experimental designs.

**Optical map construction**

Two optical maps using NLRS (nick, label, repair, and stain) and DLS (direct label and stain) technologies were constructed for W14, and one optical map using the DLS technology was constructed for AM560.

Raw DNA molecules >20 kb were collected and converted into BNX files by AutoDetect software to obtain basic labelling and DNA length information. Those molecules >180 kb and also passing quality control were aligned, clustered, and assembled into an optical map using the Bionano Genomics assembly pipeline (Bionano Genomics, San Diego, CA, USA). The thresholds used for pairwise assembly, extension/refinement, and final refinement stages were 1 × 10^−8^, 1 × 10^−9^, and 1 × 10^−15^, respectively. A consensus optical map was *de novo* assembled using the Bionano Solve v3.3 package with significance cutoffs of P < 1 × 10^-8^ to generate draft consensus contigs, P < 1 × 10^-9^ for draft consensus contig extension, and P < 1 × 10^-15^ for final merging of the draft consensus contigs. A recipe of “haplotype”, “noES”, and “noCut” was chosen. The initial optical map constructed by using either method was then checked for potential chimeric contigs and further refined.

**Genome assembly,** correction and validation

By ONT ultra-long sequencing, a total of 138G clean reads were obtained for FLA4047 with sequencing depth of 184X, N50>42K, a total of 142G clean reads were obtained for AM560 with sequencing depth of 189X, N50>42K. A total of 127G clean reads were obtained from W14, sequencing depth was 170X, N50>42K. FLA4047 was initially assembled by NextDenovo (read_cutoff = 1k，seed_cutoff = 65K、pa_correction = 20、seed_cutfiles = 20、sort_options = -m 20g -t 10 -k 40、minimap2_options_raw = -x ava-ont -t 8) , then using the purge_ Dups to remove redundant sequences and Nextpolish software combined with next generation sequencing data for error correction, the final preliminary assembly of FLA4047 genome size was 678M, contig N50 26.8M. And the preliminarily assembled FLA4047 genome was aligned to the reference genome by using the MUMmer genome sequence alignment software. Among them, six complete contigs were aligned to six different chromosomes, and the remaining 12 chromosomes were spliced through the ragtag toolkit (among the other 12 chromosomes, chromosomes 18, 17, 12 and 11 can be aligned to two different long contigs. After the contigs are spliced, they are aligned again. The other 8 chromosomes are spliced through ragtag toolkit). Finally, FLA4047 genome was obtained with six chromosomes with no gap and four chromosomes with only one gap. In the same way, W14 through NextDenovo (read_cutoff = 1k,seed_cutoff = 73k, pa_correction = 20, seed_cutfiles = 20, sort_options = -m 20g -t 10 -k 40, minimap2_options_raw = -x av -ont -t 8) preliminary assembly as well as Nextpolish next generation error correction, mummer alignments, ragtag alignments with incomplete chromosome resplicing results in a final genome size of 713M, contig N50 32.98M, among which 12 chromosomes had No gap,5 chromosomes had only one gap, and 1 chromosome had 3 gaps. AM560 finally obtained genome size of 645M, contig N50 of 35.9M, including 16 chromosomes with No gap,1 chromosome with 2 gaps, and 1 chromosome with 1 gap.

**Repeat identification and gene annotation**

Repeat elements in the 3 cassava genomes were screened using RepeatMasker with the Repbase library (*Smit, AFA, 2013-2015*), and transposable element encoded proteins were identified using RepeatProteinMask. The identification of de novo transposable elements was performed using RepeatModeler2 (Flynn, JM., *et al*, 2020). Three approaches were combined to predict genes. AUGUSTUS (Stanke, M. *et al.*, 2004), Glimmer-HMM (Majoros, W.H, 2004), and SNAP (Korf, Ian, 2004) were used for ab initio prediction. We also assembled RNA-seq reads into tran- scripts using Trinity (Grabherr, M.G. *et al.*, 2011) for gene structure annotation. Protein sequences from *Arabidopsis thaliana*, *Populus trichocarpa*, and *Ricinus communis* were aligned to the cassava genome using Exonerate (Slater, G.S. & Birney, E, 2005) for homology-based gene prediction. The gene sets predicted by the three approaches were integrated with EVM (Haas, B.J. et al., 2008) to produce consensus gene models, which were then updated by PASA (Haas, B.J, 2003). The cassava protein sequences were aligned to the NCBI non-redundant (nr) protein database using BLAST (*E-value <=1e-5*) (v2.2.6), and the best hit of alignment was used to infer their biological function. Gene Ontology annotation was performed using Blast2GO ([Flynn](https://pubmed.ncbi.nlm.nih.gov/?term=Flynn+JM&cauthor_id=32300014) JM. *et al.*, 2020) which assigned homologous sequences aligned by BLAST with NCBI nr database to GO terms. The cassava protein sequences were also compared with KEGG GENES database by BLAST (*E-value <=1e-5*) for KO (KEGG Orthology) assignments and pathway mapping.

**Pan SV genome construction**

Using AM560-2 v7.0 assembly as the reference genome, we have counted the gene density and repetitive sequence information of 21 cultivated cassava Fig. 13A, used sniffles to detect SV, and analyzed the SV statistics and found that the number of INS and DEL is the largest, followed by TRA(Fig. 13A; Fig. 13B; Table 8). Counting 5 types of mutations, the maximum length of missing SV is greater than the maximum length of INS, followed by INV, DUP (Fig. 13C). Among 21 cultivated cassava, ZM9781 has the least variation, and GR4 has the most SV (Fig. 13D). For INS, we counted the number and size of fragments in 21 cultivated cassava. We found that the number of inserts was between 2734-7391. The minimum insert size was 674,329 bp, the maximum was 1,923,718 bp, and the average insert size. It is ~243bp (Fig. 14).

**Identification of structural variations (SVs) and copy number variations (CNVs)**

MUM&Co^[1]^ is a single bash script to detect structural variations (SVs) utilizing whole-genome alignment(WGA). Using MUMmer’s nucmer alignment, MUM&Co can detect insertions, deletions, tandem duplications, inversions and translocations greater than 50 bp. MUM&Co’s initial inputs are two genomes for comparison. The AM560 genome was set as the reference genome for comparison.

**Phylogenetic tree reconstruction**

A population of 482 cassava accessions (supplementary table) from around the world was collected for resequencing. Genomic DNA extracted from fresh leaves was used for 350bp Illumina library preparation. The sequencing protocol was the same as described above. Raw data were filtered with fastp software to obtain clean data. Paired end reads were mapped to the AM560 genome using the ‘mem’ parameters of BWA^i^ (v0.7.8). Duplicate reads were removed using SAMtools^ii^(v.0.1.19), and the data were converted to a genome analysis toolkit 97 (GATK)^iii^ - identifiable sorted file. Each joined genomic variant was then identified by the haplotypecaller module of the GATK software, resulting in a gvcf file for each individual. All gvcf files were merged to identify high-quality SNPs and indels using the haplotypecaller module of GATK, filtering the following four parameters: depth ≥ 4 for individuals, genotype quality ≥ 5 for individuals, minor allele frequency (MAF) ≥ 0.05 and missingness ratio ≤ 0.1. The identified SNPs and indels were further annotated with effSNP and classified into the following groups: intergenic, upstream, and downstream.

Population genetic structure was analyzed using the program ADMIXTUR^iv^ (V1.23) with k values (number of assumed populations) ranging from 2 to 12. The k = 5 was chosen because the error value for cross validation is minimum at this time. VCF2diffmatrix was used to calculate the genetic distance of each sample from cassava populations, and the matrix files were imported into neighbor software to convert into tree files, which were visualized using ITOL (online Sketchpad website). PCA analysis was done with GCTA^v^ and visualized in R. Nucleotide diversity (π) and fixation index (FST) were calculated by vcftools^vi^ in 10 KB steps in 50 KB windows, and the 20 replicate results were plotted using boxplot in R. To estimate and compare the LD patterns between different groups, the squared correlation coefficient (R2) between pairwise SNPs was calculated using PopLDdecay (v.3.40)^[2]^ software. Parameters in the program were MaxDist 500 -- MAF 0.05 -- Miss 0.1. Average R2 values were calculated for pairs of markers in 500 KB windows and averaged across the genome.

**Selective sweeping analysis**

**Detection of genetic variation**：Based on the assembled genome of cultivated AM560, Samtools FAdix was used to construct indexes, BWA software was used to map all the re-sequencing reads of 238 cassava clones onto the reference genome. After removing the redundancy in the database, the Haplotype Caller program in GATK software was used to detect the variation of each sample and obtain the GVCF file of each genotype. Then GenomicsDBImpor and GenotypeGVCFs programs can be used to extract SNP and InDel. Finally, VariantFiltration was used for cleaning of SNPs and InDels. Filter criteria, the SNP selection "QD < 2.0 | | MQ < 40.0 | | FS > 60.0 | | SOR > 3.0 | | MQRankSum < 12.5 | | ReadPosRankSUM < - 8.0" as the filter parameters, Filtering of InDel, choose "QD < 2.0 | | FS > 200.0 | | SOR > 10.0 | | MQRankSum < 12.5 | | ReadPosRankSUM < - 8.0" as a parameter for filtering.

**Selective sweeping analysis:** We focused on the genomic evolutionary selection signals between 198 accessions of cultivated species (*Manihot esculenta* Crantz) and 28 accessions of its wild subspecies (*Manihot esculenta* ssp. *flabellifolia*). The Reduction of Diversity index (*ROD*), population genetic differentiation index (*Fst*) and their combined analysis (*ROD-Fst*) were used for analysis. In brief, a window was set every 100 KB on the 18 chromosomes of cassava, and the corresponding index of each window was calculated with the distance of 10 KB as the forward sliding distance. The *ROD* and *Fst* values of the diversity reduction index on each window of cultivated population and wild population were calculated. These two values were selected according to the criteria of top 5%, top 2% and top 1%, respectively. In addition, ITIS software were used to screen the distribution of Hat-mite, Mutator-mite, Stowaway and SINE transposons in the promoter region (within 2000bp upstream of the coding region) of cassava genes in cultivated and wild populations. With the aim to determine whether the promoter regions of the selected genes in the two populations have different transposon distribution.

**Measure of photosynthesis parameters**

**Net photosynthetic rate (Pn):** Photosynthetic parameters including the Pn per leaf area, stomatal conductance (Gs), and transpiration rate (Tr) of cassava varieties KU50, and its wild ancestor FLA4047, W14 plants were measured under field conditions at AM 9:00 to 12:00 using an LI-6400XT portable photosynthesis system (LI-COR, Lincoln, NE, USA) according to the manufacturer’s instructions. The measurement adopted artificial light with intensity (PARi value) from 500 to 2500 μmol/m^2^s.

**ATP content:** According to the Instruction manual for ATP content detection Kit of SolarBio to prepare the Reagents 1-6, ATP standard solution was diluted with distilled water to 0.625 μmol/mL solution. 0.1g of functional leaves of cassava was added to 1mL of extract for homogenization in an ice bath, centrifuged at 8000g for 10min at 4℃, and the supernatant was put into another EP tube and 500μL of chlorine was added. The samples were thoroughly shaken and mixed, centrifuged at 4℃ for 3min at 10000g, and the supernatant was taken and placed on ice. The absorbance value A1 of ATP standard for 10s at 340nm was measured immediately, and then the reaction solution with cuvette were put into a water bath at 25℃ for 3min, and was measured immediately at 3min10s obtained the absorbance value A2. The δAi (determination tube) =A2 determination tube was measured and calculated -A1 measurement tube; δAs (standard tube) =A2 standard tube -A1 standard tube. The ATP content was calculated according to the sample quality (W): ATP content (μmol/g mass) =0.625× δAi ÷ δAs ÷W. The plant funtional leaves at the top third to fouth position were were sampled at AM 9:00 to 11:00 used for ATP detection under normal field conditions.

**NADPH content**: Under the direction of Coenzyme II NADP^+^/NADPH Content Kit Instruction (www.geruisi-bio.com) preparation all Reagents include of Extract A and B solution, Reagent 1 to 4 and NADPH standard solution. Take about 0.5g functional leaves of cassava, add 1mL of extract B, and grind in ice bath. All the samples were transferred to EP tubes (supplemented with extract B to 1mL), and the samples were incubated at 95℃ for 5min. After removal, they were immediately placed in an ice bath (or refrigerator) for 5min. Centrifuge 12000rpm at 4 ℃ for 10min; 500μL of the supernatant was put into a new EP tube, and then V2 volume of the extract was added for neutralization (adjusted to PH about neutral). Centrifuge at 12000rpm for 5min at 4 ° C, and take the supernatant and place it on ice to be tested. Detection with microplate reader: Mix the assay solution and reagents according to the instructions, the absorbance value A1 was measured immediately at 37℃, 450nm, and then A2 was measured 30min later, δA =A2-A 1. The NADPH content was calculated according to the formula y = 8.6631x-0.0013 (x is NADPH molar mass (nmol), y is δA) of the standard curve. NADPH (nmol/g)=[(△A+0.0013)÷8.6631] ÷(W÷2×V sample÷V4), W: fresh weight of samples(g); V: sample volume added to the reaction system, 0.02mL; V4: NADPH extraction liquid volume, 0.5mL extract B+ V2mL extract A= (0.5+V2) mL. The cassava leaves used for NADPH detection were sampled at AM 9:00 to 11:00 under normal field conditions.

**Measurement of phytohormones**

Fresh plant materials were harvested, weighted, immediately frozen in liquid nitrogen, and stored at -80℃ until needed. Plant materials (50 mg fresh weight) were frozen in liquid nitrogen, ground into powder, and extracted with 0.5 mL methanol/water/formic acid (15:4:1,V/V/V) at 4℃. The extract was vortexed (10 min) and centrifuged at 12,000 rpm under 4℃ for 5 min. The supernatants were collected, repeat the steps above, vortexed (5 min) and centrifuged (5 min). The combined extracts were evaporated to dryness under nitrogen gas stream, reconstituted in 80% methanol (V/V), ultraphoniced (1 min) and filtrated (PTFE, 0.22 mm; Anpel) before LC-MS/MS analysis. The sample extracts were analyzed using an LC-ESI-MS/MS system (UPLC, Thermo, USA; MS, Q Exactive, Thermo, USA, https://www.thermofisher.com). The effluent was alternatively connected to an ESI-triple quadrupole-linear ion trap (Q TRAP)-MS. The analytical conditions were as follows, UPLC: column, Waters HSS T3（50*2.1 mm, 1.8μm）; column temperature, 40 ∘C; flow rate, 0.3 mL/min; injection volume, 2μL; solvent system, water (0.1% Acetic acid): acetonitrile (0.1% Acetic acid) ; gradient program, 90:10 V/V at 0 min, 90:10 V/V at 1.0 min, 90:10 V/V at 7.0 min, 90:10 V/V at 7.1 min, 90:10 V/V at 9.0 min. Data were acquired on the Q-Exactive using Xcalibur 4.1 (Thermo Scientific), and processed using TraceFinder™4.1 Clinical (Thermo Scientific). Quantified data were output into excel format.

**Quantification of cyanogenic glycosides in cassava**

Fresh storage roots and leaves were sampled and immediately frozen in liquid nitrogen, and stored at -80℃ until needed. Weigh 2.0g sample, put it into a triangle bottle with stopper, add 100ml water, seal it, place it at room temperature for 4h, transfer it into a distillation bottle, add 100ml water, 20ml zinc acetate solution (100g/ L), and distill 1-2G tartaric acid. Receive about 45ml with a small beaker containing 5mlNaOH (5%). Transfer to a 50ml volumetric flask and adjust to 50ml (V1) with distilled water. Take 1ml(V2) in 25ml colorimetric tube, make a blank with distilled water, add distilled water to 10ml, add 0.5ml 2% NaOH, 1 drop phenolphthalein, adjust with acetic acid solution (1:24) until the red color disappears, add 5ml phosphate buffer pH=7, heat to 37℃ for 10min, add 0.25ml chloramine T, Add the stopper, mix and shake, place for 5min, add 5ml pyrazolone isonicotinate solution, add water to 25ml mix, place at 20-40℃ for 40min, colorimetric absorbance value A at 638nm. The total amount of hydrocyanic acid was quantified by reference to the hydrocyanic acid standard curve.

**Quantitative real-time PCR validation**

The expression of important genes in photosynthesis, differentiation of flower primordia and storage roots and cyanide glycoside metabolism has been validated by real-time Q-PCR. The total RNA of samples was extracted using a TRIzol Kit (Invitrogen, Cat#15596018) for three biological repeats according to the user’s manual. About 1.5 mg of each RNA sample was used for cDNA synthesis (Maxima H Minus cDNA Synthesis Master Mix, with dsDNase, Thermo Scientific, Cat#M1682). Quantitative real-time PCR was performed using primer pairs (see **S Table13**) in the SsoFast EvaGreen Supermix (Bio-Rad, Cat#1725201) with the real-time PCR detection system (Bio-Rad, CFX96). PCR reactions were performed in quadruplicate for each sample, and expression levels were normalized to OaUBI.

**Fluorescence *in situ* PCR**

Fresh unfolded leaves, storage roots from cassava plants DAP 120-150 are collected In tube used for in situ PCR shown tissue specific expression of *MeGTR*, *MeMATE1* involved in cyanide transport, and *MeCOL1*, *MeTFL2* regulated flowering and storage root development. The protocol referenced as described previously (Jørgensen et al., 2011). Prior to sectioning, all tissues were fixed in FAA (2% formaldehyde, 5% acetic acid, and 63% ethanol in phosphate-buffered saline [PBS]) for 5 h at 4℃ and then washed three times with washing buffer (60% ethanol and 5% acetic acid in PBS). The tissue was embedded in 5% agarose in PBS to enable sectioning into 80um tissue sections on the Leica freezing microtome (LEICAVT1200S). The sections were immediately placed into 20ul enzyme-free water containing 4U RNase inhibitor and 40U DNase, incubation at 37℃ for 4 hours. Wash the sections with enzyme-free water twice, then soak into Pepsin (2ng in 1ul 10mM HCl) for about 15min, again wash the sections twice with enzyme-free water. The mRNA was translated to cDNA using 1mM the specific reverse primers for each gene mixing with1xRT Buffer, 2mM dNTP taking reaction at 65℃ for 5min, adding RNase Inhibitor and Sensicript at 4℃, return reaction at 45℃ for 60min, 97℃ for 1min until store at 4℃. Before the cDNA was amplified, the reverse transcriptase solution was removed and the PCR reagents were added. DIG-labeled dUTP was incorporated as the basis for visualization. After PCR amplification add 175ul of anti-DIG-POD working Solution at 37℃ and gently shake the shaker for 30min, wash 4 times with washing buffer. Discard the solution, add 175ul of ABTS solution, and gently shake the shaker for 30min at 37℃ under dark environment. Clean the sections and examine them by laser scanning confocal microscope（LEICA/TCS SP8）.

**Yeast one-hybrid assay**

Yeast one-hybrid assay was performed according to the method described by Li et al (2016). The yeast one-hybrid bait vector pAbAi-target gene (*MeCYP79D2*, *MeCOL5*) was constructed, and the cassava leaf total RNA was isolated. Then, the double-stranded cDNA was synthesized following the protocol as previous study (Ma et al. 2017). The double-stranded cDNAs were linked with Sma I linearized prey vector pGADT7-Rec, transformed into the yeast positive strain Y1HGold/pAbAi target gene, and then cultivated on SD/-Ura-Leu medium with right Aureobasidin A (AbA) concentration at 30℃ for 3 d. The positive colonies were identified by plasmid PCR and sequencing. To confirm the interaction between TF (*MebHLH2*, *MebHLH4*, *MeFT1，MeFT2* and *MeTFL2*) and target gene, the full-length TF coding sequence (CDS) was cloned by PCR. The CDS was ligated into pGADT7 vector through Nde I and Sal I and named pGADT7-TF. pGADT7-TF and pAbAi-target gene were co-transformed into the Y1HGold yeast strains. pGADT7-Rec53+p53-AbAi, pAbAi-target gene, pGADT7-TF, and pGADT7-TF+pAbAi were used as controls. The transformed cells were grown on SD/-Leu selective medium with right AbA concentration at 30℃ for 3 d.

**QUANTIFICATION AND STATISTICAL ANALYSES**

All details of the statistics applied in this study are provided alongside the respective analysis in the Method Details section. The t-test and Fisher exact test were performed using R 3.26 package.
