## Supplementary Information for "Pan-genome and Haplotype Map of Cultivars and Their Wild Ancestors Provides Insights into Selective Evolution of Cassava (*Manihot esculenta* Crantz)"

[Fig. S22. Fst and ROD analysis of population selection: The red broken line is a significant region containing the](#_Toc117449229) *[MeTFL2](#_Toc117449229)* [gene, and domestication-related signals are detected in both Fst and ROD detection	39](#_Toc117449229)

### Supplementary Note 1. Cultivated cassava clones and accessions of its wild relatives selected across continents

In order to analyze the domestication of important agronomic traits of cultivated cassava, we selected 496 wild and cultivated types of cassava from Central and South America, Southeast Asia and Africa. Among them, 12 duplicated clones were eliminated by re-sequencing analysis, and the remaining 484 were used for deciphering the population genetic variations. These materials include: 278 cultivated clones and 3 wild accessions conserved in the National Cassava Germplasm Bank of China; 15 wild species from EMBRAPA, Brazil, and 4 landraces originated from Ecuador provided by CIAT. Other sets of accessions of 179 cultivated clones and 17 wild relatives were obtained by downloaded resequencing data from the Genbank (**Ramu *et al*, 2017**) (**Table S15, Fig. S2**). Three complete genomes were constructed by AM560, an artificial selfing S3 generation line of cultivar, and wild ancestors FLA4047 and W14, which were used as reference genomes for comparative analysis of genetic variation between wild and cultivated populations. The above three genomes and another 21 representative cultivated lines were selected to construct a pan-genome for SV evolution analysis between wild progenitor and cultivated populations (**Fig. S1**).

### Supplementary Note 2. Genome size evaluation

Flow cytometry was used to evaluate the genome size of several cultivars and wild species. The results showed that the genome size of cultivar KU50 and near wild ancestor FLA4047 were 0.72±0.02 Gb and 0.72±0.02Gb, which were almost consistent with each other. The genome size of W14 was 0.83±0.016Gb larger than the above clones, and that of GLAyn and GLAb in *M.glaziovii* were 0.73±0.016Gb (**Table S1**).

### Supplementary Note 3. Complete genome assembly, correction and validation

For complete genome assembly of AM560 and FLA4047, 100X Nanopore long reads were independently assembled into contigs (<https://github.com/ruanjue/smartdenovo>) and aligned Illumina reads against these genome assemblies using FALCON. Then the assemblies were took hybrid scaffolding with BioNano optical physical mapping data to the chromosomal level. For complete genome assembly of W14, 200X PacBio single-molecule long-reads were assembled into contigs, which were subsequently used to construct scaffolds using Bionano optical mapping. The total assembly length of AM560, FLA4047 and W14 is range from 730 Mb, 674 Mb to 681Mb. The scaffold N50 of the three cassava genomes ranges from 36 Mb to 39 Mb, which indicates that the genome assemblies approach the chromosome-level completeness (**Table S2**). After that, two cells of ONT ultra-long reads sequenced for AM560, FLA4047 and W14 genomes were added and integrated into the former assemblies respectively. These allow us resulted almost complete reference genome sequences of the three representative genotypes. Total assembly length is 645Mb，672Mb and 709Mb with scaffold N50 36.1Mb, 35.4Mb and 40.7Mb respectively (**Table 1**). The completeness of the assemblies was evaluated using the annotated genes collected from theBenchmarking Universal Single-Copy Orthologs (BUSCO) program (v3.0.1). The completeness of the three cassava genomes is 97.7%, 96.9% to 97.5%, respectively **(Fig. S9).** The protein coding genes were predicted using a combination of homology-based search and *ab initial* prediction methods. The genome of AM560, FLA A4047 and W14 consist of 30,571, 33,913 and 32,406 protein-coding genes, respectively (**Table 1**). By searching against public databases, we assigned these protein-coding genes to KEGG pathways and Gene Ontology terms.

### Supplementary Note 4. Optical map, scaffolding and pseudomolecules construction

**Optical map construction**

The ultra high-molecular-weight (uHMW) DNA was isolated from young leaves of *Manihot esculenta* genotype W14 and AM560 by the Amplicon Express (Pullman, WA, USA). Two optical maps using NLRS (nick, label, repair, and stain) and DLS (direct label and stain) technologies, respectively, were constructed for W14, and one optical map using the DLS technology was constructed for AM560.

For the NLRS method, the nicking endonuclease Nt.*Bsp*QI (New England BioLabs, Ipswich, MA, USA) was selected to nick uHMW DNA molecules at specific sequence motifs based on publicly available W14 genome sequence (Wang et al., 2014). The nicked DNA molecules were then labeled and stained according to the instructions of the IrysPrep Reagent Kit (Bionano Genomics, San Diego, CA, USA) as described in detail in Luo et al. (2017). The DNA sample was loaded onto the nanochannel array of an IrysChip (Bionano Genomics, San Diego, CA, USA) and was automatically imaged by the Irys system (Bionano Genomics, San Diego, CA, USA). Raw DNA molecules >20 kb were collected and converted into BNX files by AutoDetect software to obtain basic labelling and DNA length information. Those molecules >180 kb and also passing quality control were aligned, clustered, and assembled into an optical map using the Bionano Genomics assembly pipeline (Bionano Genomics, San Diego, CA, USA). The thresholds used for pairwise assembly, extension/refinement, and final refinement stages were 1 × 10^−8^, 1 × 10^−9^, and 1 × 10^−15^, respectively.

For the DLS method, the uHMW DNA molecules were labeled with the DLE-1 enzyme (Bionano Genomics, San Diego, CA), and were stained according to the instructions of the Bionano Prep™ Direct Label and Stain (DLS) Kit (Bionano Genomics, San Diego, CA). A consensus optical map was *de novo* assembled using the Bionano Solve v3.3 package with significance cutoffs of P < 1 × 10^-8^ to generate draft consensus contigs, P < 1 × 10^-9^ for draft consensus contig extension, and P < 1 × 10^-15^ for final merging of the draft consensus contigs. A recipe of “haplotype”, “noES”, and “noCut” was chosen.

The initial optical map constructed by using either method was then checked for potential chimeric contigs and further refined.

**Optical map validation of sequences and resolving of chimeras**

To compare sequence assemblies with the optical maps, the W14 sequences were digested *in silico* with Nt.*Bsp*Q1 nickase using the Knickers (Bionano Genomics, San Diego, CA, USA), and aligned on the corresponding NLRS optical map using RefAligner with alignment cutoff of P < 1 × 10^−10^ (Bionano Genomics, San Diego, CA, USA). Likewise, the AM560 sequences were digested *in silico* with the DLE-1 restriction site information and aligned to the corresponding DLS optical map. The mis-assembled sequences were then disjoined accordingly.

**Optical map guided sequence scaffolding**

The conflict-free sequences were scaffolded using the Stich algorithm (Shelton et al., 2015) with guides of the optical maps. The filtering parameters of Stitch were trained to be suitable for the current datasets, and Stitch was performed in iterations until no additional scaffolds could be produced. The gaps were filled with the respective number of N’s by using the estimated length between the flanking restriction sites. For W14 genome, the NLRS optical map was applied first, the resulting sequences were further scaffolded with the DLS optical map (**Table S3**).

**Pseudomolecule construction**

The information of annotated genes of AM560 genome (Bredeson et al., 2016) was used to determine the order and orientations of the scaffolds on each chromosome in both W14 and AM560 genomes. These scaffolds were then linked with 100 N’s accordingly and anchored onto the chromosomes.

**Gap closure of W14 genome assembly**

The gap closure was performed for W14 genome using PacBio contigs and optical maps. The PacBio contigs were validated by the two optical maps, only those contigs that agree with both optical maps were used to close gaps. The conflict-free PacBio contigs were aligned to the W14 genomes using BLAST and substituted the N bases with corresponding nucleotides with the aid of optical maps, and the final W14 pseudomolecules were generated.

**Identification of large-scale inversions and translocations between W14 and AM560 genomes**

Gene collinearities were used to identify inversions and translocations between W14 and AM560 genomes. We transferred the gene annotations of v7 (Bredeson et al., 2016) to the newly assembled AM560 pseudomolecules to acquire new coordinates of the genes on AM560_2020 using BLAST with default settings. The coordinates of each gene on the AM560_2020 were obtained from the best alignment based on the maximum coverage and similarity. These genes were then mapped to the W14 pseudomolecules produced in this study using BLAST with default settings. The coordinates of top hit were recorded. Collinearity of top hits in the W14 genome was ordered according to the order of genes along the AM560_2020 pseudomolecules. An inversion was determined if three or more genes were in a descending order; a translocation was determined if three or more genes interrupted a sequence of collinear genes.

### Supplementary Note 5. Hi-C library sequencing, scaffolding and chromosome assembly

We performed denovo sequencing and assembly of high-quality genomes at the chromosome level on wild accessions FLA4047, W14, and cultivated variety AM560. Among them, W14 used an average coverage depth of 80X's SMRT single-molecule real-time sequencing and Bionano optical atlas technology mount the contig to the chromosome level. The final assembly result is 751.98Mb, and the contigN50 is 7.76Mb (Table S2); AM560 uses Nanopore technology with Bionano-assisted assembly resulted in genome of 675.64Mb, and the contigN50 was 6.13Mb (**Table S2**); Nanopore sequencing was used for FLA4047 and high-throughput chromosome conformation capture (Hi-C) library was constructed at the same time. The FLA4047 genome is 1,233.45Mb, and the contig N50 is 4.09Mb (**Table S2**). We used BUSCO to evaluate the completeness and accuracy of the three genomes assembly, and found that the BUSCO evaluation exceeded all of them in the genome assembly with 97.7%, 96.9% to 97.5% respectively for AM560, FLA4047 and W14.

### Supplementary Note 6. Pan genome sequencing and assembly

21 representative accessions include of 20 cultivars and one accession of *M. glaziovii* GLAyn have been sequenced with Nanopore ONT long reads (>50 kb) platform and assembled independently (**Table S5**). We obtained high quality assembly with scaffold N50 of 24.67Mb to 36.28Mb in size of 638.07Mb to 849.08Mb, annotated 30245 to 41950 gene models (**Table S6**). Togather with three reference genomes, we constructed the pan SV genome of cassava species (**Fig.S11, Fig. S12**). These 24 accessions with name of SC6068, SC8, SC9, Taiyin No. 1, Wenchang Red, ZM9781, SC205, SC5, Mianbao, Fuxuan 01, HB60, Brazil-1, BC008, BC006, ITBB04, IB003, IB005, NZ199, GR4, GLAyn, GLAb and three accessions of AM560, FLA4047 and W14 used for references were all collected in the germplasm resource nursery in Chengmai County, Haikou City.

### Supplementary Note 7. Core and dispensable genes in the pan genome

We performed pan-genomic analysis on the assembled genomes of 3 wild relatives and 21 cultivated cassava varieties. Orthologs investigation classified all genes from the 24 cassava genomes into 30,517 families. With the increase in the number of genomes, the total gene set gradually increased, and the number of genomes increased to 22, and then stabilized (**Fig. 2**), indicating that these 24 cassava genotypes are relatively representative toward to a pan genome of cassava. In all 24 genomes, 6,386 gene families were defined as core genes, 5,011 families in 22 to 23 clones were defined as soft core genes, and 18,315 families in 2 to 21 clones were defined as dispensable genes , 808 genes only appeared in a single genome are defined as private genes (**Fig. 2；Table S7**). Although core genes only account for ~21% of gene families, they account for ~38.28% of the genome. Dispensable accounts for ~60% of gene families and ~39.29% of a single genome (**Fig. 2B**), We performed KEGG enrichment analysis on function of the above gene columns, found that the core gene is considerable carbon metabolism and spliceosome (**Fig. S13a**), softcore gene set is mismatch repair, RNA degradation, non-homologous end-joining (**Fig. S13b**); and the dispensable set is ascribed into sesquiterpenoid and triterpenoid biosynyhesis, stilbenoid, diarylheptanoid and gingerol biosynthesis, ribosome biogenesis in eukaryotes (**Fig. S13c**). The private gene set is involved in butanoate metabolism, pyrimidine metabolism, synthesis and degradation of ketone bodies (**Fig. S13d**).

### Supplementary Note 8. Structural Variation (SV) Analysis

Using AM560 v7.0 assembly as the reference genome, we have counted the gene density and repetitive sequence information among 24 cassava genomes. Detection of SV with sniffles called out all 5 types of SVs include of insertions (INS), deletions (DEL), translocations (TRA), inversions (INV) and duplications (DUP) as in **Fig.S11**. Further statistics found that the proportion of DEL (51.05%) and INS (23.32%) is higher, then TRA (18.33%), DUP (5.5%) and INV (1.81%) in total (**Table S8**). SV size analysis showed that the maximum length of DEL was larger than that of INS, followed by INV and DUP (**Fig. S11c**). Among the 21 cultivated cassava varieties, ZM9781SV had the least variation, while GR4 had the most SV variation (**Fig. S11d**).

We counted the number and fragment size of INS in 21 cultivated cassava plants, and found that the number of inserted fragments ranged from 2734 to 7391bp, with the smallest insertion size being 674,329 bp and the largest being 1,923,718 bp, and the average insertion size being ~243bp (**Fig. S12**).

### Supplementary Note 9. Present /absent of genes and copy number variation from wild ancestors to cultivars

We examined the expansion and contraction of gene families occurring between the three wild ancestors FLA4047, W14 and GLAyn named FLA, to 21 cultivated clones named ME (**Table S17**). There were 382 homologous gene families, involving 1015 genes, were uniquely presented in the FLA, which were lost to varying degrees in modern cultivated varieties. There were 6812 gene families involving 34,241 genes presented in ME only. GO enrichment analysis shown that genes special in FLA are adversity adaptation related, like biosynthesis of secondary metabolites, photosynthesis, cysteine and methionine metabolism those sulfur-containing amino acids (**Fig. S14**). The gene and gene families expanded in cultivated cassava mainly include of biosynthesis of secondary metabolites, starch and sucrose metabolism, amino sugar and nucleotide sugar metabolism, protein processing in endoplasmic reticulum, RNA transport. In addition, MAPK signaling pathway, ABC transporter and fatty acid degradation are found enriched (**Fig.S14**). CNVs considerable occurred in photosynthesis, starch and sugar metabolism, hormone metabolism and signaling, and part of the transcription factor families. In photosynthesis, 13 gene families in 30 KEGG pathways of light reaction pathways expanded to varying degrees in cultivars, mainly involving photosystem I-related *psaB* (K02690), *psaK* (K02698) and Reaction Center Subunit IV-B *psaE* (K02693); Also Photosystem II associated with *psbB* (K02704), PSII 43 kDa protein gene *psbC* (K02705), coding the 10 kDa protein *psbR* (K03541) and Photosystem II P680 Reaction Center protein *Psb28* (K08903) (**Fig. S15-S16, Table S18**). In sucrose and starch metabolism, CNVs in large scale were found in 17 of 31 gene families. Gene expansion significantly happened in alpha-1,4 glucan phosphorylase L-2 isozyme (K00688), alpha-glucosidase-like (*STP-1*, K01187), sucrose synthase (K00695), beta-glucosidase 40 (K01188) and beta-glucosidase 18 (K05350). The other 14 gene families showed some shrinkage in cultivated species, such as beta-amylase 7 (K01177), endoglucanase 12 (K01179) and phosphoglucomutase (*PGM*, K01835) (**Fig.S17-S18; Table S18**). Several significant gene expansion events were found in phytohormone metabolism, such as brassinosteroid insentitive 1-associated receptor kinase 1 (*SERK3*, K13416), regulatory protein *NPR3* (K14508) and transcription factor *PIF4* (K6189). Other gene families contracted in different extents, such as, mitogen-activated protein kinase (*STMEK2,* K13413), transcription factor *MYC2-like* (*MYC2,* K13422), auxin-responsive protein *SAUR36* (K14488), abscisic acid receptor *PYL4-like* (K14496) and ethylene insensitive 3-like (*EIN3*, K14514) (**Fig. S19, Table S18**). Comparison of homologous genes of major expansion and contraction transcription factors in cassava showed that FLA retained a large number of transcription factors related to stress, such as *bHLH*, *C2H2,* *C3H* and *ERF*. The transcription factors of ME expansion mainly included *bHLH*, *SBP* and M-type *MADS* (**Fig. S20, Table S18**).

### Supplementary Note 10. Population structure and diversity analysis

Based on re-sequencing data of 484 accessions include of 441 cultivated clones, 21 wild progenitor accessions (FLA) and 22 GLA and other relatives of cassava, total of 90,373,720

SNPs, 21,562,482 InDels have been identified. By clarified and removed redundant, there were 6,021,758 SNPs used for construction of classification trees. At K=5, the whole population was divided into five large groups (**Fig. 3C**):  Group A consists of 22 *M. grazovii* and other relative accessions including Tree Cassava, GLZyn, GLZb, W14 and hybrids with cultivated cassava.  W14, former we sequenced, own the background of GLZ, but PCA cluster showed it might be a hybrid with cultivated species supported by it could produce storage root low in starch. That means GLZ was one of the few cassava species exchanged genes with cultivated cassava in natural evolution (Bredeson, 2016). Group B includes 22 accessions of FLA, which comes from several states on the southern edge of Amazon in Brazil. Group C is the domesticated type nearest to FLA, including 43 landraces distributed in Ecuador and their hybrid offspring with cultivars. Group D is cultivars with two subgroups: D1 consist of 68 clones, which are landraces and hybrid progeny distributed in GO and MT states in central and western Brazil. D2 cover 85 genotypes, with sugary cassava, SC5, SC8 and their derived hybrids named K series.  Group E covered 248 lines could be further divided into 7 subgroups E1-E7 according to their genetic distance: E1 and E2 total 56 lines deputy of cultivated germplasm from Southern Brazil, and the relatively cold tolerant clones such as SC124, F201, CH16, etc.  E3 includes of several specific varieties from Parana state of Brazil, as well as many edible clones from Asian and Africa countries, which could be assigned to the geographic subtype of southern Brazil; E4 consist of landraces and hybrids from Uganda in Eastern Africa, and a few African collections introduced from Colombia.  E5 is mainly landraces collected from Nigeria, Uganda western Africa and their hybrid progenies. E6 and E7 all 71 accessions are landraces and their hybrids from West Africa (**Table S16**).  According to PCA analysis among the species or subspecies (**Fig.3B upper**), accessions from FLA, GLA distributed away from the accessions of ME clearly. Inside the ME, the cultivated lines from group C, D and E merged in each other, can’t be divided into independent groups (**Fig.3B down**).

### Supplementary Note 11. Selective sweeping analysis

We compared the nucleotide diversity whole genome wide between population consisted of 198 accessions of cultivated cassava (ME) to population of *flabelifolia* with 28 accessions (FLA). Total of 13,735,173 and 22, 032,832 SNPs were obtained from 298 cultivated accessions and 28 wild accessions respectively (**Table S9**). These SNPs mainly distributed in no-gene region, and in gene region prefer to intron than in CDS and UTR region **(Fig. S21)** not only in ME but also in FLA. Also 2,719,151and 4,533,604 InDels were found in cultivar population and wild population, they mainly distributed in the non-gene region, and concentrated in intron rather than CDS and UTR in the gene region. By calculated Fst and ROD values along each chromosome within 100Kb window, there were 1519 and 202 potentially selected genes have been identified under the top 5% and 2% of ROD and Fst, respectively (**Fig. S21; Table S19-21**). These genes are important candidate genes selected during domestication from wild ancestor to modern cultivars.

Go term enrichment of the 1509 genes involved in selective sweeping shown that the considerable genes are being for regulation of transcription, response to water deprivation, protein autophosphorylation in biology process, nucleus, cytoplasm, plasma membrane and chloroplast in cell cycle, and protein binding, DNA-binding transcription factor activity, protein serine/threonine kinase activity and so on in molecular function (**Fig.S23** upper). KEGG terming further found that genes for biosynthesis of second metabolites, metabolic pathways, ubiquinone and other terpenoid quinone biosynthesis, thiamine metabolism, alpha linolenic acid metabolism and plant hormone signal transduction were highly enriched (**Fig.S23** down), that indicated these genes have been predominantly selected in the domestication of cassava.

### Supplementary Note 12. Structural evolution and differential expression validation of genes in C_3_-C_4_ intermediate photosynthesis from wild ancestors to cultivars of cassava

Based on selective sweeping analysis, A systematic comparison of the gene and its upstream 2-4Kb promoter sequences only for photosynthesis between 198 cultivated clones to 28 FLA relative wild ancestor accessions was carried out (**Table S23，Table S23**). We found that definite insertion & deletions and SNP mutations in the promoter regions of *MeCSK* (13G022900), *MeLHCB2.1* (01G175700) and *MeFNR*3 (02G10500) among the 14 genes related to photochemical reactions. Only the promoter regions of *MeNADP-ME* (11G034000) associated with carbon fixation showed varying degrees of deletion and structural variation. It was found that there was an 83bp transposon insertion 390bp upstream of the start codon of *MeCSK* (13G022900) in most cultivated cassava cultivars (167/198), while only 1 (1/28) of wild subspecies FLA had the same transposon insertion. A SINE element of length was inserted at 364bp upstream of the start codon of *MeLHCB2.1* (01G175700) in most members of the cultivated group (160/198), while it was almost absent in the wild species FLA group (1/28). There were also multiple insertion and deletion variants upstream of the promoter of photosynthetic electron transport terminal *MeFNR3* (02G105100) synthesized by coupled reducing NADPH (**Fig.4F**, **Table S23**). There was a deletion of a 107bp transposon (24/30) at -568bp upstream of the start codon of C4 carbon fixation-related *MeNADP-ME* (11G034000) gene in the ME population, and the deletion was mostly inserted (19/28) in the FLA. In addition, the structural variation in CDS region of genes for photosynthesis between ME to FLA showed that there were no mutations leading to amino acid substitution in all genes except *MeCSK*. There were 30 non-synonymous substitutions in the cds of *MeCSK* in wild species, but only 12 in cultivated species. Among them, two base substitutions resulted in amino acid variation, which occurred in the coding region of 173bp, and the frequency of C-A allele variation was 0.33, resulting in encoding alanine (A) to aspartic acid (D). The other occurred at 1632bp with a g-c variant frequency of 0.61, resulting in a variant encoding lysine (K) to asparagine (N). These two purification selection sites are involved in the change of amino acid properties, especially the change of amino acid charge properties, and their evolutionary significance is worthy of further study.

We compared their transcriptomic expression profiling of genes for photo reaction and carbon assimilation in photosynthesis of three wild relatives and 21 cultivated varieties plants DAP120 grown in the field located in Cheng Mai of Hainan. It is convinced that *MeCSK, MeFNR2, MeFNR3, MeCytb6f* and *MeATPase* that genes involved in photochemical conversion and electron transport higher expressed in cultivated clones than in wild ancestors FLA4047, W14 and GLAyn, and only *MePPDK, MePEPC* and *MeNADP-Me*, the genes of C4 enzymes high expressed in cultivars than in the wild accessions (**Fig. S24**). The quantitative RT-PCR further validated that 11 genes for light reaction include of *MeCSK、MeCytb6f、MePC、MeFNR2、MeFNR3* and *MeFd* significantly high expressed in cultivars than wild progenitors, in particular, the *MepsbA* and *MepsbD,* the chloroplast genes encode D1 and D2 proteins consist of PSII unit did. Also the considerable expression of three C4 genes, *MePPDK*, *MePEPC* and *MeNADP-ME* in cultivated clones but not in wild ancestors was confirmed (**Fig. 4E, Fig. S26**).

### Supplementary Note 13. Co-evolution of Flower primordium degeneration and root tuber initiation rooting from mutation of several key genes from wild ancestors to cultivars of cassava

Through deep literature analysis on development of floral primordium (FP) and initial of tuber and storage root (SR) of high plants, especially in Arabidopsis, rice and potato (Corbesier, 2007; Zhu et al., 2020; Daniela et al., 2020; Zierer *et al*., 2021), total of 43 candidate genes have been selected for structural variation and different expression identification in FP and SR between FLA to ME population. Screening in CDS region of the 43 genes, almost no amino acid variation in protein level found between wild accessions to cultivars in large scale accessions. Systemic comparison of sequences in promoter region of genes reveal that there were InDels variation in 19/43 genes in varying degrees, include of *MePHYB* (11G052800), *MeGI* (05G043800), *MeCOL1* (06G028200), *MeFT2* (13G000800), *MeTFL2* (13G011900), *Me14-3-3* (02G146600), *MeEFL* (04G083700), *MeMADS* (05G187800) and others in statistics (**Table S23**). We focused on *MeCOL1*, a CONSTANS gene, regarded as co-regulator of FTs (Freytes et al, 2021) and *MeTFL2,* located in selective sweeping regions could promote storage root initiation and inhibit the flowering in cassava. There were double linked deletions at -2130 to -1662 with 468bp, and at -1552 to -1276 with 276bp in promoter of *MeCOL1* for the most of the 198 cultivar and landraces but almost not in FLA accessions（1/28）(**Fig.5E**). Also a typical transposon *hAT*-MITE at -7.0kb to -8.2kb and double deletions at -1811 to -1363bp with 448bp and at -1356 to -1009bp with 347bp were identified in promoter of *MeTFL2.* There are 183/198 accessions own the *hAT*-MITE in cultivated ME group, but only 2/28 accessions in FLA (**Fig.5E, Table S22**).

We evaluated the expression of these genes candidate in developing FP and SR of specific cultivar SC205 and KU50 vs wild accession FLA4047 and W14. *MeCOL1* preferentially expressed in leaves of 4 genotypes day time, FP of wild clones and SAM of cultivars but low expressed in xylem and phloem of storage roots of all wild and cultivars. *MeFT1* and *MeFT2* predominantly expressed in leaves of 4 varieties day and night, especially *MeFT1* expressed higher in wild FLA4047 and W14 than in cultivars. Both very low expressed in SR and FP/SAM of the 4 varieties. The *MeTFL2* selectively expressed in SAM of cultivars and SR of cultivar and wild clones. *MeGI* and *MeBEL5* prefer to high expressed in leaves night time than day time but low in FP/SAM and SR of all varieties (**Fig.5C right**).We further examined the expression profiling of these candidate genes in leaves and storage roots of these wild and cultivated clones during developmental process from DAP45 to DAP97. It is shown that *MeCOL5* sustainable expressed in leaves, vein and storage root of cultivars but only expressed in leaves and vein of wild W14 at DAP79 to 97, and can be not found expression in root phloem and xylem of W14. *MeFT1* keep very low expression in leaves, vein and storage root of all wild and cultivated cassava. *MeFT2* expressed in leaves and veins of SC205, KU50 on DAP45 to DAP79, expressed in leaves and veins of W14 only on DAP79 to DAP97, but almost not expressed in root xylem and phloem of these three varieties in all six stages tested. It verified that the *MeFT2* is a storage root initial signal derived from leaves. *MeTFL2* predominantly expressed in root xylem and phloem but not in leaves and veins of cultivars during developing process, especially higher expressed in root xylem of KU50 at DAP58, and root xylem of SC205 at DAP67. It coincided with storage root earlier initial of SC205 than KU50. However, expression of *MeTFL2* in root xylem and phloem, leaves and veins of W14 was too low to detectable. Quantative RT-PCR validated the expression of these key genes in leaves, SAM/FP and storage root of KU50, SC205 FLA4047 and W14 at DAP60. Interestingly, *MeCOL5* and *MeFT1* significantly higher expressed in leaves of wild FLA4047 and W14 than SC205 and KU50 (**Fig.5G upper**). For MeFT2 (SP6A), it higher expressed in root of SC205 and KU50 than FLA4047 and W14, but its expression was lower in SAM of cultivars than FP of wild accessions，also slight lower in leaves of cultivars than wild ancestors. The *MeTFL2* significantly higher expressed in storage root, SAM and leaves of cultivar SC205 and KU50 than wild FLA4047 and W14 (**Fig. 5G down**). Summary above, the *MeCOL5,* *MeFT1,* *MeFT2* and *MeTFL2* and interaction of them are essential for differentiation of flower primordium and storage roots in cassava.

### Supplementary Note 14. Structural evolution and differential expression validation of genes in cyanide metabolism from wild ancestors to cultivars of cassava

We analyzed the structural variation of all 22 genes for cyanogenic synthesis, transportation and decomposition between 28 wild accessions and 198 cultivated clones. The results showed that common InDel variations were found only in the promoter region of *MeGTR* (15G176100), *MeMATE1* (16G007900), *MeCYP71E7* (12G132900) and *MeHNL10-like* (13G092100), and not found in other genes. Especially, significant structural variations have been discovered in promoter region of *MeGTR* and *MeMATE1*. There were 385bp insertion at -100 to -485 site and 3pb deletion at upstream -48 to -50 of *MeGTR* (**Fig.6C**); And a 1413bp insertion at -602 to -2015 position, 10bp insertion and a deletion with 11bp inside of the promoter *MeMATE1* found (**Fig. 6D, Table S23**). Comparison of CDS region of 12 genes with different expression between wild and cultivar accessions show that only *MeMATE1*, *MeCGTR1* and *MeCYP71E7* had 1-3 amino acid differences, others had no any variation in their *cds* region. For *MeMATE1,* single nucleotide substitution mutation (G→A) took place in *cds* in more than half population (54%) of cultivars predominantly higher than that (3%) in the wild accessions, This mutation may altered its transport efficiency. Meanwhile, a set of transcription factors, *bHLHs* shown systemic structural variation in their promoter region.

RNA-Seq analysis of storage roots and leaves for 20 accessions with different cyanide content revealed that genes for cyanide decomposition as *MeHNL10-like, MeHNL24-like* and *MeB-CASa*, cyanide transportation, *MeGTR, MeCGTR1, MeMATE1* and *MeH^+^-ATPase* were preferentially selective down- regulated in storage root of sweet cultivars different from bitter varieties and wild accessions, but only *MeHNL24-like, MeMATE1,* and cyanide synthesis related, *MeCYP79D2, MeUGT85K5* down-regulated in leaves of the cultivars (**Table S11, S12, S13**). It characterized the expressive specificity in roots and leaves of these genes, especially, *MeMATE1* only expressed in storage roots, and *MeGTR* mainly expressed in leaves. Quantative PCR further validated that *MeGTR, MeMATE1, MeCYP79D1, MeCYP79D2, MeHNL10-like* significantly lower expressed in pericarpium, stele of storage root and leaves of FLA4047 and W14 than in the sweet cassava KU50, SC16 and SC205 (**Fig.6B，Fig. S28**). That coincided expression identification indicated that the sequence mutation in *MeGTR* and *MeMATE1* could be one key factor cause of sweet cassava domestication.

### Supplementary Figure

#### Fig. S1. Plants, flowering and storage root forming habitus of wild ancestor and cultivars of cassava

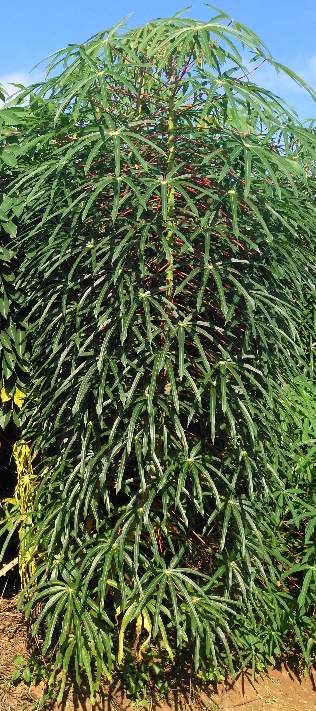

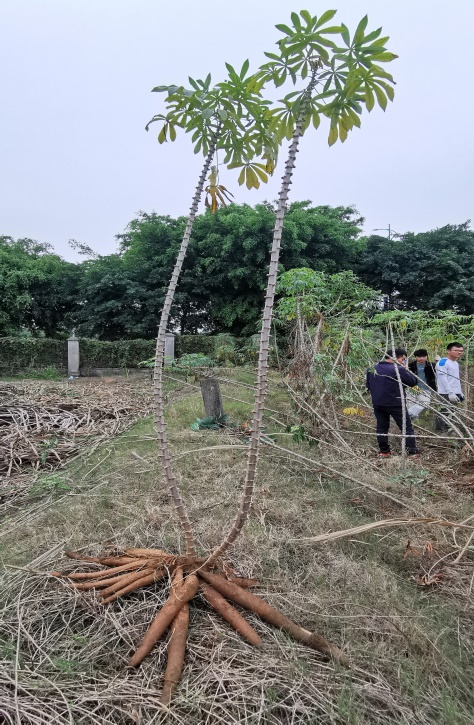

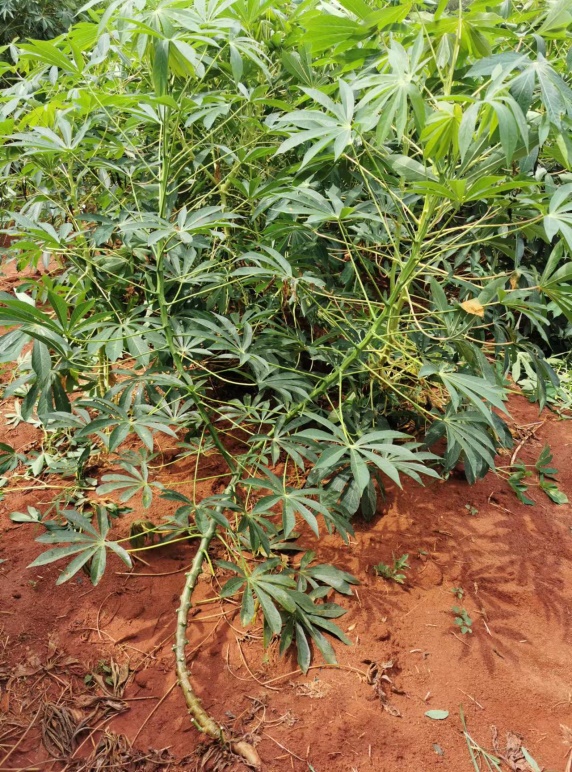

**AM 560 S3**

**SC16**

**SC205**

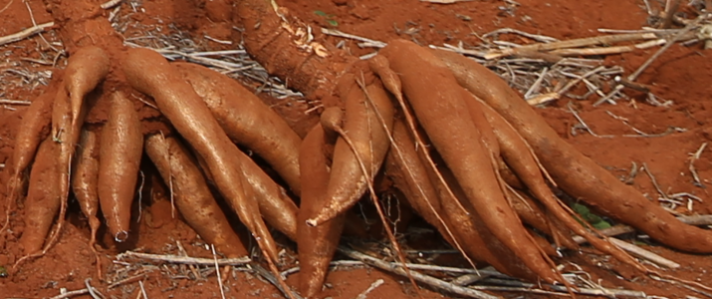

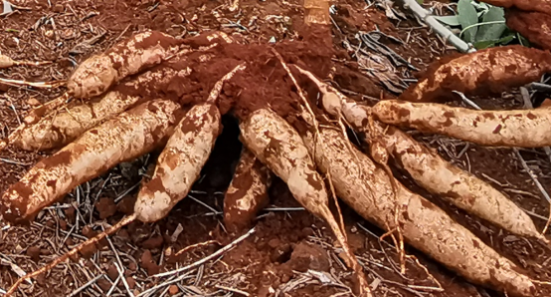

**SC205 Storage root**

**AM 560 Storage root**

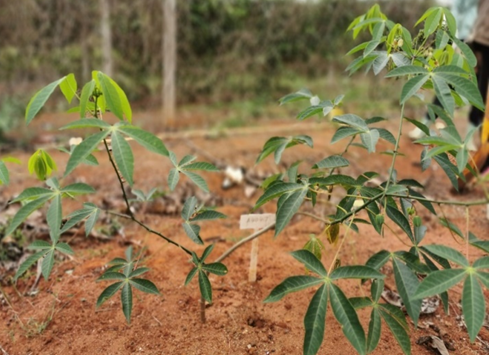

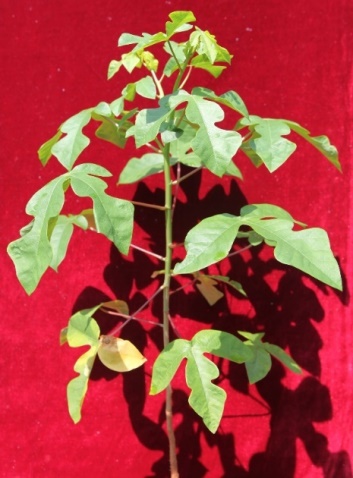

**W14**

**FLA4047**

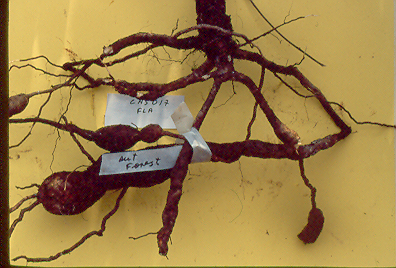

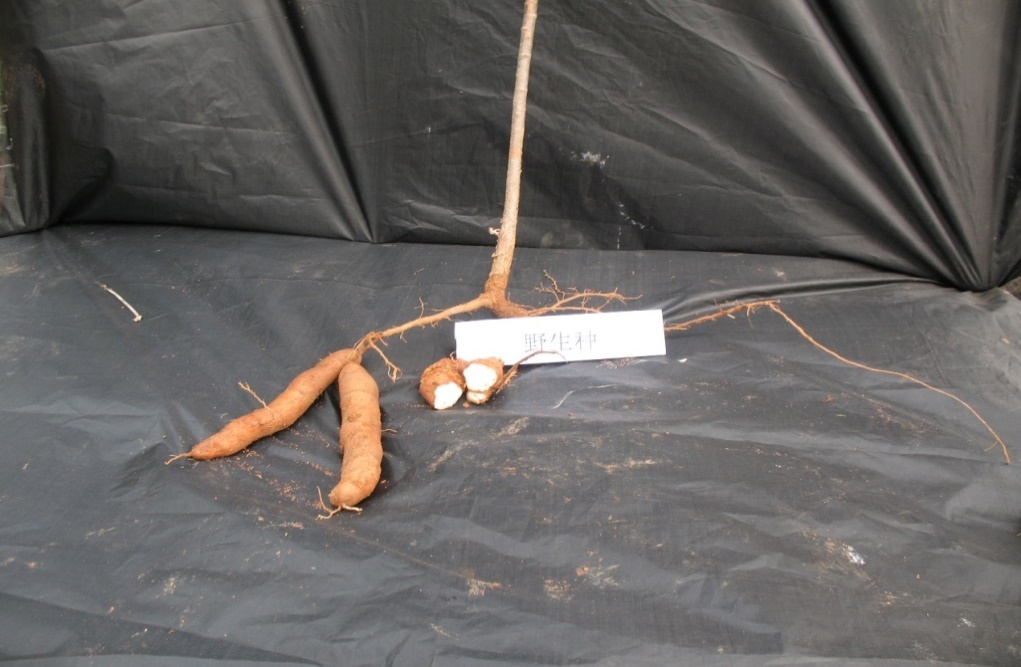

**Slightly expanded root of FLA4047**

**Storage Root W14**

Note: Upper：Plants of several cultivars (*Manihot esculenta* Crantz) not yet flowering at 180-240DAP but with larger storage roots; Down: Plant type of wild ancestors (*Manihot esculenta* SSP. *flabellifolia*) already flowered at about 40-50DAP but have slightly expanded roots (FLA4047) or very small storage root (W14).

#### Fig. S2. Geographic locations of all 484 accessions used in the study

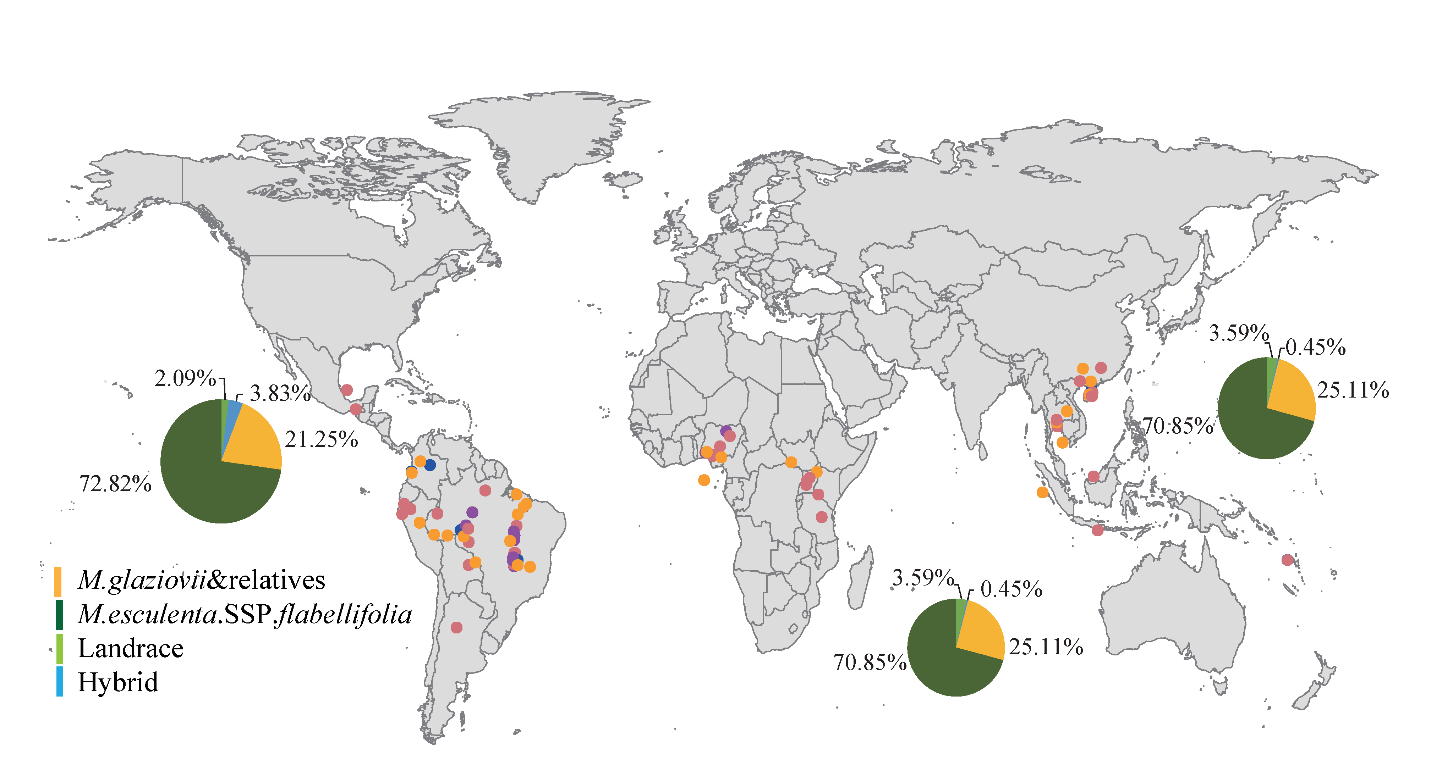

#### Fig. S3. Estimation of genome size by flow cytometry

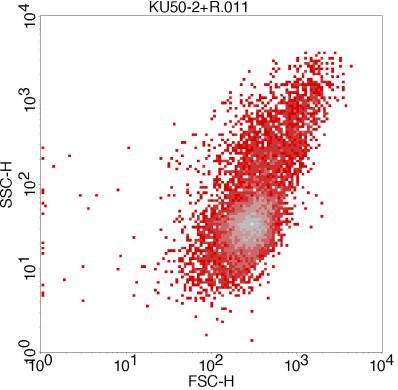

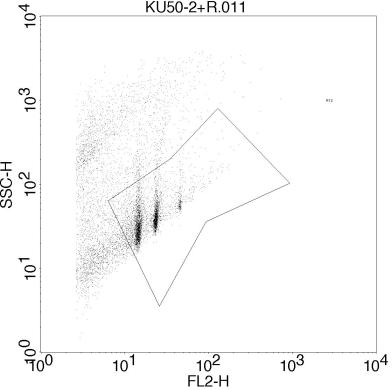

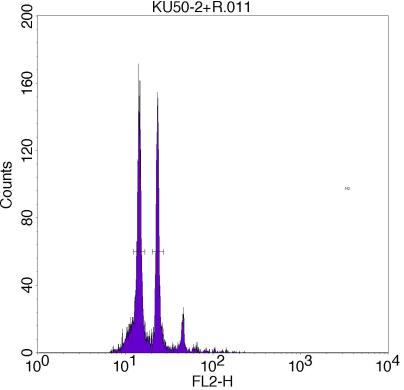

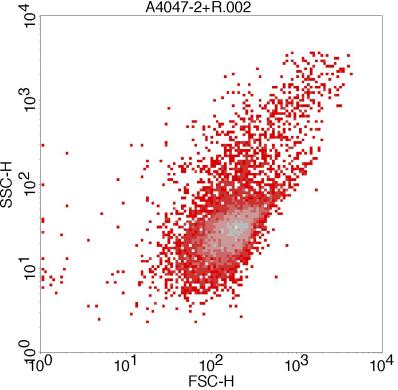

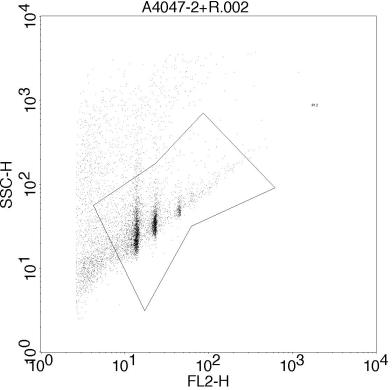

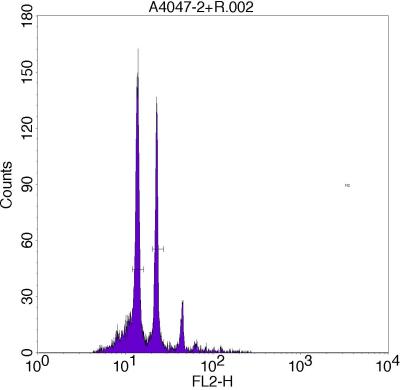

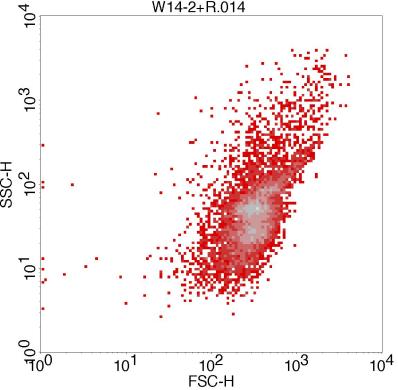

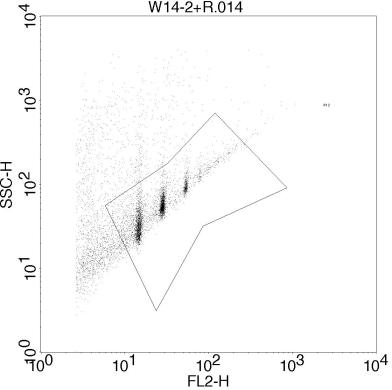

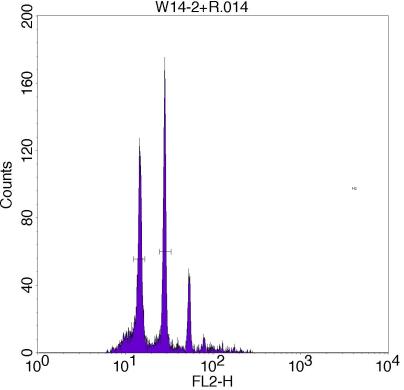

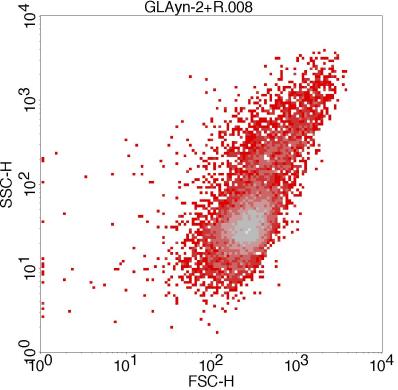

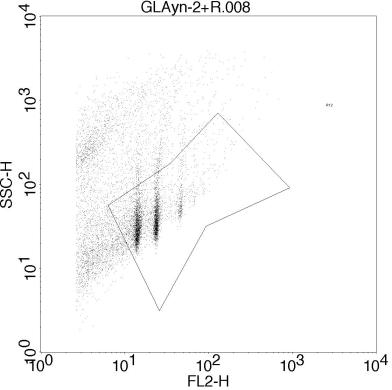

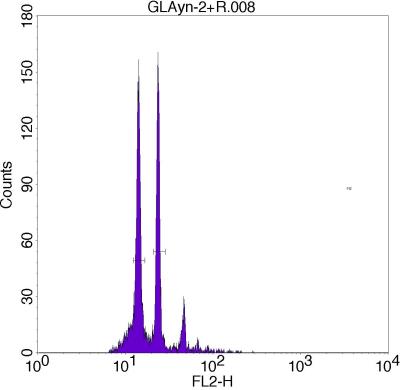

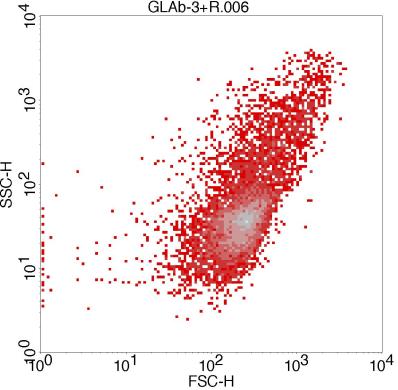

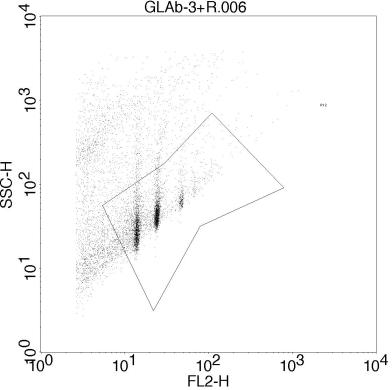

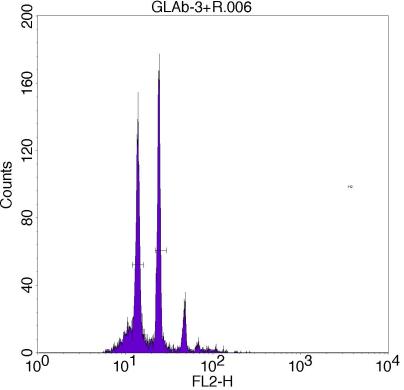

#### Fig. S4. DNA Sequencing and Data Processing Workflow pipeline

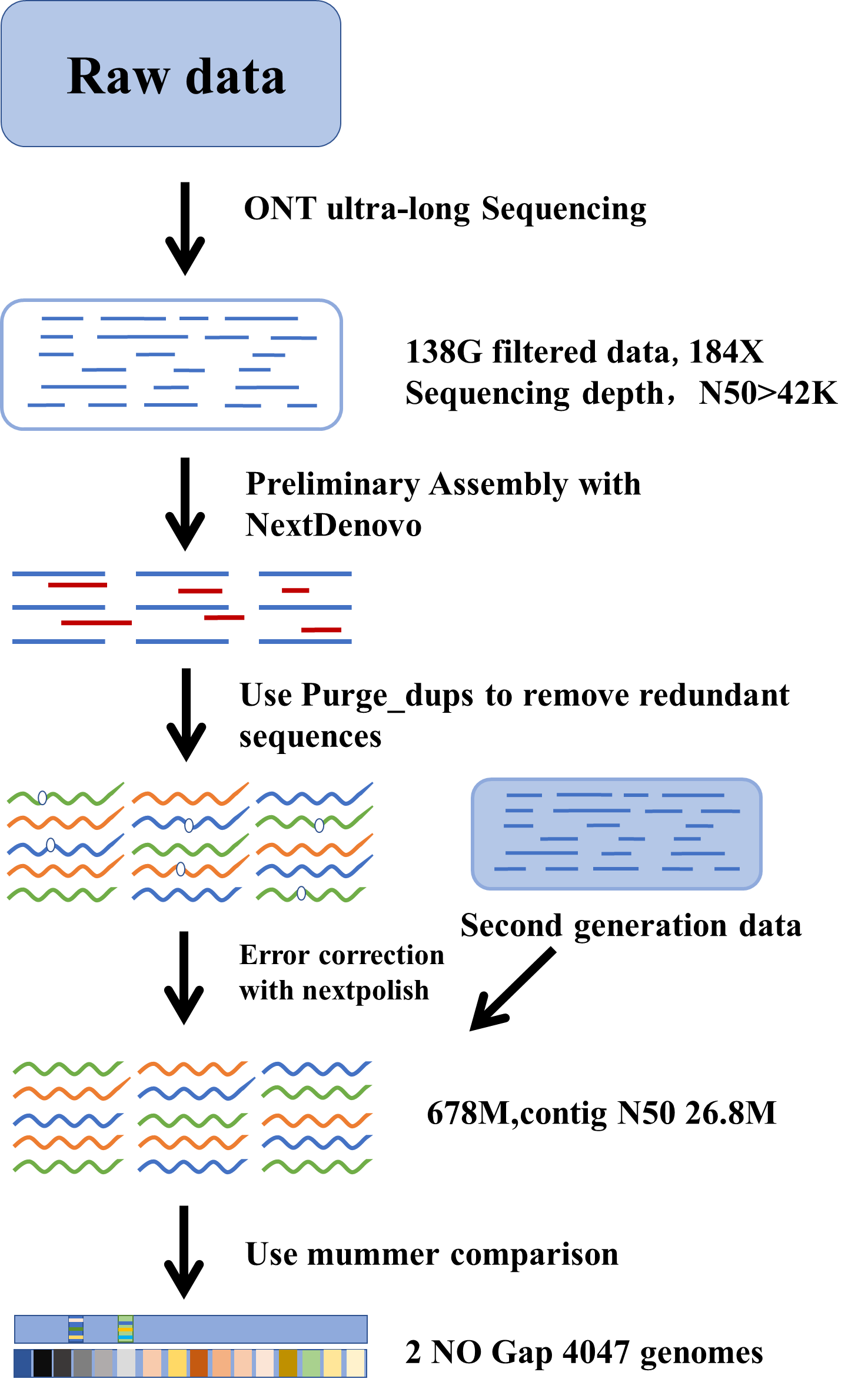

#### Fig. S5. Structure Variation (SV) calling strategy

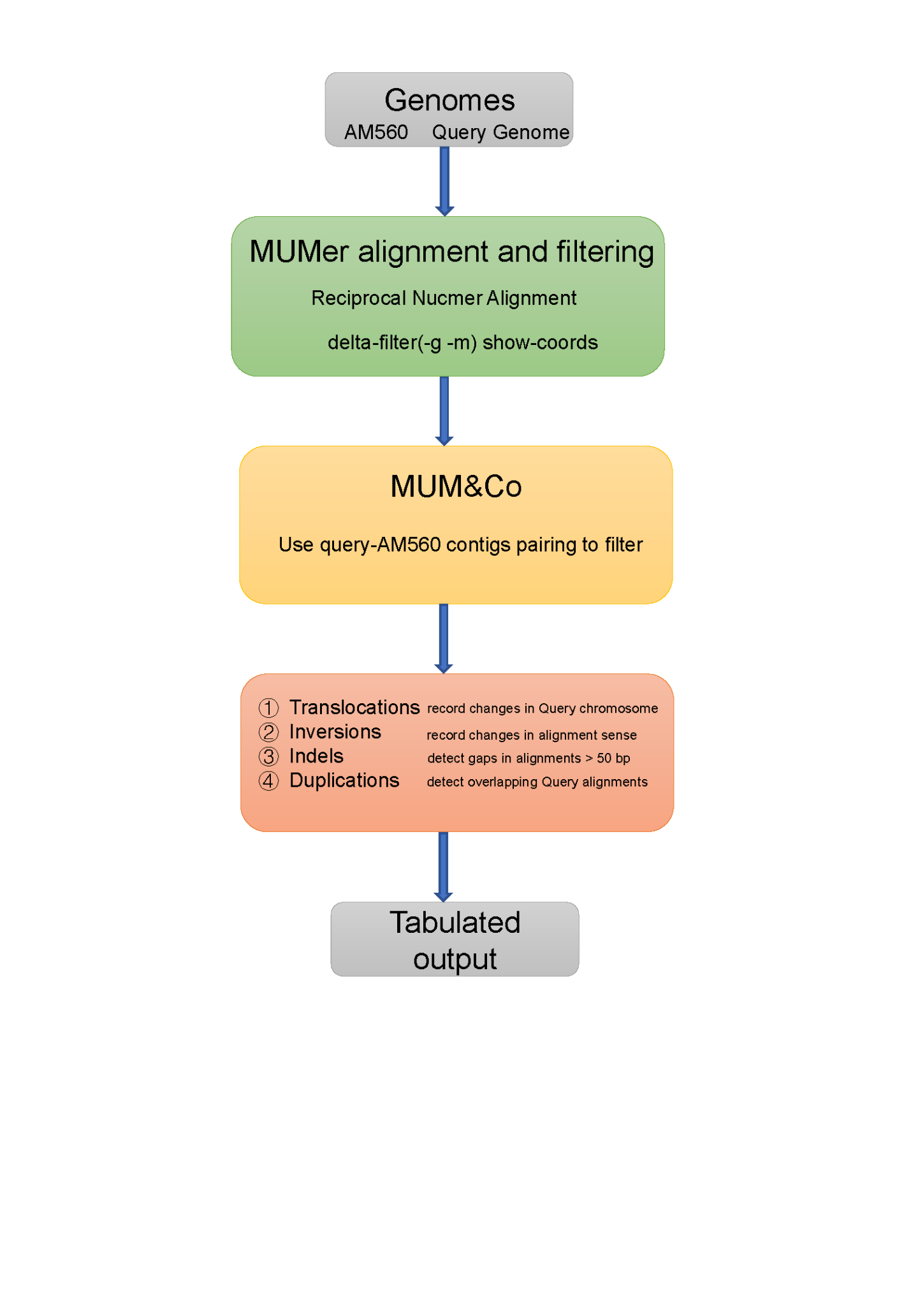

#### Fig. S6. The completeness assessment of the assembled cassava genome AM560, FLA4047 and W 14

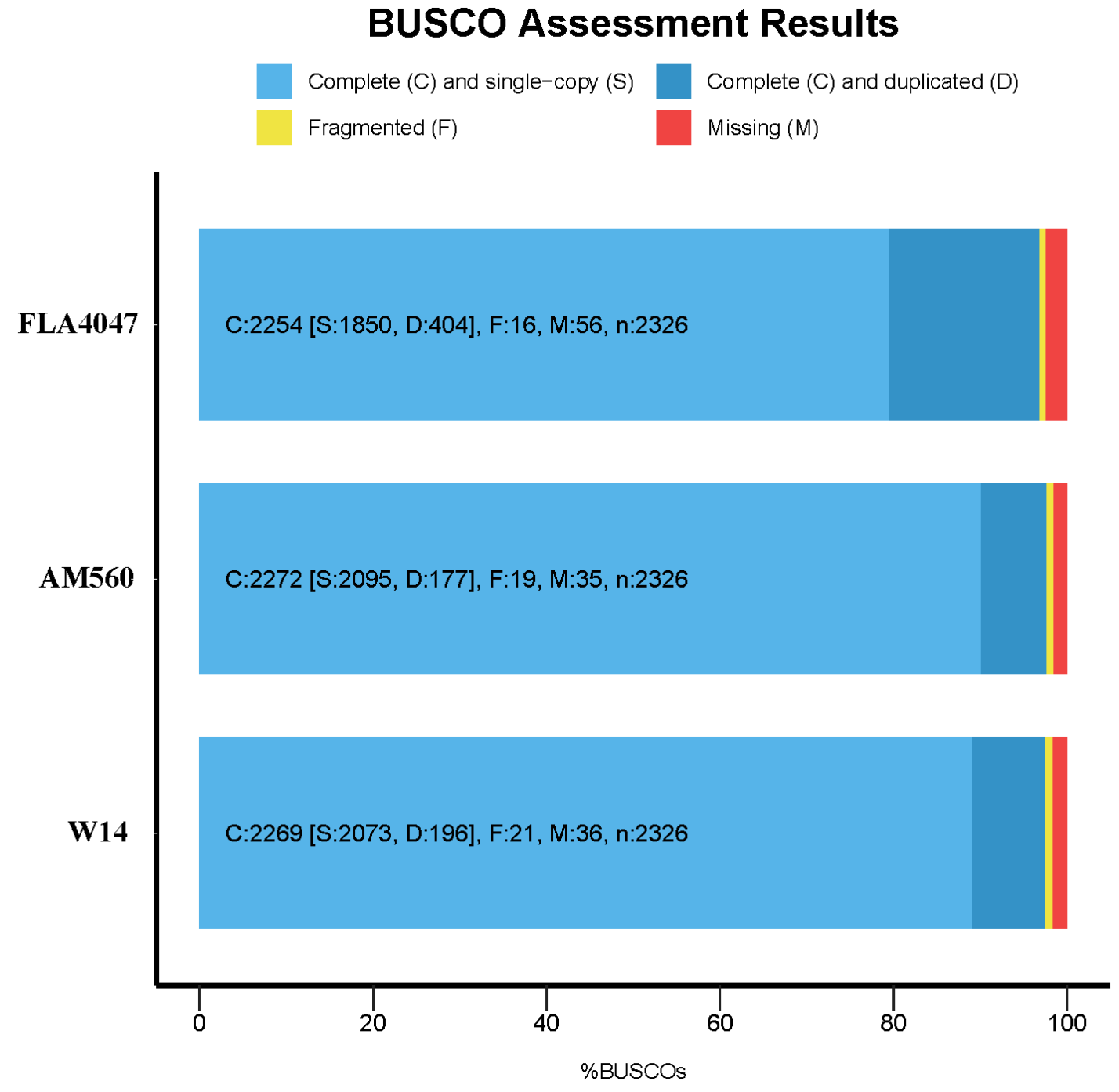

#### Fig. S7. The structure variation and co-linearity pairwise of genome AM560，FLA4047 and W 14 in chromosomal level

A. FLA4047 vs AM560

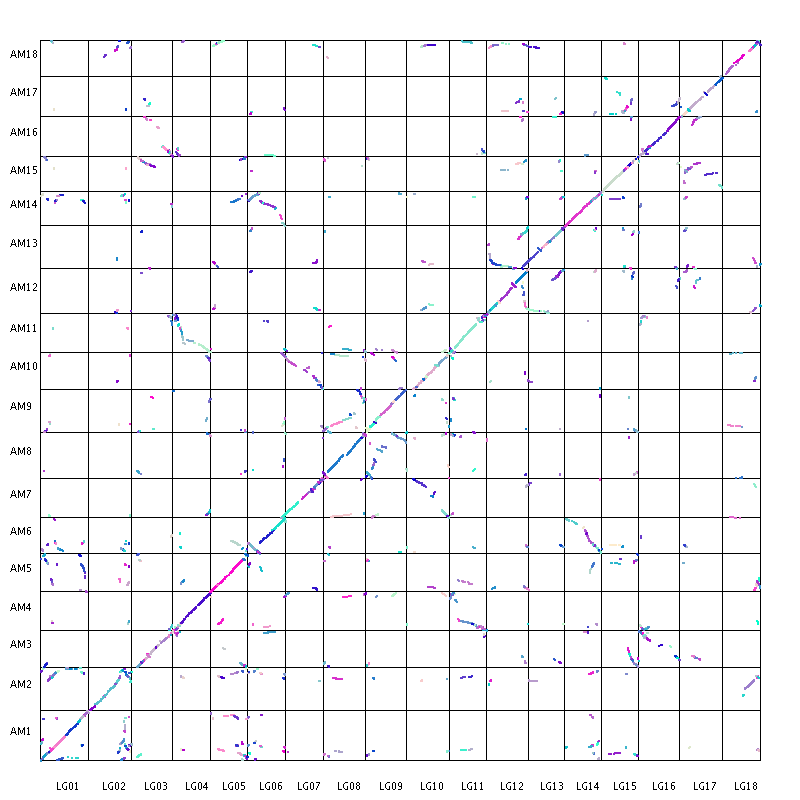

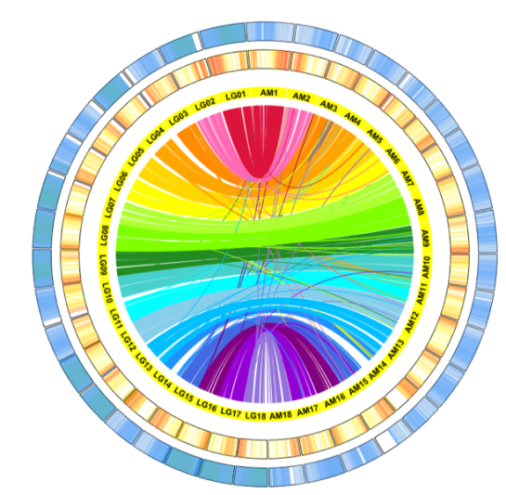

B. W14 vs FLA4047

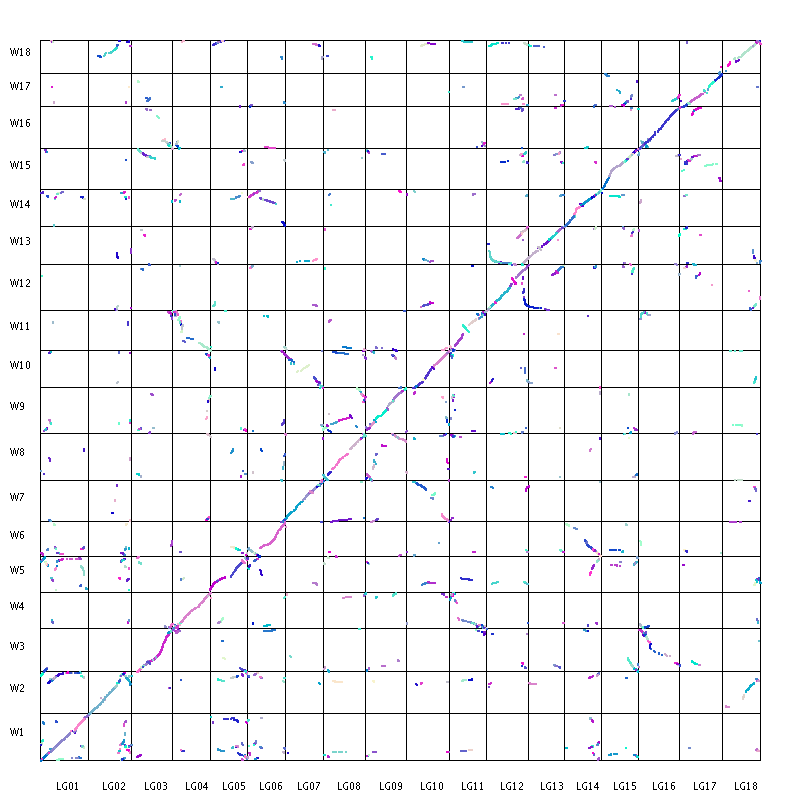

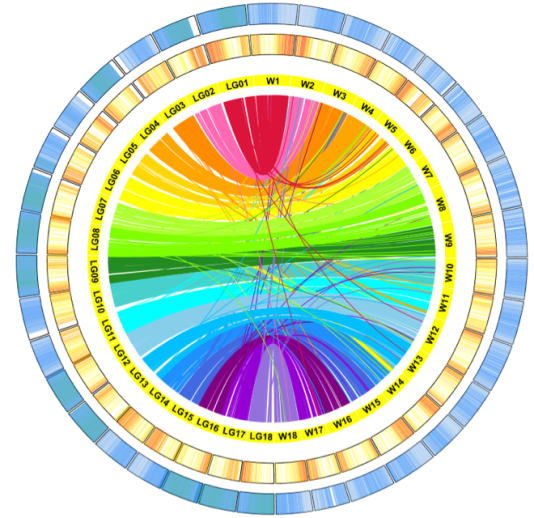

C. W14 vs AM560

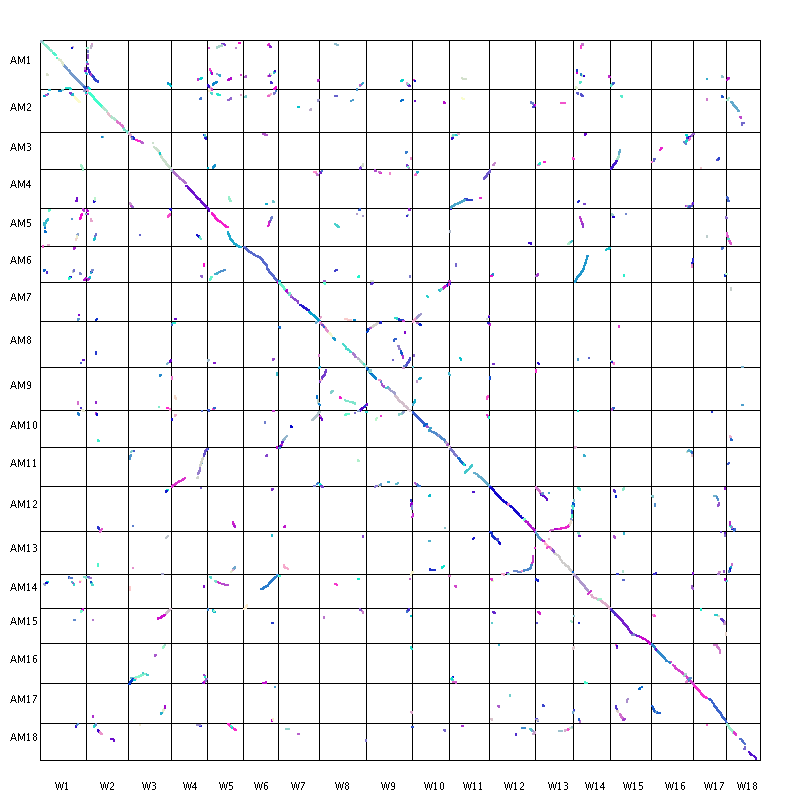

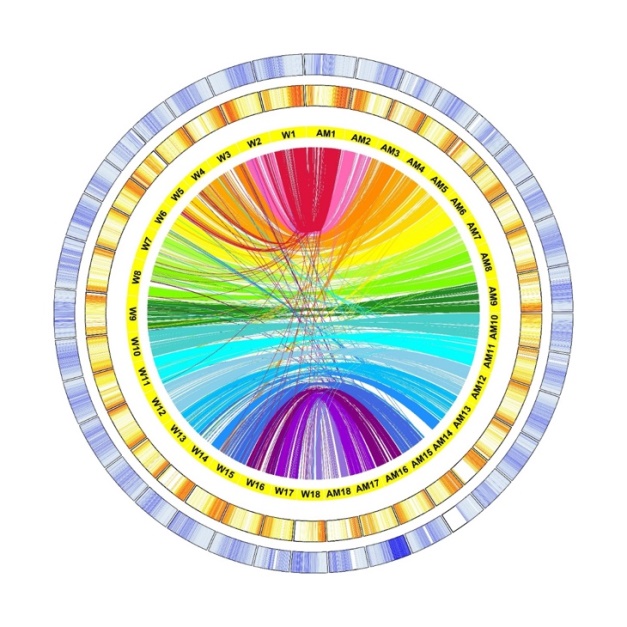

#### Fig. S8. Synteny and Structure Variation among genome of AM560, FLA4047 and W14

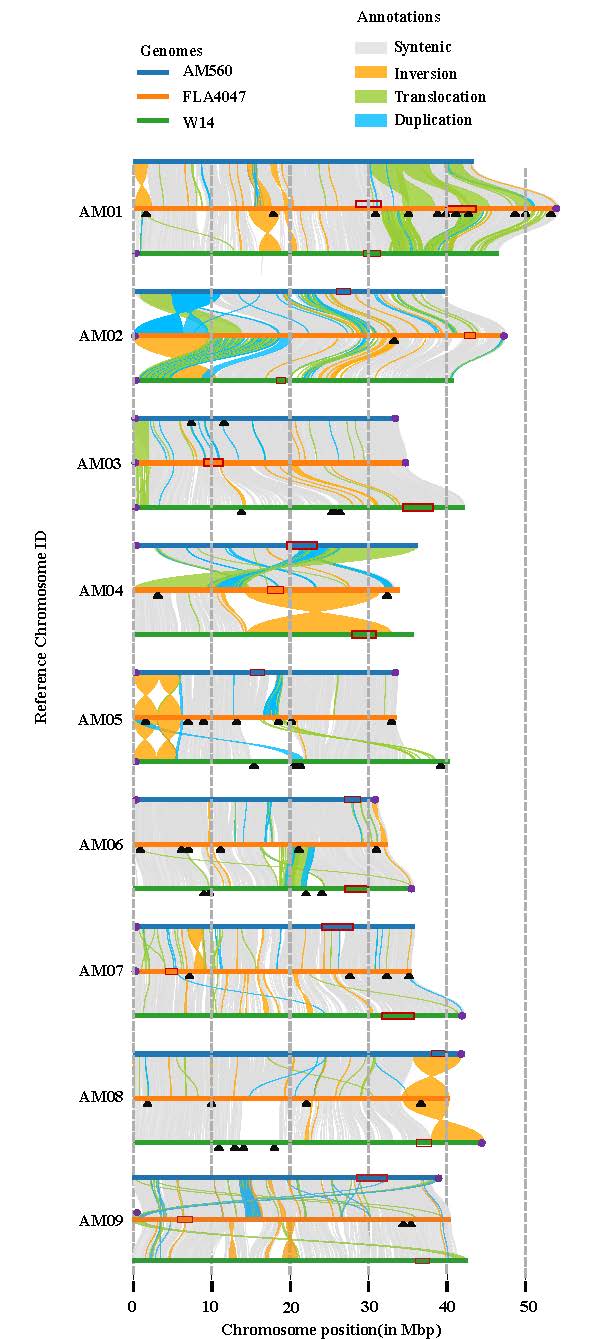

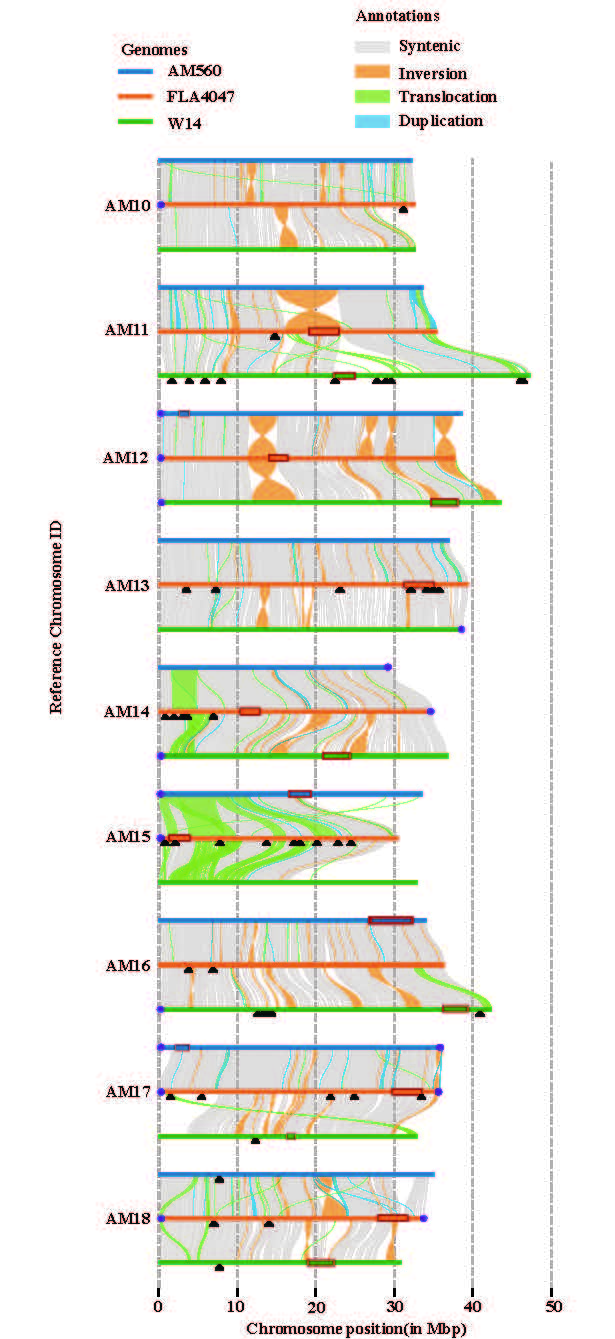

#### Fig. S9. The gene density map and distribution across the 18 chromosomes in AM560, FLA 4047 and W14

**AM560:**

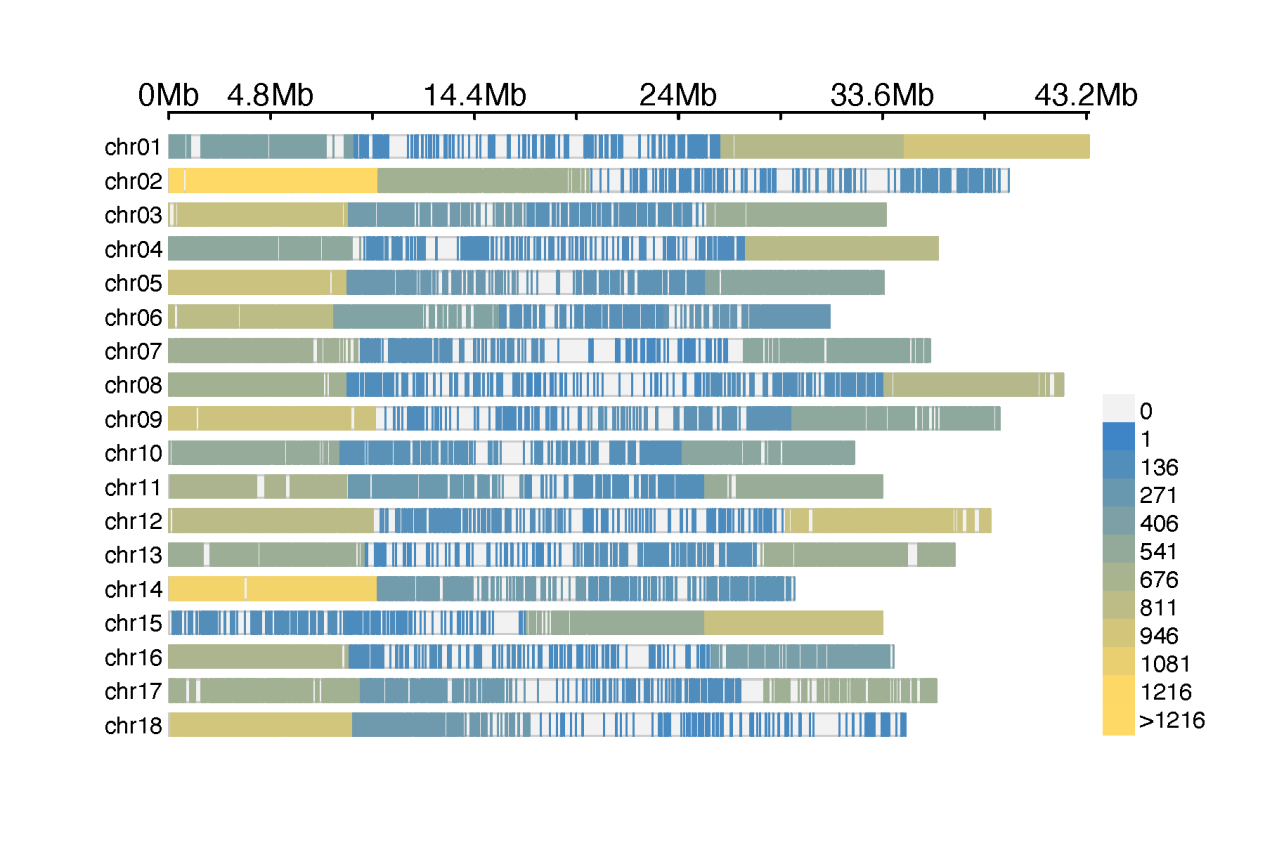

**FLA 4047：**

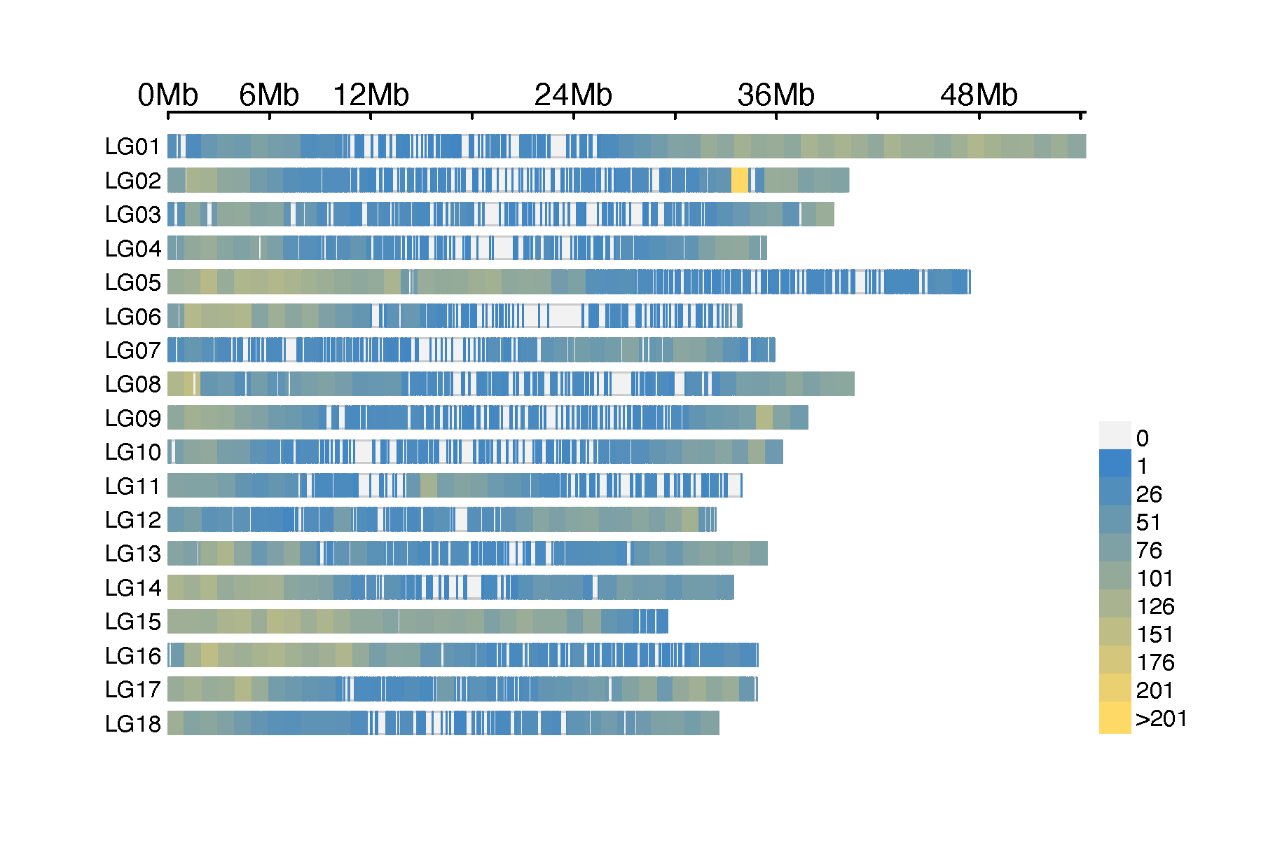

**W14:**

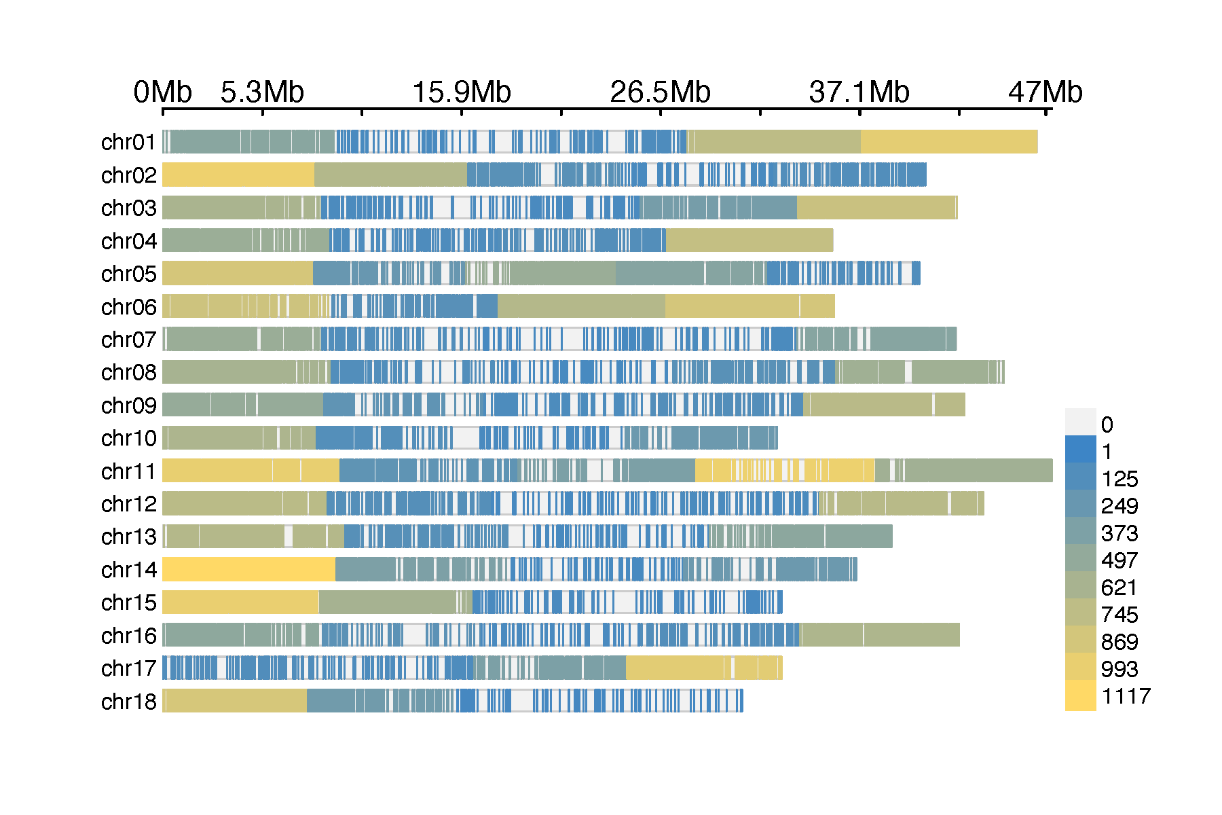
**Fig. S10. The common and specific genes annotated in genome of AM560, FLA4047 and W14**

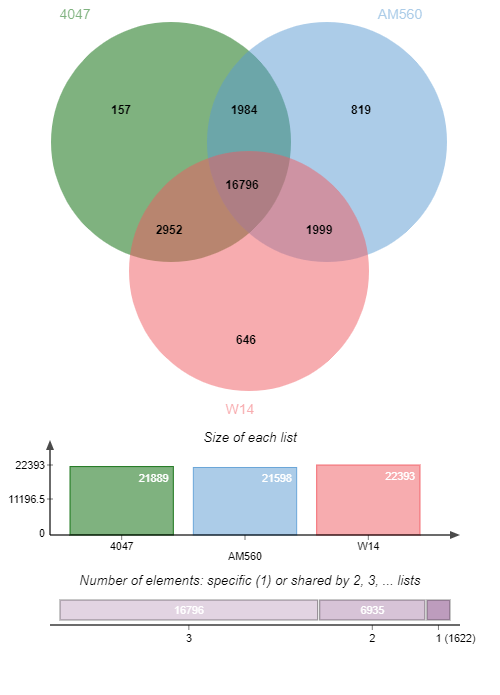

#### Fig. S11. Structural variations in pan genomes of cassava

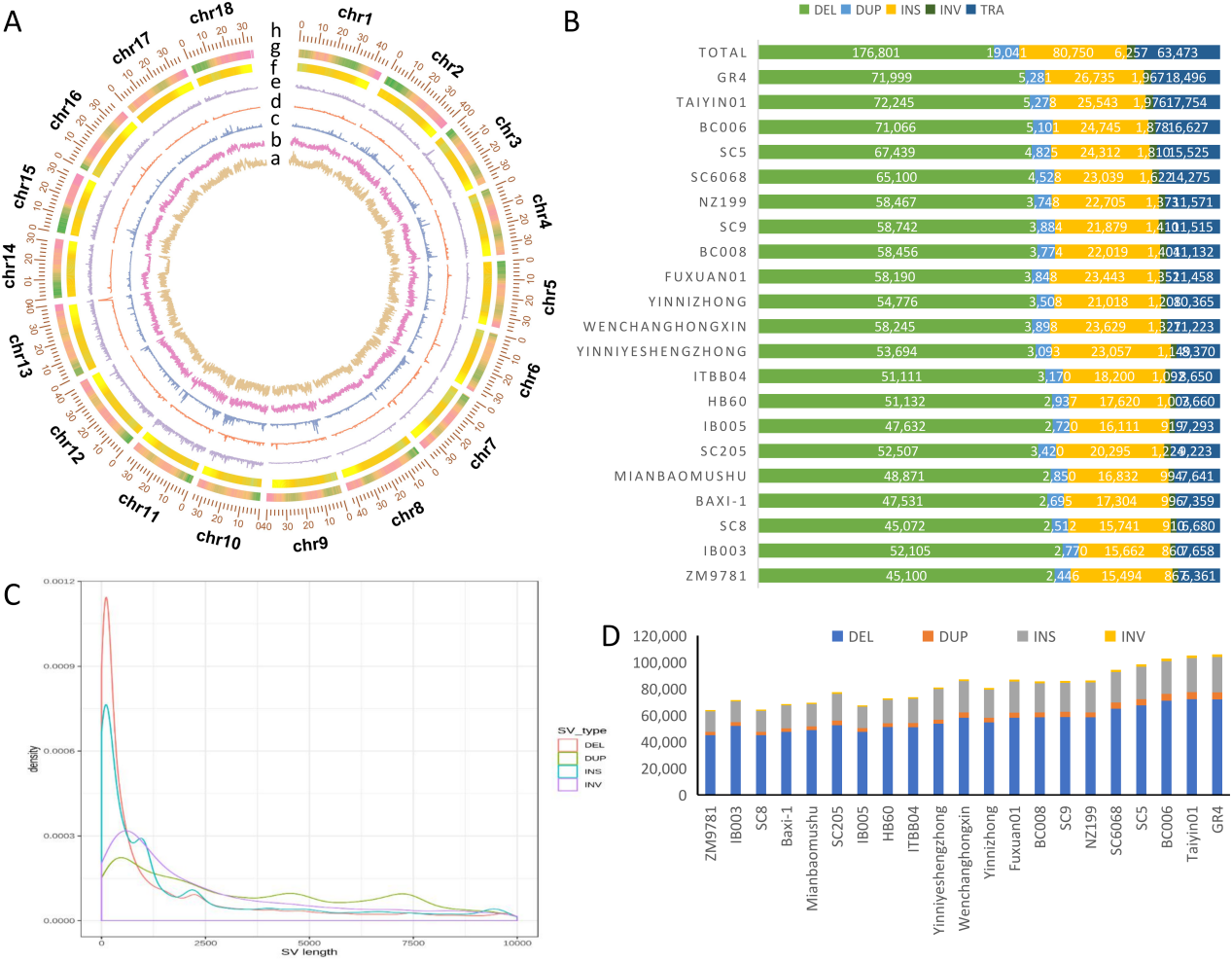

(A) Distribution of genetic variations from 21 genomes. (a) Insertions. (b) Deletions. (c) Duplications. (d) Inversions. (f) Translocations.

(B) SV in different varieties

(C) SV length distribution, The TRA type is the breakpoint of the swap, so no length is drawn

(D)The number of SV in different varieties

#### Fig. S12. Number and length of INS in Different Cassava Varieties

Above: Comparison of number and length of Insertions (INS) among 21 cassava genotypes; Down: INSs distribution of 21 cultivated clones and wild FLA4047 and W14

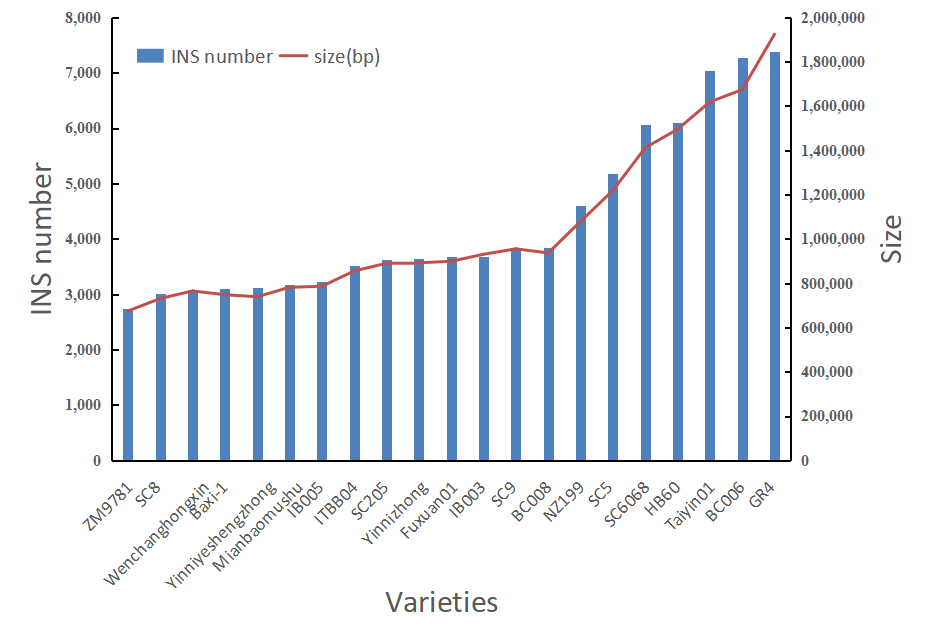

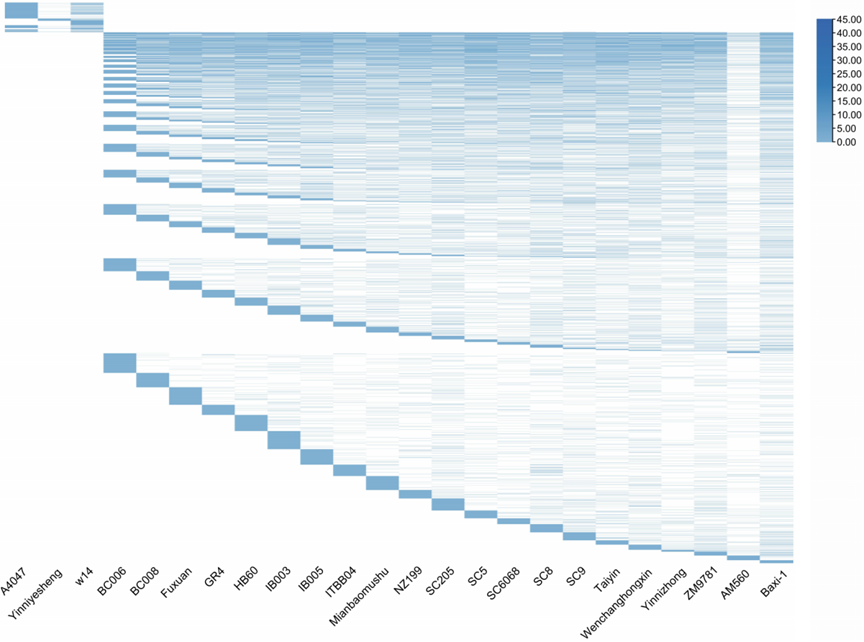

#### Fig. S13. KEGG gene enriched analysis of core and private function

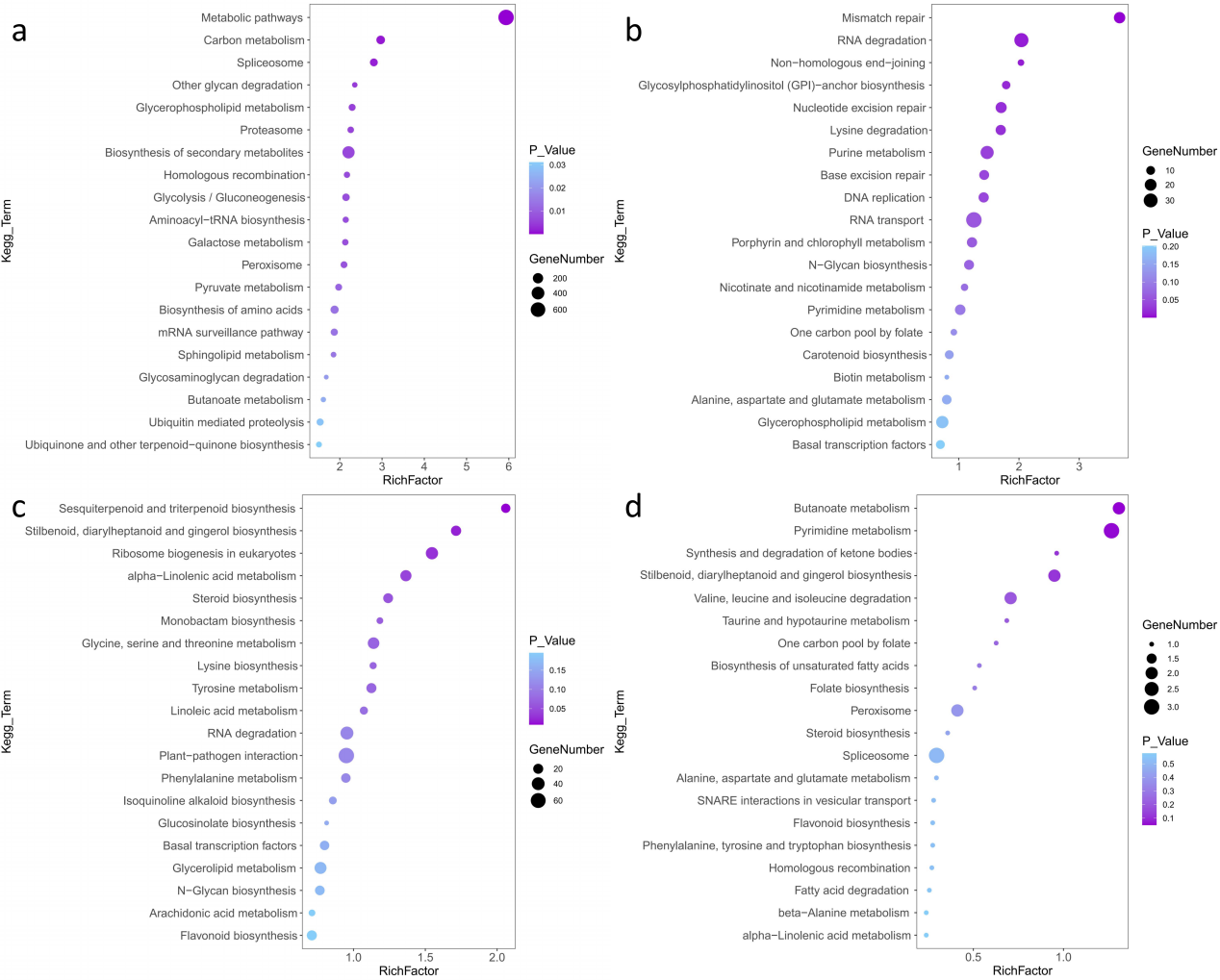

Notes: a Core genes；b Softcore genes；c Dispensable genes；d Private genes

#### Fig. S14. GO enrichment of genes only present in the genomes of FLA (A) and ME (B)

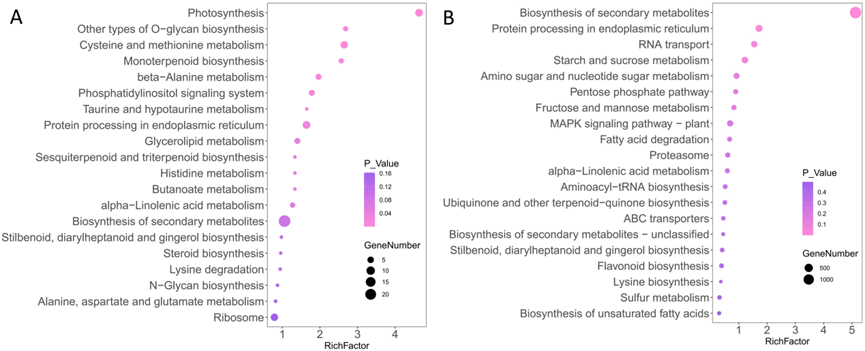

#### Fig. S15 Expansion and contraction of genes for photosynthesis of FLA and cultivated ME

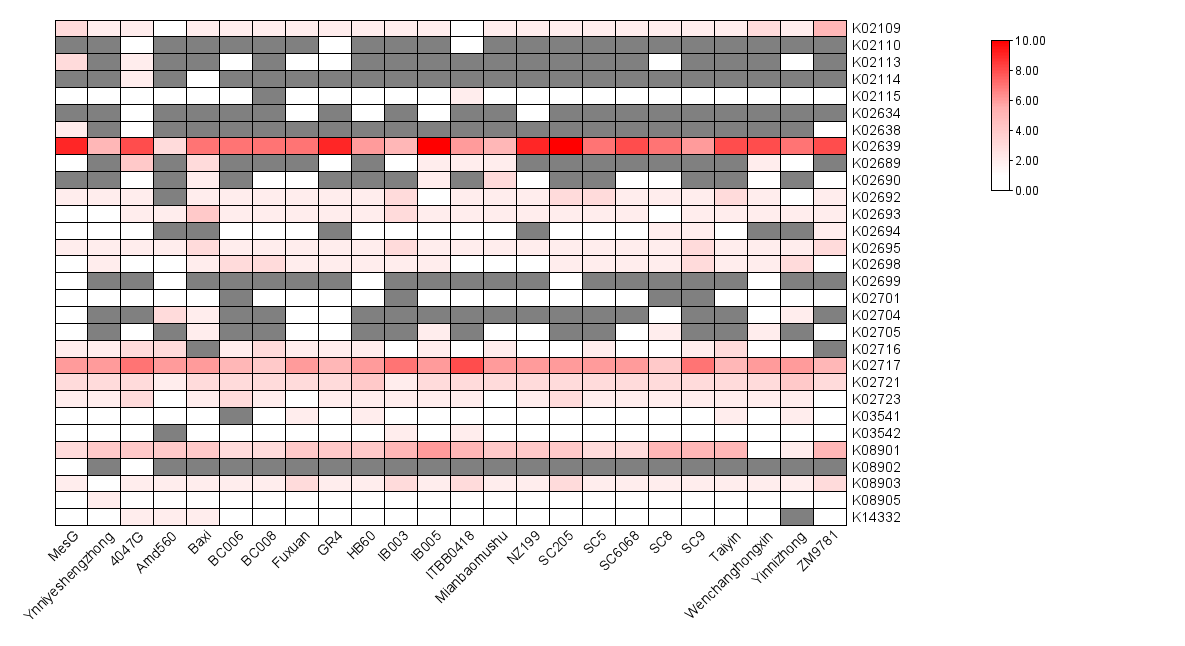

#### Fig. S16. KEGG showed expansion of genes for photoreaction pathway in cultivated cassava

(https://www.genome.jp/kegg-bin/show_pathway?mesc00195/mesc:6000019%09red/mesc:5999978%09red/mesc:6000014%09red)

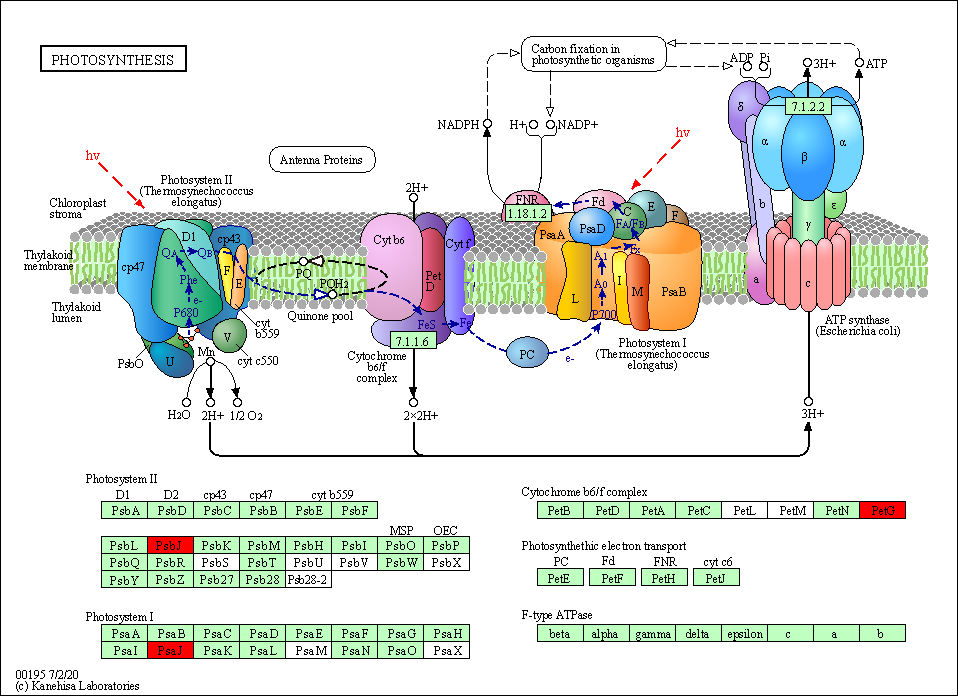

#### Fig. S17. Expansion and contraction of KEGG gene families involved in sucrose and starch metabolism in wild and cultivated cassava

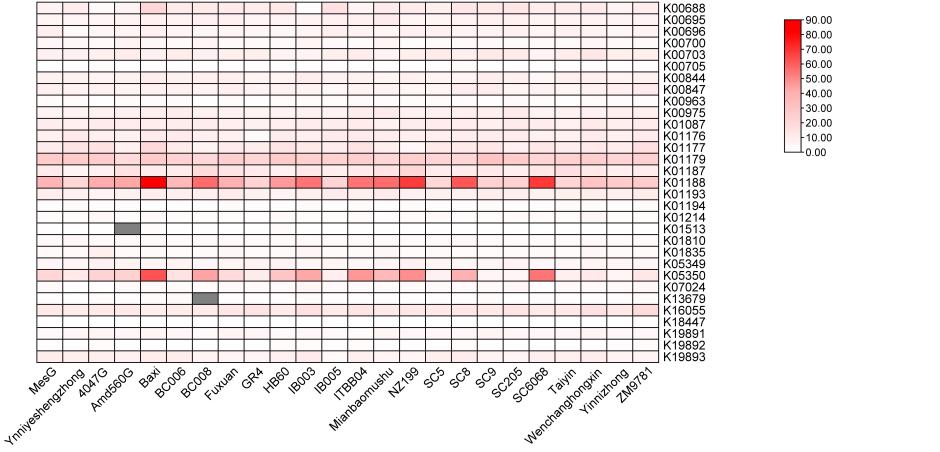

#### Fig. S18. Expansion of genes for starch and sucrose metabolic pathway in cultivated cassava

(https://www.genome.jp/kegg-bin/show_pathway?mesc00500/mesc:110606036%09red/mesc:110619165%09red/mesc:110615034%09red/mesc:110618773%09red/mesc:110607081%09red/mesc:110604864%09red/mesc:110610863%09red/mesc:110625210%09red/mesc:110618866%09red/mesc:110625212%09red/mesc:110601116%09red/mesc:110621386%09red/mesc:110605795%09red/mesc:110607072%09red/mesc:110612024%09red/mesc:110601324%09red/mesc:110600404%09red/mesc:110625211%09red/mesc:110618188%09red/mesc:110611765%09red/mesc:110618460%09red/mesc:110625145%09red/mesc:110625146%09red/mesc:110625147%09red/mesc:110603286%09red/mesc:110625143%09red/mesc:110611172%09red/mesc:110613770%09red/mesc:110602852%09red/mesc:110615840%09red/mesc:110602161%09red/mesc:110624372%09red/mesc:110630655%09red/mesc:110599614%09red/mesc:110628418%09red/mesc:110599718%09red/mesc:110602420%09red/mesc:110609378%09red/mesc:110627162%09red/mesc:110626256%09red/mesc:110624426%09red/mesc:110602151%09red/mesc:110605391%09red/mesc:110609238%09red/mesc:110614949%09red/mesc:110616004%09red/mesc:110607665%09red/mesc:110607664%09red/mesc:110617247%09red/mesc:110607661%09red/mesc:110607663%09red/mesc:110607662%09red/mesc:110604394%09red/mesc:110612297%09red/mesc:110607668%09red/mesc:110618520%09red/mesc:110619584%09red)

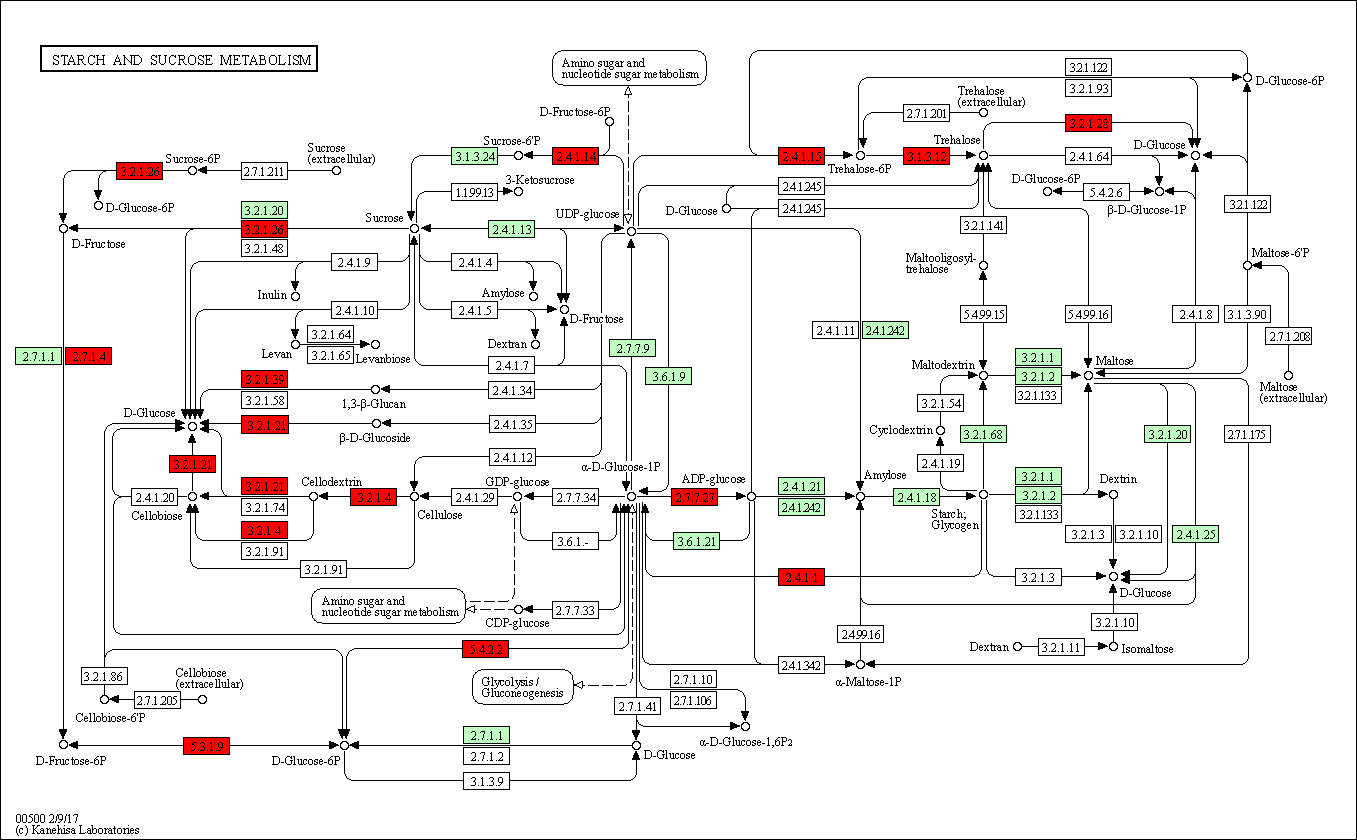

#### Fig. S19. Expansion and contraction of genes for phytohormone pathway in wild and cultivated cassava

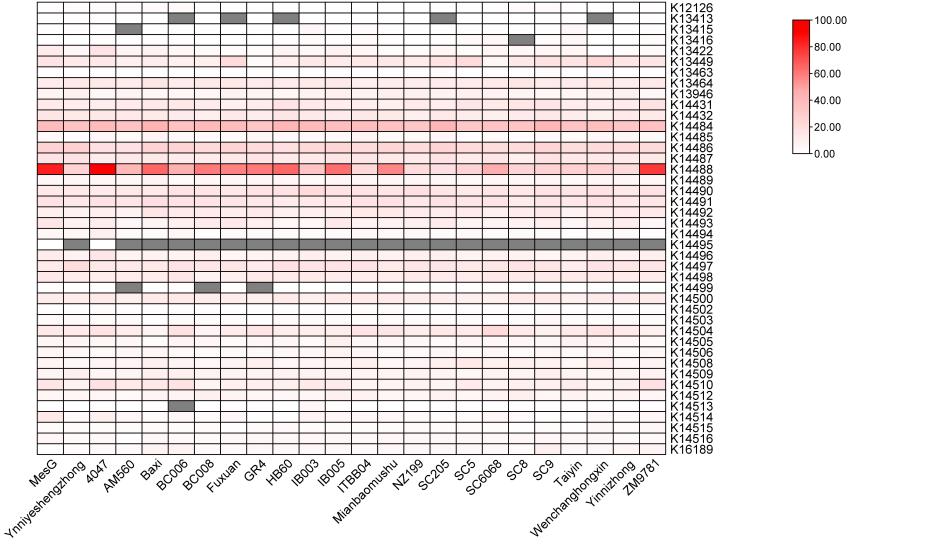

#### Fig. S20. Enrichment of gene families of important transcription factors

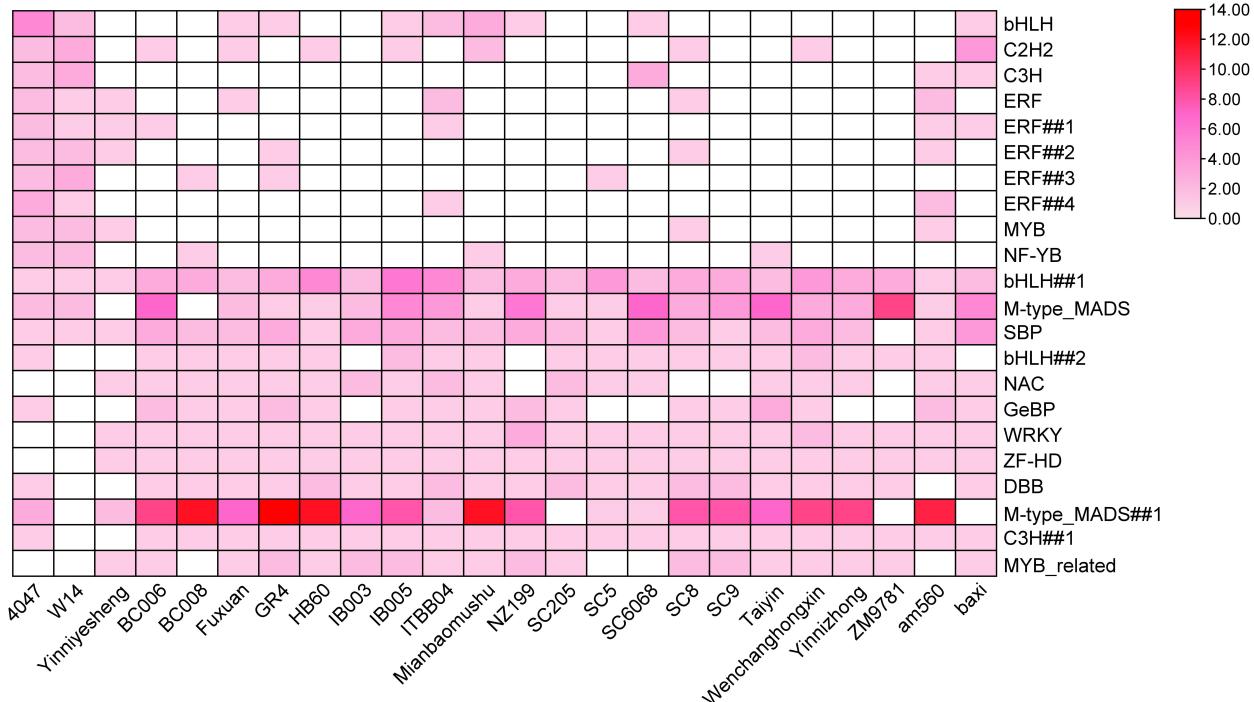

#### Fig. S21. The distribution of SNPs and InDels in cultivated cassava ME and wild progenitor subspecies FLA

SNPs in cassava and FLA populations

SNP numbers

| regions in genome | cassava | flabellifolia |
| --- | --- | --- |
| Total: | 13735173 | 22032832 |
| gene region | 2891925 | 3679623 |
| UTR | 189432 | 251873 |
| CDS | 614140 | 944295 |
| intron | 2088353 | 2483455 |
| Non-gene region | 10843248 | 18353209 |

InDels in cassava and FLA populations

InDel number

| regions in genome | cassava | flabellifolia |
| --- | --- | --- |
| Total: | 2719151 | 4533604 |
| gene region | 434384 | 681160 |
| UTR | 40407 | 67259 |
| CDS | 25213 | 45949 |
| intron | 368764 | 567952 |
| Non-gene region | 2284767 | 3852444 |

#### Fig. S22. Selective Sweeping from Wild Progenitors（FLA）to Cultivars （ME）with Fst and ROD paramater The red broken line is a significant region containing the *MeTFL2* gene, and domestication-related signals are detected in both Fst and ROD detection

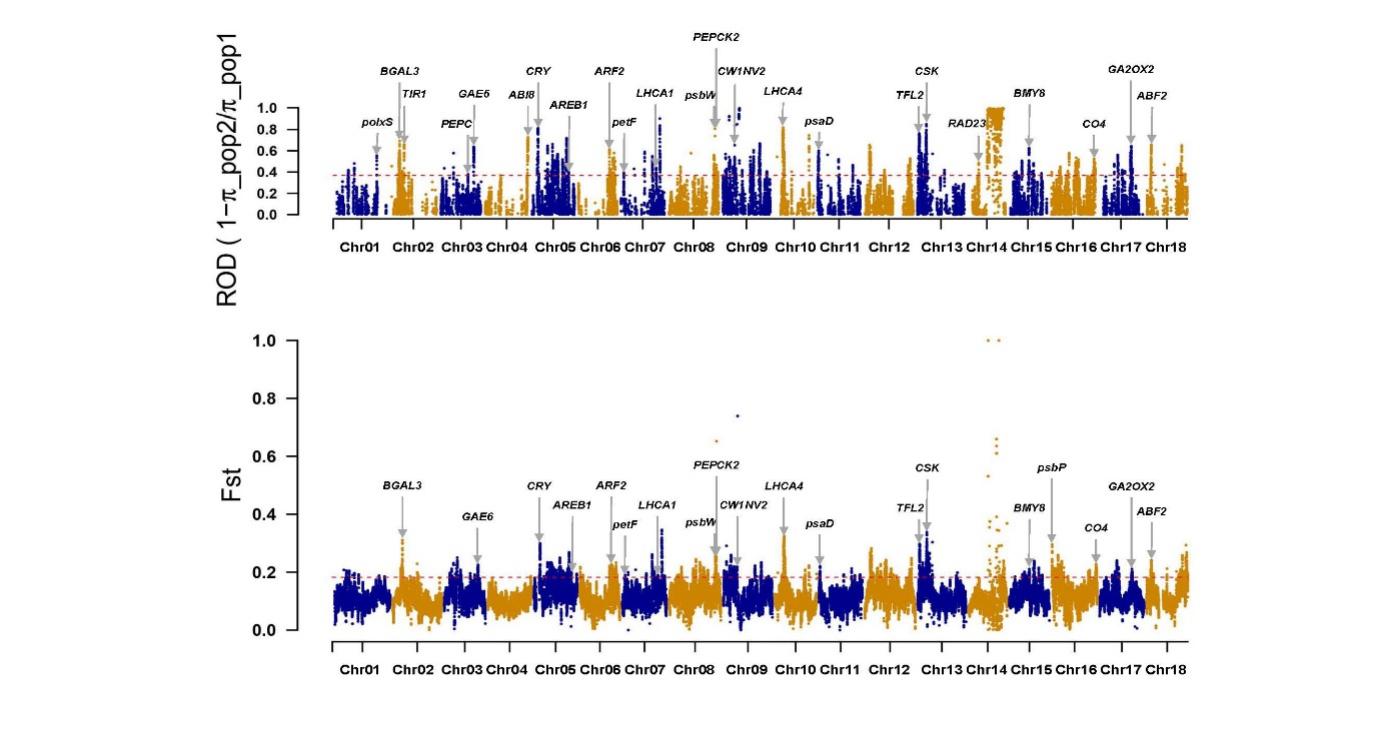

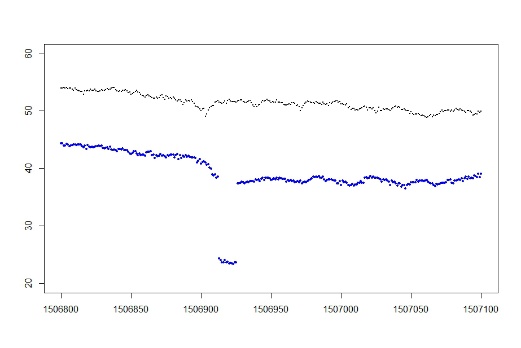

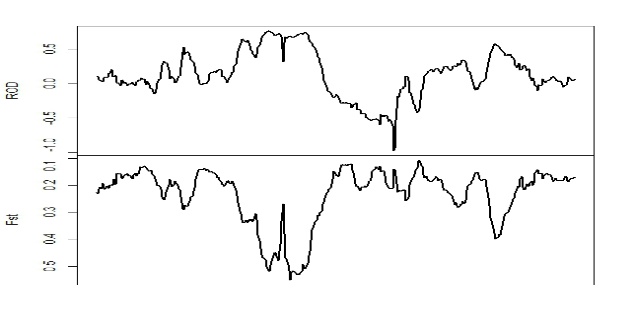

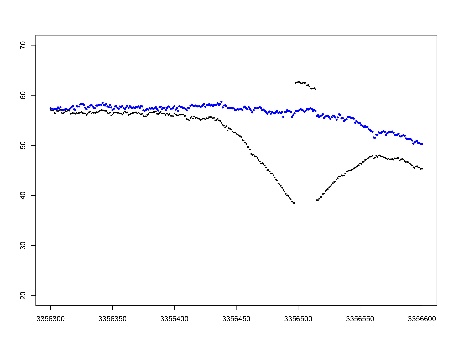

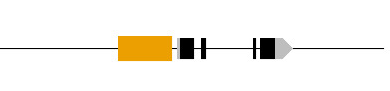

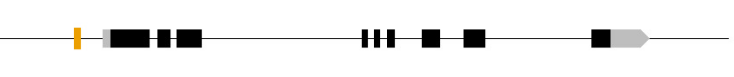

*TFL2 (13G011900)*

insert	fla 16/27

cassava 1/111

*CSK (13G022900)*

SINE fla 1/28

cassava 1153/198

Mean Reseq reads coverage

cassava

fla

fla

cassava

#### Fig. S23. GO and KEGG terms of the genes presented in selective sweeping regions

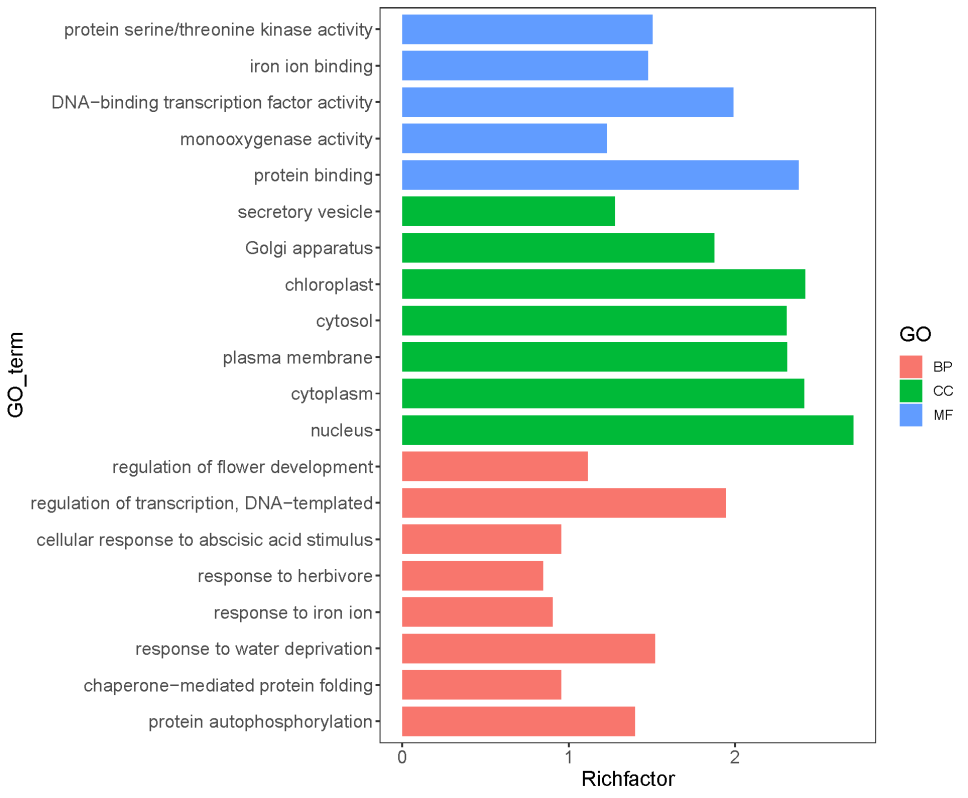

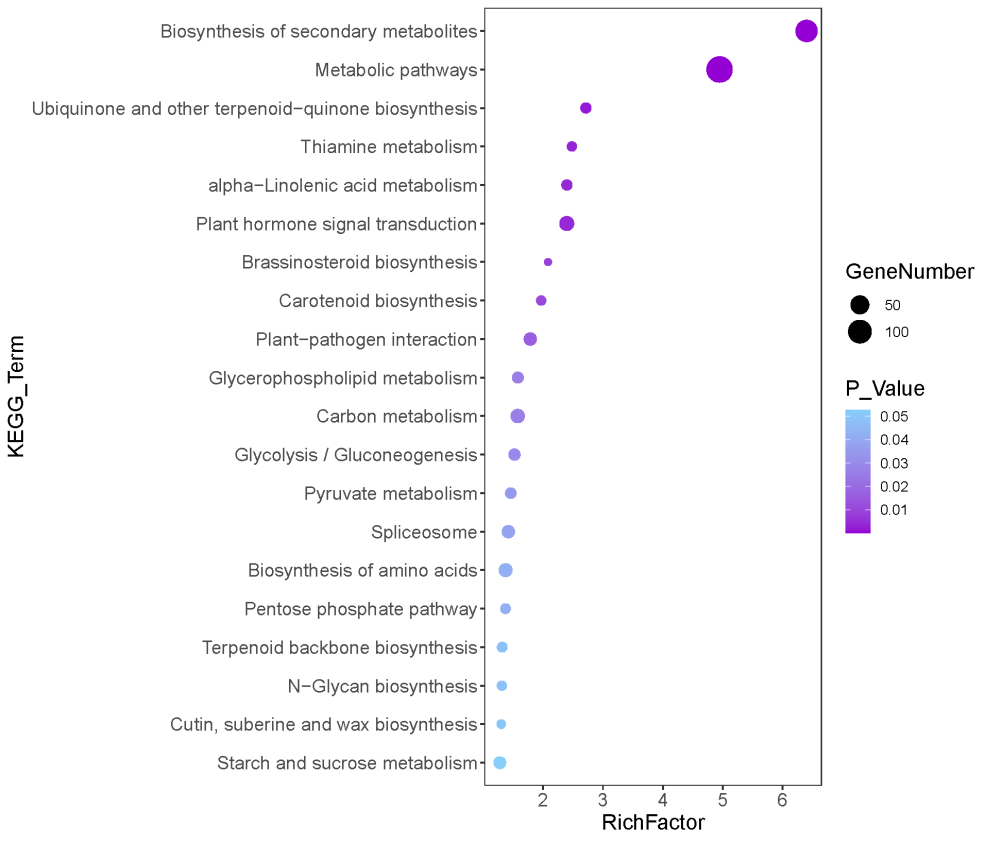

#### Fig. S24. The transcriptomic expression profiling of genes for photo reaction (left hotmap) and carbon assimilation (right hotmap) in photosynthesis of cassava plants DAP120 in the field Cheng Mai. Hainan island

Comparison of FLA4047, W14 and GLAyn, the three wild relatives with 19 cultivated varieties, *MeCSK, MeFNR2, MeFNR3, MeCytb6f* and *MeATPase* that genes involved in photochemical conversion and electron transport higher expressed in cultivated clones than in wild ancestors (left hotmap); But only *MePPDK, MePEPC* and *MeNADP-Me*, the genes of C4 enzymes high expressed in cultivars than in the wild accessions (right hotmap).

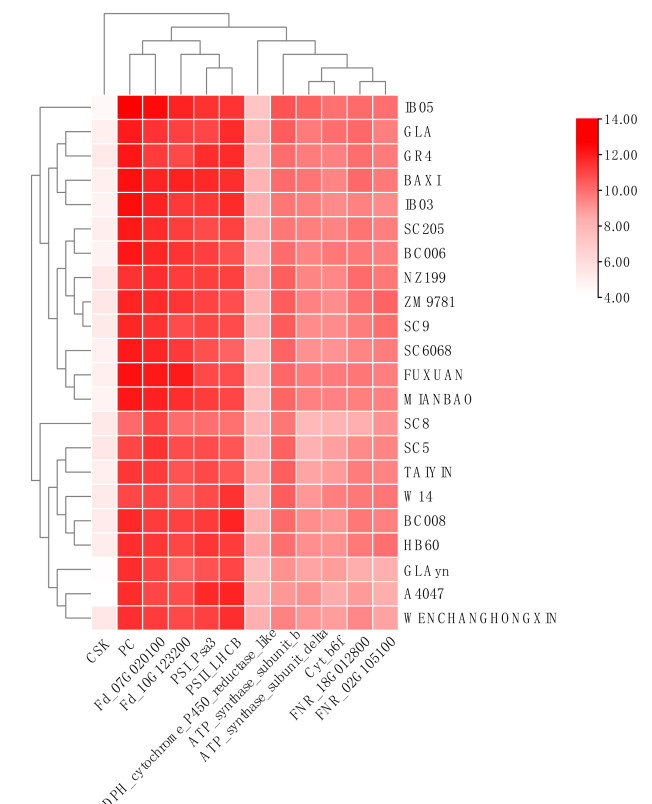

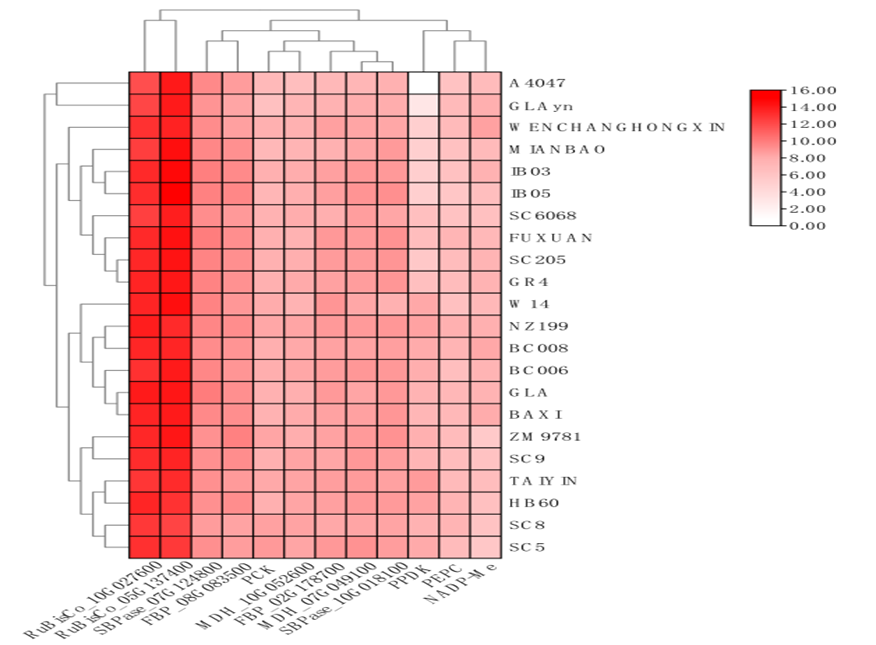

#### Fig. S25. The transcriptomic expression profiling of genes for cyanide metabolism in leaves and storage root (R) of plants DAP 120 with 22 accessions include of sweet and bitter clones in field of Cheng Mai, Hainan island

Comparison of a dozen varieties with low cyanide content in storage root with another dozen accessions include of wild ancestor with high cyanide content: Among the total of 22 genes, *MeCYP79D2, MeCYP79D1, MeUGT85 K5*and *MeCYP71E7* involved in biosynthesis, *MeGTR* and *MeCGRT1,*the transporter of cyanide, and the *MeHNL10-like* and *MeHNL24-like* responsible for the decomposition of cyanoside higher expressed in leaves of clones with high cyanide; But only *MeCYP79D2, MeCYP79D1, MeMATE1 and MeHNL10-like* high expressed in storage root of bitter varieties but not sweet varieties (upper hotmap). In a set of sweet clones, genes for cyanide transport, *MeGTR，MeCGTR1, MeCGTR2*, and decomposition like *MeHNL10-like*，*MeHPP*，*MeBCASb* and *MeH-ATPase* significantly down-regulated in storage roots(middle hot-map). In other bitter group include of the wild ancestor, only *MeGTR，MeCGTR1* , *MeHNL10-like*，*MeHPP* and *MeBCASb* have been selectively low expressed in the storage roots (down hot-map).

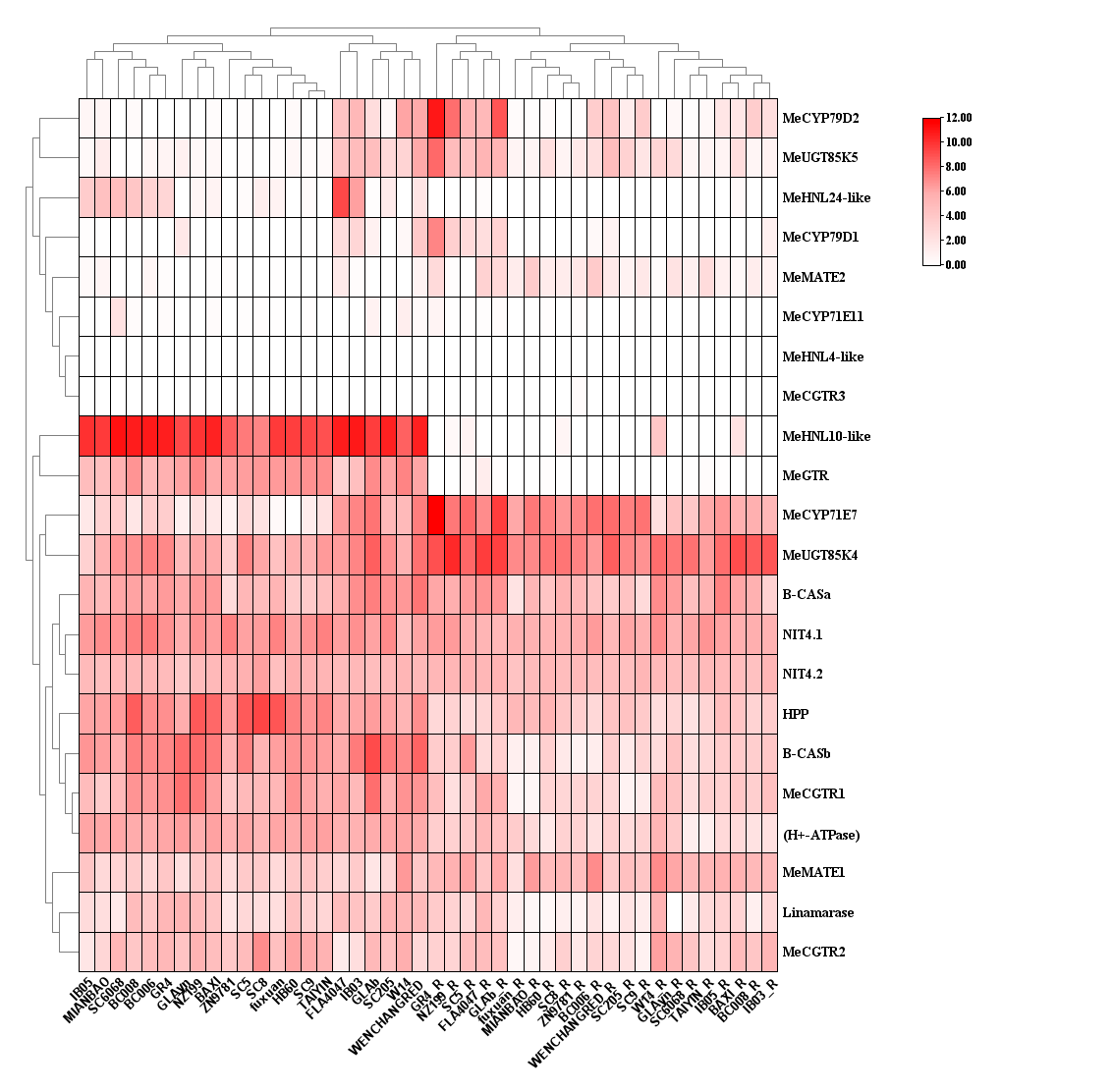

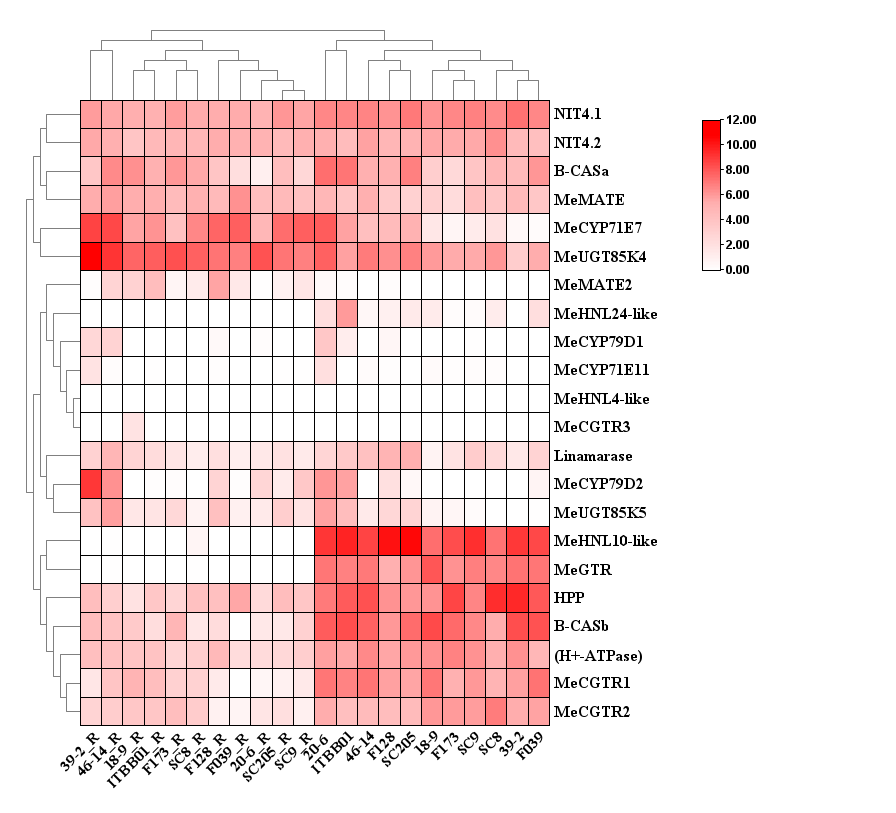

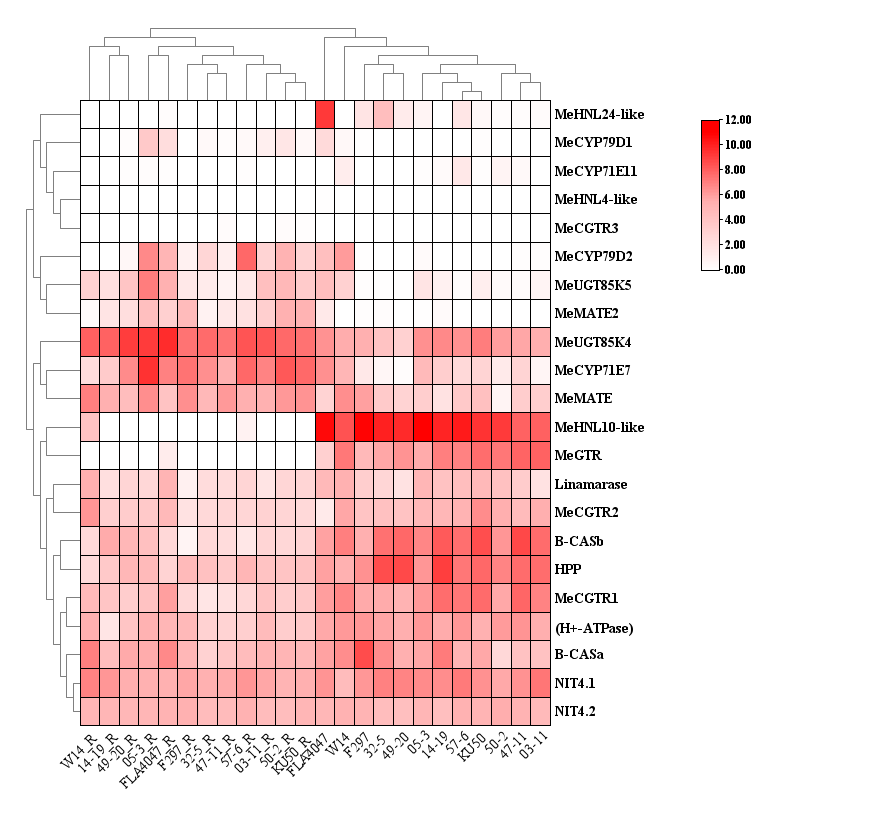

#### Fig. S26. Differential expression of genes in photosynthesis pathway between wild ancestors to cultivars validated by Real Time Q-PCR

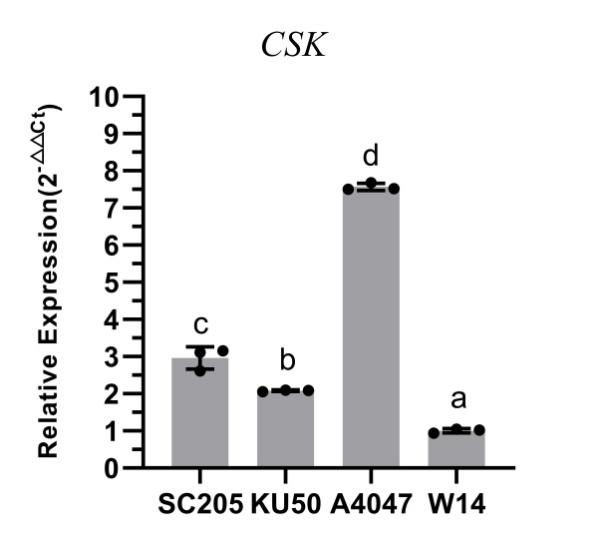

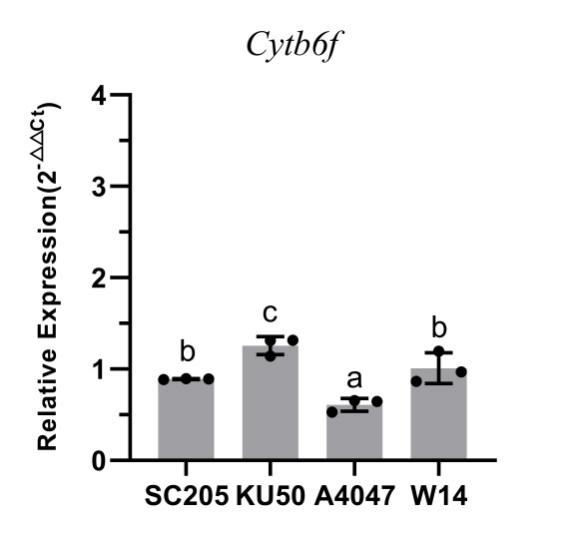

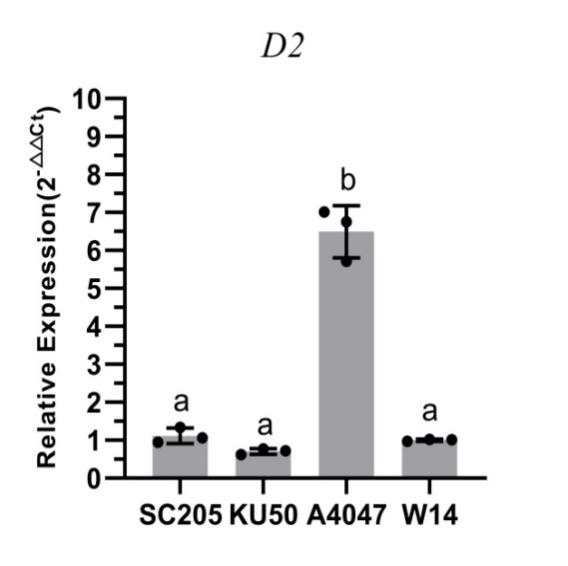

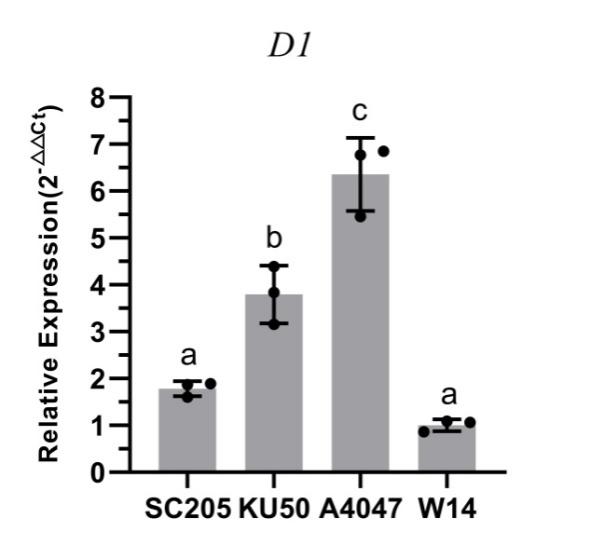

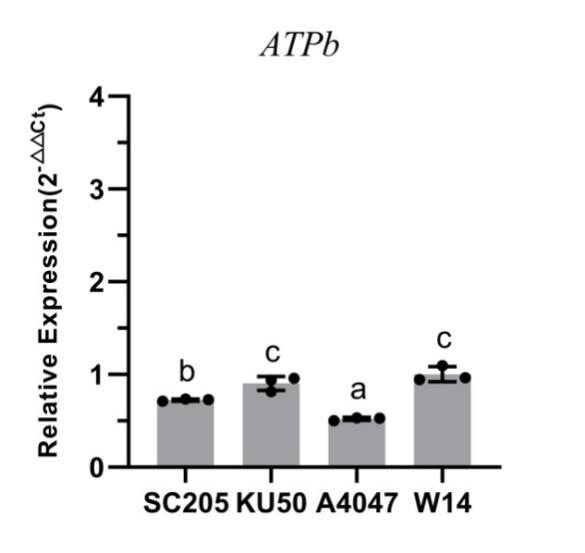

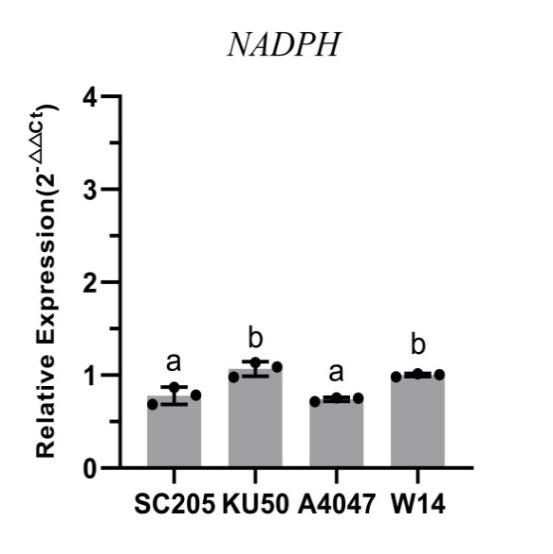

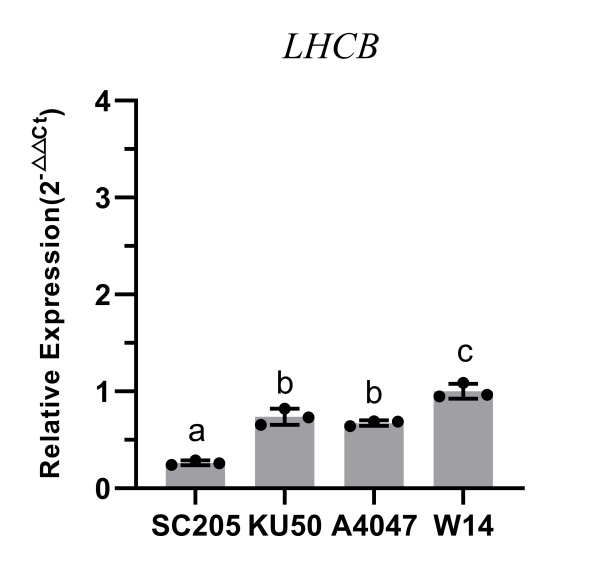

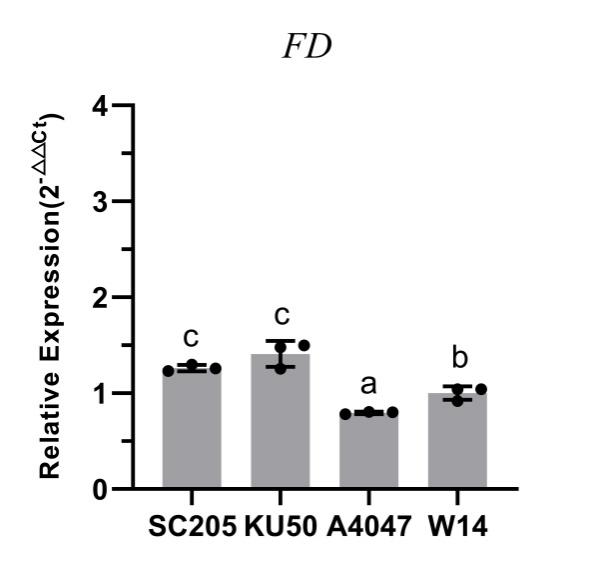

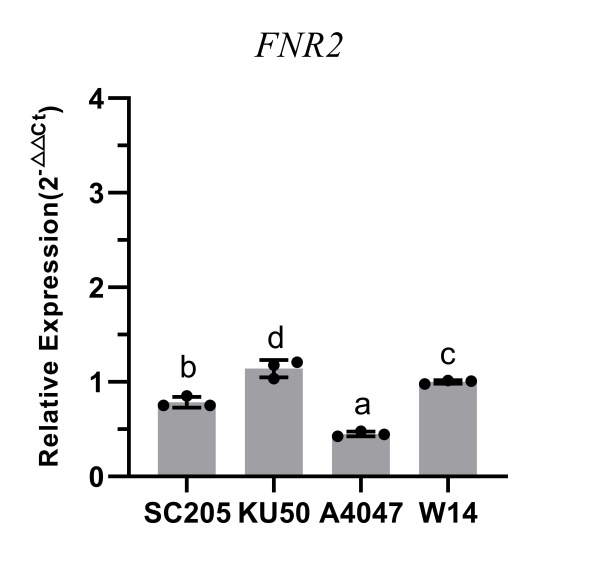

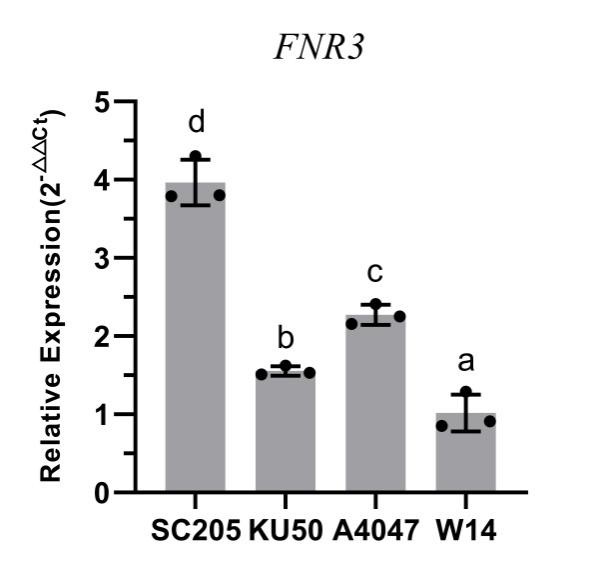

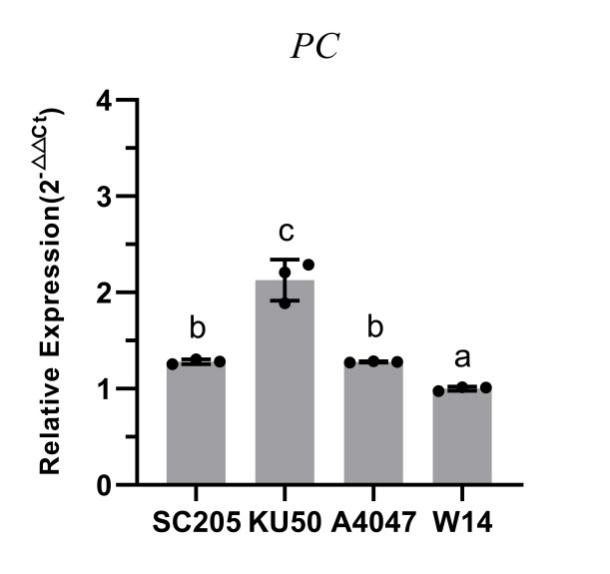

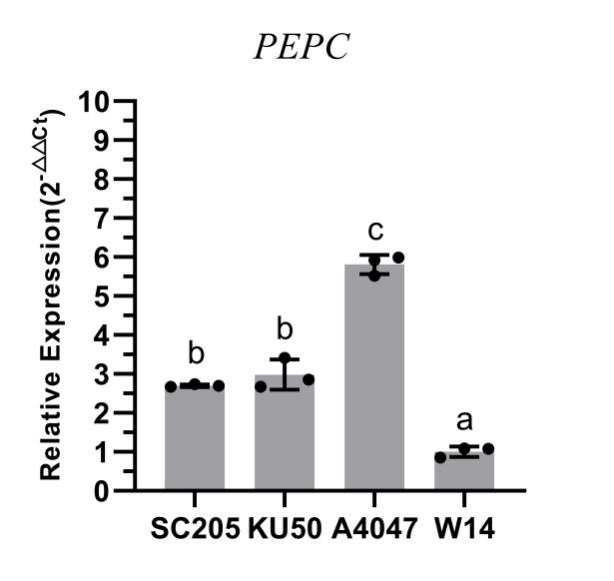

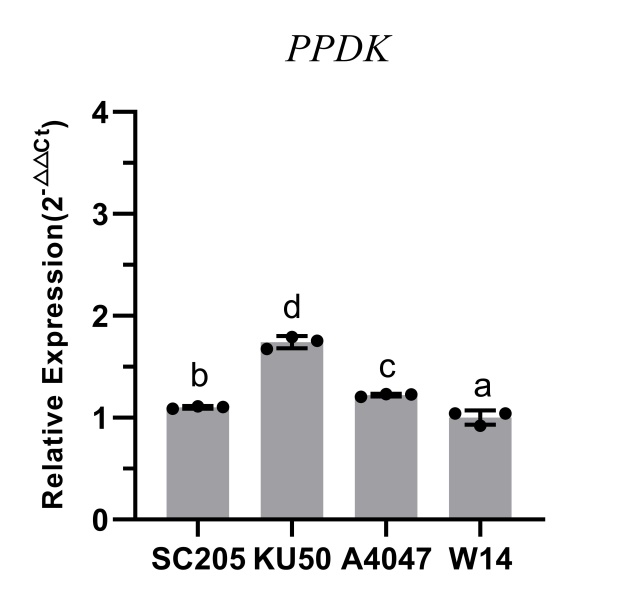

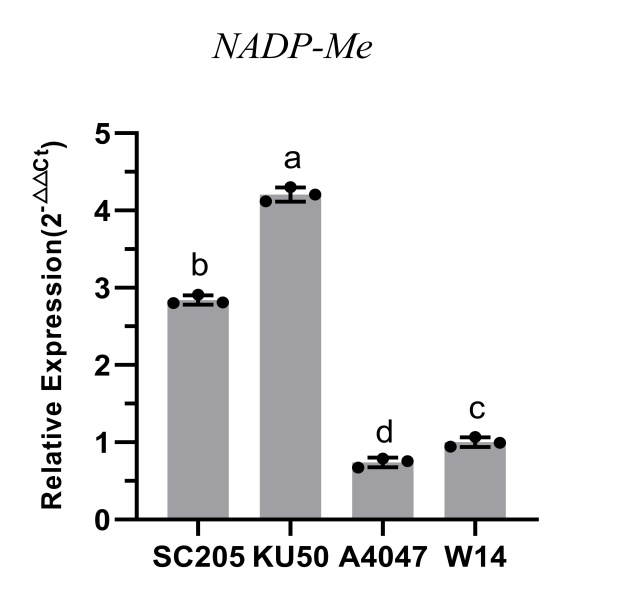

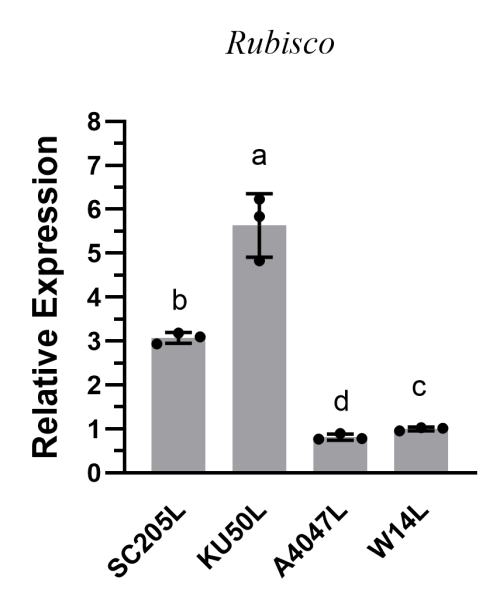

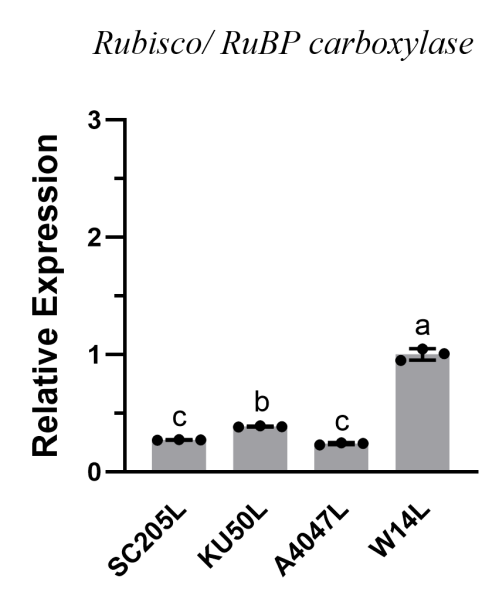

#### Fig. S27. Differential expression of genes involved in formation of floral primordia and storage root validated by Real time Q-PCR

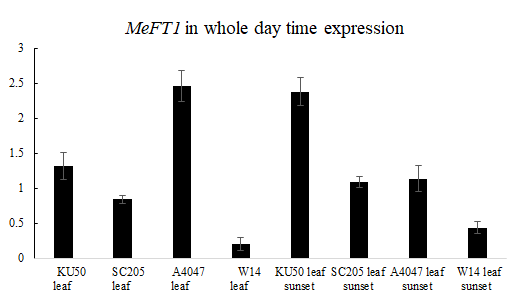

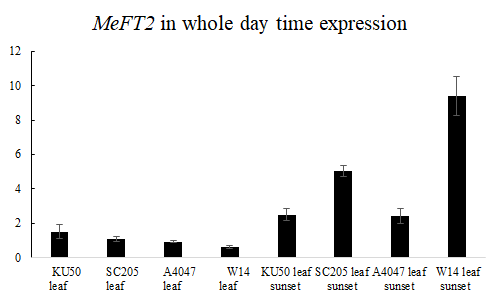

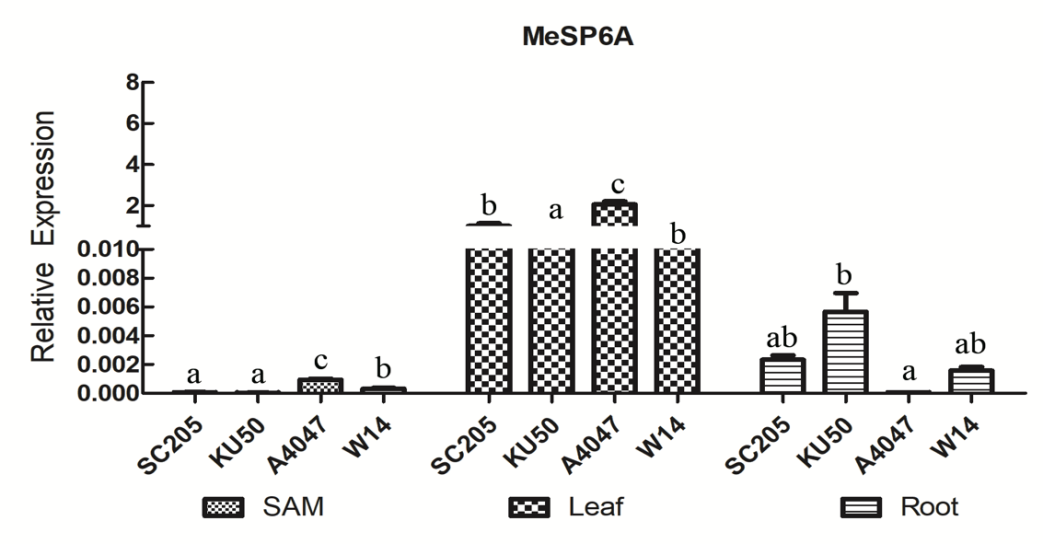

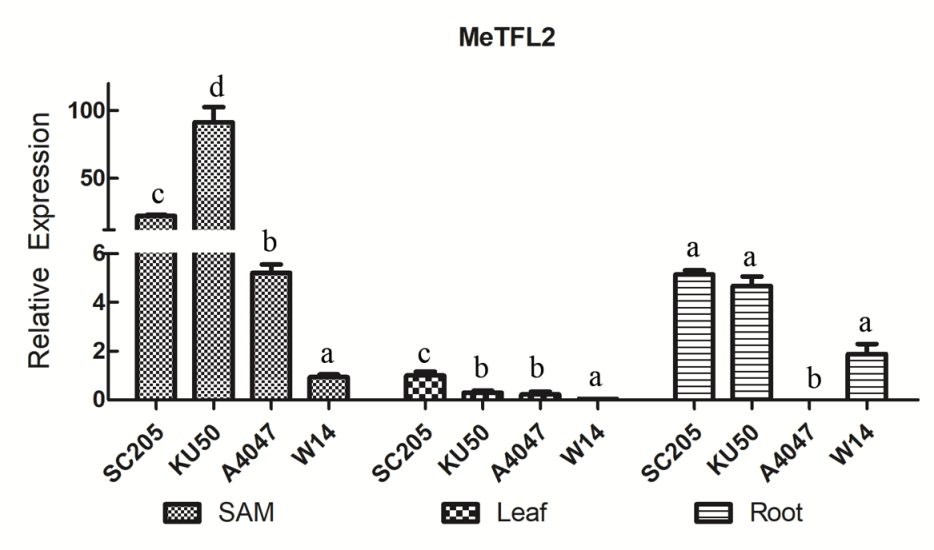

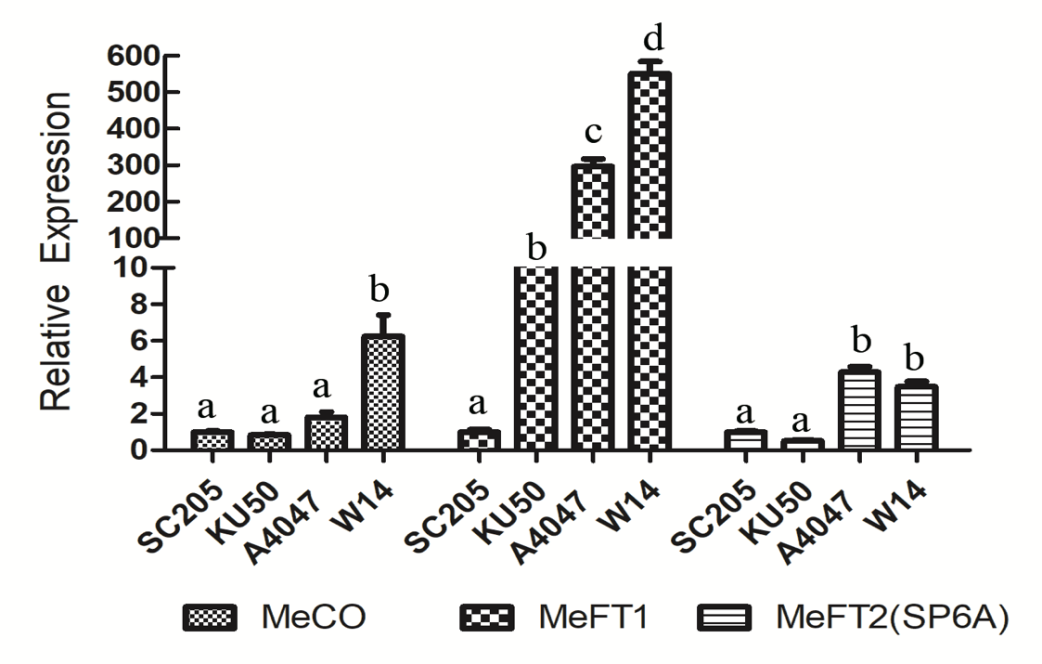

#### Fig. S28. Differential expressions of genes in cyanide metabolism between wild ancestors to cultivated cassava validated by Real-Time Q-PCR

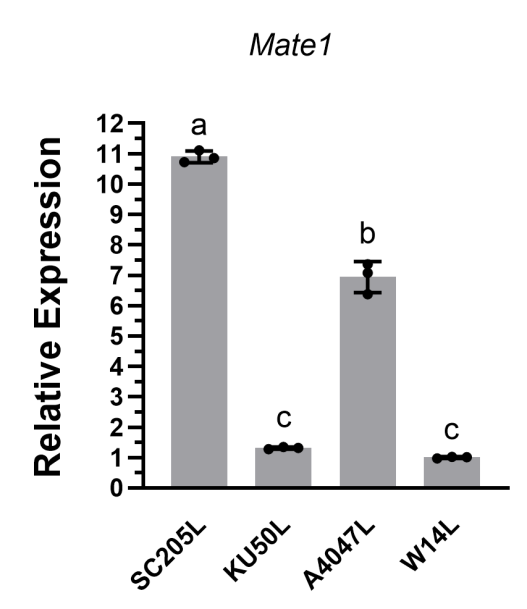

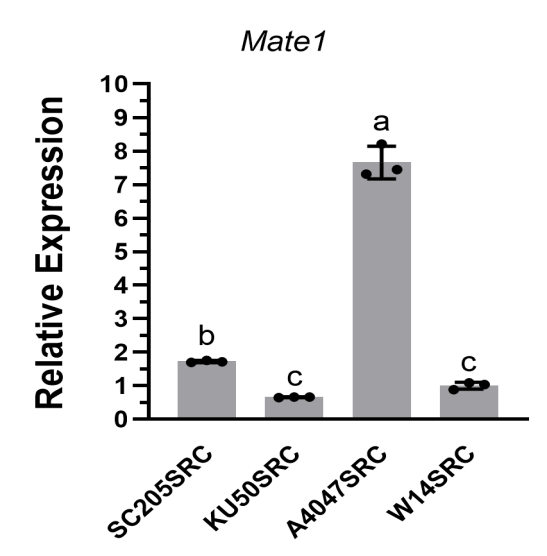

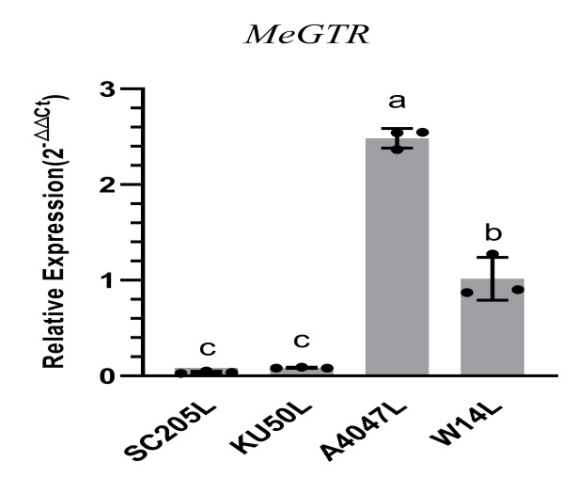

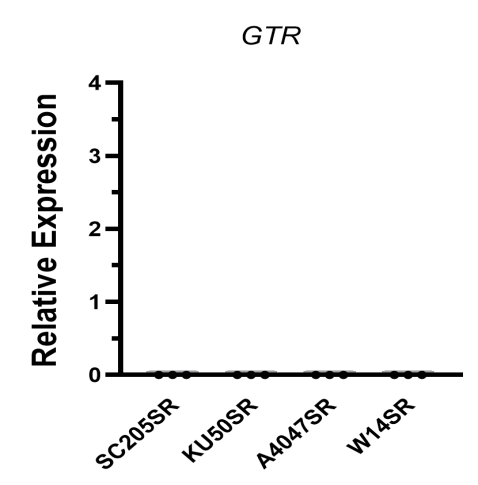

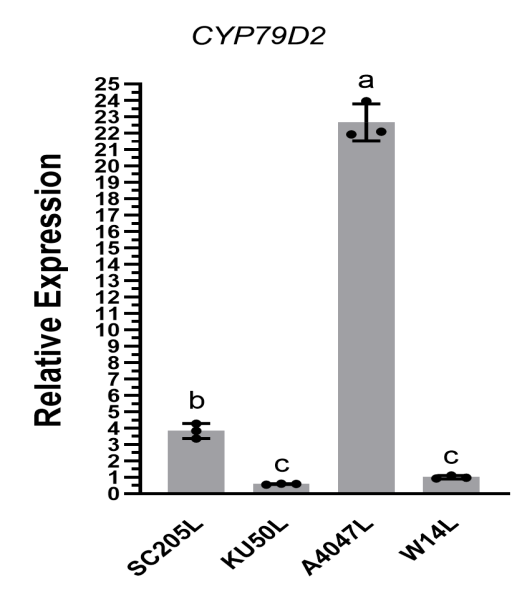

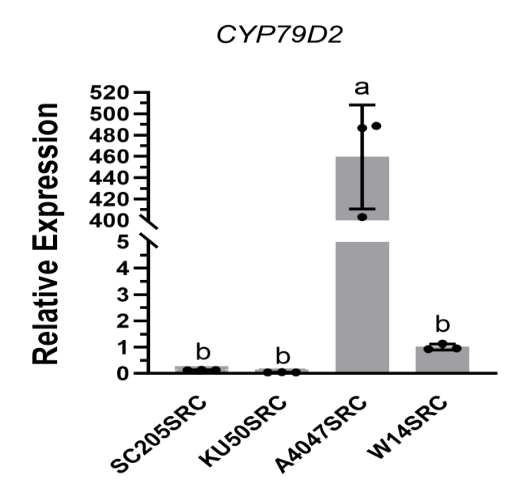

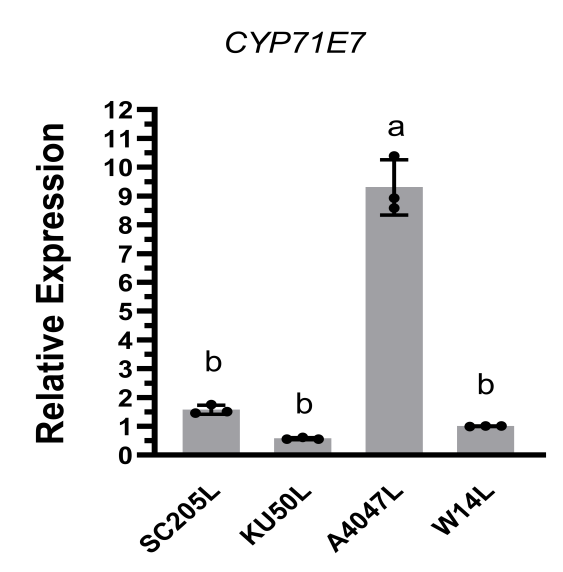

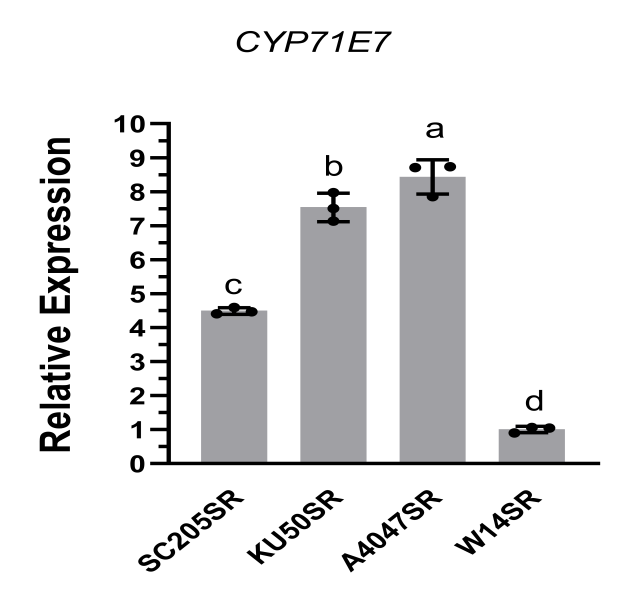

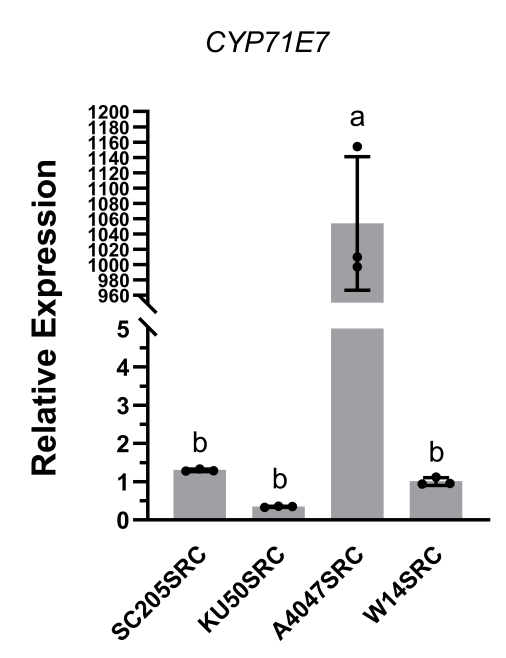

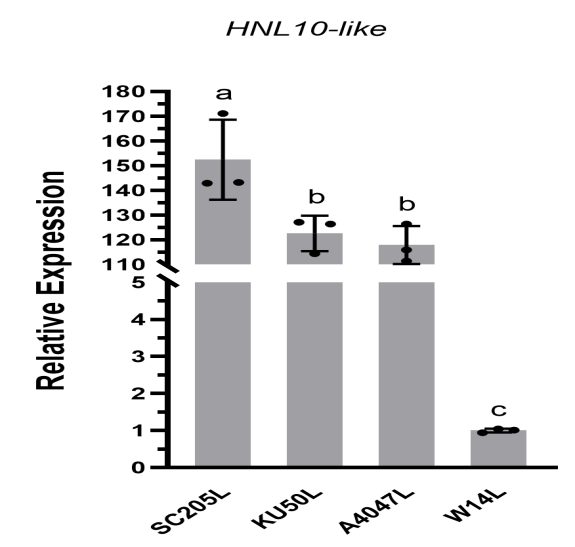

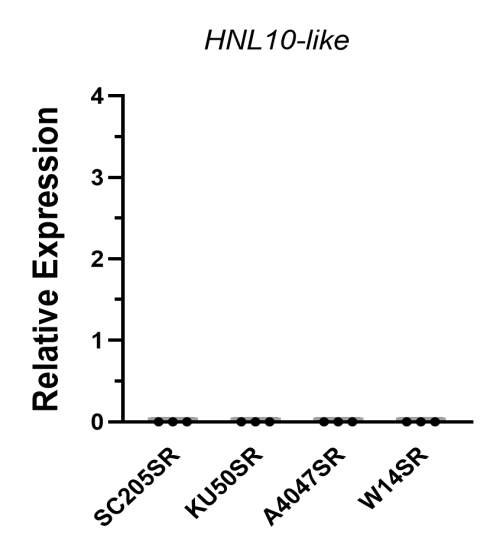

Notes: L, leaves; R, roots; SR, storage roots; SRC, Column of storage roots.

### Supplementary Table

#### Table S1. The genome sizes estimated by the flow cytometry

| **Species & Genotype** | **Reference genome** | **Genome size(Gb)** | **Average(Gb)** | **Estimated genome size** |
| --- | --- | --- | --- | --- |
| *M.esculenta* Crantz, KU50 | Maize & rice | 0.72; 0.72; 0.70 0.74 0.70 | 0.72 | 0.72±0.02 |
| *M.esculenta* SSP. *flabellifolia*, FLA4047 | Maize & rice | 0.72; 0.70; 0.70; 0.74; 0.72 | 0.72 | 0.72±0.02 |
| *M.esculenta* SSP. *flabellifolia*, W14 | Maize & rice | 0.84; 0.81; 0.81; 0.86; 0.83 | 0.83 | 0.83±0.016 |
| *M.glaziovii*, GLAyn | Maize & rice | 0.73; 0.71; 0.71; 0.77; 0.74 | 0.73 | 0.73±0.016 |
| *M.glaziovii* , GLAb | Maize & rice | 0.74; 0.72; 0.72; 0.75; 0.73 | 0.73 | 0.73±0.01 |

#### Table S2. Assembly and genome characteristics of the three cassava genomes before adding 2 cells of the ONT ultra-long reads each one

|  | **W14** | **AM560** | **FLA4047** | **AM560JGIv7.1** |
| --- | --- | --- | --- | --- |
| Sequence number | 18chr | 18chr/2scf/1387ctg | 18chr/137ctg | 18chr/881scaf |
| N50(Mb) | 39.04 | 36.23 | 36.53 | 33.08 |
| GC content | 0.37 | 0.37 | 0.36 | 0.37 |
| Assembly length (Mb) | 681.30 | 730.56 | 674.41 | 669.37 |
| Repeats(bp) | 421,935,433 | 471,151,664 | 454,937,826 | 402,958,641 |
| % | 61.93% | 64.49% | 67.46% | 60.20% |
| Coding gene number | 30,320 | 31,585 | 34,153 | 33,849 |
| Average transcript length(bp) | 4276.13 | 4286.81 | 3937.05 | 4190.73 |
| lncRNA number | 8,687 | 12,253 | 12,683 | NA |

#### Table S3. Gene Annotation of the three cassava genomes

**A.**

|  | W14 | AM560 | FLA4047 | JGIv7.1 |
| --- | --- | --- | --- | --- |
| Gene number | 30320 | 31585 | 34153 | 33849 |
| Average transcript length | 4276.13 | 4286.81 | 3937.05 | 4190.73 |
| Average CDS length | 1182.8 | 1169.63 | 1111.08 | 1255.97 |
| Average exons per gene | 5.37 | 5.13 | 5.11 | 5.13 |
| Average exon length | 220.14 | 227.98 | 217.46 | 244.68 |
| Average intron length | 583.61 | 524.12 | 489.2 | 513.93 |

**B.**

|  | FLA4047 | AM560 | W14 |
| --- | --- | --- | --- |
| Number of genes | 30336 | 38601 | 37046 |
| Number of genes in orthogroups | 28992 | 37254 | 35305 |
| Number of unassigned genes | 1344 | 1347 | 1741 |
| Percentage of genes in orthogroups | 95.6 | 96.5 | 95.3 |
| Percentage of unassigned genes | 4.4 | 3.5 | 4.7 |
| Number of orthogroups containing species | 19676 | 19194 | 20111 |
| Percentage of orthogroups containing species | 89.2 | 87 | 91.1 |
| Number of species-specific orthogroups | 173 | 573 | 408 |
| Number of genes in species-specific orthogroups | 917 | 5226 | 4770 |
| Percentage of genes in species-specific orthogroups | 3 | 13.5 | 12.9 |

**C.**

**Orthologues Stats Totals**

|  | FLA4047 | AM560 | W14 |
| --- | --- | --- | --- |
| FLA4047 | 0 | 24287 | 26155 |
| AM560 | 24720 | 0 | 28130 |
| W14 | 25002 | 26428 | 0 |

**D.**

**Orthogroups Species Overlaps**

|  | FLA4047 | AM560 | W14 |
| --- | --- | --- | --- |
| FLA4047 | 19676 | 17214 | 18296 |
| AM560 | 17214 | 19194 | 17414 |
| W14 | 18296 | 17414 | 20111 |

#### Table S4. Non-coding RNA annotated in AM560, FLA4047 and W14 genomes

|  | **W14** | **AM560** | **FLA4047** |
| --- | --- | --- | --- |
| miRNA | 175 | 164 | 162 |
| lncRNA | 3,212 | 4,019 | 4,126 |
| rRNA | 132 | 438 | 239 |
| tRNA | 1,067 | 1,064 | 1,011 |
| total | 4,586 | 5,685 | 5,538 |

#### Table S5. Assembly and annotation of 21 genomes used for pan-genome analysis

| Accession  Name | Contig N50 (Kb) | Scaffold N50  (Mb) | Assembly  Size (Mb) | Annotated gene number |
| --- | --- | --- | --- | --- |
| BC006 | 843.15 | 34.60 | 843.15 | 34088 |
| BC008 | 550.11 | 25.21 | 638.06 | 37798 |
| Baxi-1 | 155.40 | 36.04 | 675.17 | 33826 |
| Fuxuan01 | 246.30 | 24.67 | 638.07 | 35536 |
| GR4 | 5229.75 | 30.85 | 664.27 | 32155 |
| HB60 | 330.02 | 29.51 | 739.51 | 37247 |
| IB003 | 152.53 | 27.45 | 777.33 | 41950 |
| IB005 | 265.69 | 30.90 | 763.88 | 37541 |
| ITBB04 | 286.39 | 28.77 | 749.74 | 38987 |
| Mianbao | 209.30 | 27.23 | 719.32 | 37085 |
| NZ199 | 257.66 | 27.86 | 718.74 | 34336 |
| SC205 | 134.11 | 25.06 | 692.31 | 41013 |
| SC5 | 1380.10 | 31.14 | 768.39 | 33213 |
| SC6068 | 1818.90 | 33.68 | 827.94 | 33971 |
| SC8 | 188.79 | 26.22 | 699.83 | 36704 |
| SC9 | 403.18 | 27.59 | 742.47 | 39612 |
| Thai 01 | 246.30 | 36.28 | 849.08 | 34846 |
| Wenchang red | 133.20 | 27.07 | 769.97 | 30245 |
| ZM9781 | 180.76 | 27.85 | 718.25 | 36795 |
| GLAyn | 239.30 | 26.09 | 829.22 | 35069 |
| GLAb | 1187.36 | 27.65 | 614.95 | 30104 |

#### Table S6. Number and structure of genes annotated in 24 genomes integrated into pan genome of cassava

|  | Number of gene | cds region(M) | Average cds length(bp) | Average exons per gene | Bases masked |
| --- | --- | --- | --- | --- | --- |
| SC8 | 36704 | 39.6 | 1081.2 | 5.2 | 62.11% |
| SC5 | 33213 | 39.1 | 1161.1 | 5.3 | 67.20% |
| Baxi | 33826 | 38.1 | 1127.5 | 5.4 | 61.94% |
| Wenchang red | 44945 | 47.5 | 1057.8 | 4.8 | 61.80% |
| Fuxuan | 35536 | 40.1 | 1128.3 | 5.3 | 60.36% |
| SC6068 | 33971 | 39.4 | 1161.1 | 5.3 | 68.80% |
| IBO3 | 41950 | 44.9 | 1071.9 | 52 | 61.16% |
| BC008 | 34088 | 39.3 | 1147.2 | 5.3 | 61.51% |
| ZM9781 | 36795 | 39.7 | 1078.2 | 52 | 62.54% |
| HB60 | 37247 | 41.6 | 1117.6 | 5.3 | 63.13% |
| SC9 | 39612 | 43.8 | 1107.1 | 5.2 | 61.51% |
| IBO5 | 37541 | 40.6 | 1084.1 | 52 | 63.54% |
| Mianbao | 37085 | 40.3 | 1085.4. | 52 | 62.58% |
| BC006 | 37798 | 423 | 1119.8 | 5.2 | 66.17% |
| GR4 | 32155 | 37.9 | 1178.7 | 5.4 | 64.28% |
| ITBBO4 | 38987 | 45.4 | 1166.6 | 5.3 | 62.22% |
| NZ199 | 34336 | 39.7 | 1157.11 | 5.4 | 64.60% |
| SC205 | 41013 | 432 | 1054.8 | 5.1 | 58.10% |
| Taiyin | 34846 | 39.5 | 1135 | 5.3 | 69.02% |
| GLAyn | 35069 | 42.9 | 1224.1 | 5.5 | 67.60% |
| AM560 | 35872 | 44.16 | 1231.1 | 5.6 | 65.70% |
| FLA4047 | 30336 | 36 | 1188 | 5.3 | 63.73% |
| W14 | 37046 | 42.9 | 1160.6 | 5.1 | 67.86% |

#### Table S7. The common and private gene numbers annotated among 24 genomes

| Orthogroup | core | Softcore | dispensable | private | Total |
| --- | --- | --- | --- | --- | --- |
| W14 | 11679 | 6439 | 15544 | 2851 | 36513 |
| GLAyn | 10981 | 6236 | 10268 | 24 | 27509 |
| GLAb | 10790 | 6160 | 10402 | 14 | 27366 |
| FLA4047 | 12140 | 6476 | 11186 | 62 | 29864 |
| AM560 | 10870 | 5763 | 17127 | 1681 | 35441 |
| BC006 | 11457 | 6430 | 12290 | 49 | 30226 |
| BC008 | 10844 | 6144 | 10402 | 28 | 27418 |
| Fuxuan | 11084 | 6409 | 10920 | 23 | 28436 |
| GR4 | 10931 | 6171 | 10443 | 23 | 27568 |
| HB60 | 11231 | 6497 | 11234 | 24 | 28968 |
| IB003 | 11750 | 6784 | 11449 | 35 | 30018 |
| IB005 | 11413 | 6567 | 11660 | 41 | 29690 |
| SC16 | 11406 | 6607 | 11914 | 16 | 29943 |
| Mianbao | 11142 | 6389 | 11150 | 57 | 28738 |
| NZ199 | 11090 | 6349 | 11055 | 22 | 28516 |
| SC205 | 11527 | 6658 | 11536 | 48 | 29769 |
| SC5 | 10898 | 6254 | 11183 | 29 | 28364 |
| SC6068 | 11139 | 6388 | 11194 | 15 | 28736 |
| SC8 | 11268 | 6403 | 10802 | 57 | 28530 |
| SC9 | 11521 | 6685 | 11886 | 44 | 30136 |
| Taiyin | 10996 | 6318 | 11804 | 29 | 29147 |
| Wenchang red | 11401 | 6683 | 11580 | 57 | 29721 |
| ZM9781 | 11413 | 6514 | 11008 | 56 | 28991 |
| Baxi | 12516 | 7097 | 12536 | 69 | 32218 |

#### Table S8. SV types and number statistic in the pan genome

| Sample ID | DEL | DUP | INS | INV | TRA |
| --- | --- | --- | --- | --- | --- |
| GLAyn | 53,694 | 3,093 | 23,057 | 1,149 | 8,370 |
| GLAb | 54,776 | 3,508 | 21,018 | 1,208 | 10,365 |
| ZM9781 | 45,100 | 2,446 | 15,494 | 867 | 6,361 |
| IB003 | 52,105 | 2,770 | 15,662 | 860 | 7,658 |
| SC8 | 45,072 | 2,512 | 15,741 | 910 | 6,680 |
| Baxi-1 | 47,531 | 2,695 | 17,304 | 996 | 7,359 |
| Mianbao | 48,871 | 2,850 | 16,832 | 994 | 7,641 |
| SC205 | 52,507 | 3,420 | 20,295 | 1,224 | 9,223 |
| IB005 | 47,632 | 2,720 | 16,111 | 919 | 7,293 |
| HB60 | 51,132 | 2,937 | 17,620 | 1,003 | 7,660 |
| SC16 | 51,111 | 3,170 | 18,200 | 1,092 | 8,650 |
| Wenchang red | 58,245 | 3,898 | 23,629 | 1,327 | 11,223 |
| Fuxuan01 | 58,190 | 3,848 | 23,443 | 1,352 | 11,458 |
| BC008 | 58,456 | 3,774 | 22,019 | 1,404 | 11,132 |
| SC9 | 58,742 | 3,884 | 21,879 | 1,410 | 11,515 |
| NZ199 | 58,467 | 3,748 | 22,705 | 1,373 | 11,571 |
| SC6068 | 65,100 | 4,528 | 23,039 | 1,622 | 14,275 |
| SC5 | 67,439 | 4,825 | 24,312 | 1,810 | 15,525 |
| BC006 | 71,066 | 5,101 | 24,745 | 1,878 | 16,627 |
| Taiyin01 | 72,245 | 5,278 | 25,543 | 1,976 | 17,754 |
| GR4 | 71,999 | 5,281 | 26,735 | 1,967 | 18,496 |
| Total | 176,801 | 19,041 | 80,750 | 6,257 | 63,473 |
| % | 51.05 | 5.5 | 23.32 | 1.81 | 18.33 |

DEL：Deletion； INS：Insertion； DUP：Duplication；INV：Inversion；TRA：Translocation

#### Table S9. SNP and InDel diversity statistic between ME (298 accessions) and FLA (28 accessions)

| **SNV** | **Location** | **ME (198 accessions)** | **FLA (28 accessions)** |
| --- | --- | --- | --- |
| SNP |  |  |  |
|  | Total: | 13735173 | 22032832 |
|  | **gene region** | **2891925** | **3679623** |
|  | UTR | 189432 | 251873 |
|  | CDS | 614140 | 944295 |
|  | Intron | 2088353 | 2483455 |
|  | **Non-gene region** | **10843248** | **18353209** |
| InDel |  |  |  |
|  | Total: | 2719151 | 4533604 |
|  | **gene region** | **434384** | **681160** |
|  | UTR | 40407 | 67259 |
|  | CDS | 25213 | 45949 |
|  | Intron | 368764 | 567952 |
|  | **Non-gene region** | **2284767** | **3852444** |

#### Table S10. Summary of the RNA-Seq data used for photosynthesis, flowering and storage root development and cyanide metabolism

| Code | Sample Name | Clean Reads | Clean Base | Read Length | Q20 (%) | GC(%) |
| --- | --- | --- | --- | --- | --- | --- |
| 1 | FLA433-2_L_BR1 | 24061821 | 7218546300 | PE150 | 96.55 | 45.62 |
| 2 | FLA433-2_L_BR2 | 24127123 | 7238136900 | PE150 | 96.41 | 44.86 |
| 3 | FLA433-2_R_BR2 | 23716101 | 7114830300 | PE150 | 97.29 | 44.43 |
| 4 | FLA449-1_L_BR1 | 24047043 | 7214112900 | PE150 | 96.5 | 44.5 |
| 5 | FLA449-1_L_BR2 | 24144383 | 7243314900 | PE150 | 96.67 | 44.49 |
| 6 | FLA449-1_R_BR1 | 24076305 | 7222891500 | PE150 | 98.03 | 44.43 |
| 7 | FLA449-1_R_BR2 | 24134900 | 7240470000 | PE150 | 97.08 | 44.28 |
| 8 | FLA467-4_L_BR2 | 24190338 | 7257101400 | PE150 | 97.06 | 44.63 |
| 9 | FLA467-4_R_BR1 | 24150158 | 7245047400 | PE150 | 97.34 | 44.51 |
| 10 | FLA467-4_R_BR2 | 24235310 | 7270593000 | PE150 | 97.28 | 44.7 |
| 11 | FLA467-7_L_BR1 | 24044889 | 7213466700 | PE150 | 96.55 | 45.47 |
| 12 | FLA467-7_L_BR2 | 24000499 | 7200149700 | PE150 | 96.52 | 44.44 |
| 13 | FLA467-7_R_BR1 | 24135660 | 7240698000 | PE150 | 97.07 | 44.43 |
| 14 | FLA4047 L | 24105217 | 7231565100 | PE150 | 97.66 | 44.21 |
| 15 | FLA4047 L night | 24064592 | 7219377600 | PE150 | 97.69 | 44.12 |
| 16 | FLA4047 XSR | 24105368 | 7231610400 | PE150 | 97.68 | 43.7 |
| 17 | FLA4047 PSR | 24100581 | 7230174300 | PE150 | 97.72 | 44.59 |
| 18 | A4047 FP | 24033408 | 7210022400 | PE150 | 97.58 | 44.14 |
| 19 | KU50 L | 24014991 | 7204497300 | PE150 | 97.54 | 46.58 |
| 20 | KU50 L night | 24073819 | 7222145700 | PE150 | 97.53 | 44.73 |
| 21 | KU50 XSR | 24096842 | 7229052600 | PE150 | 97.58 | 47.77 |
| 22 | KU50 PSR | 24125789 | 7237736700 | PE150 | 97.65 | 45.84 |
| 23 | KU50 SAM | 24136279 | 7240883700 | PE150 | 97.61 | 46.15 |
| 24 | SC205 L | 24051624 | 7215487200 | PE150 | 97.63 | 44.42 |
| 25 | SC205 L night | 24004688 | 7201406400 | PE150 | 97.64 | 44.4 |
| 26 | SC205 XSR | 24058498 | 7217549400 | PE150 | 97.84 | 44.38 |
| 27 | SC205 PSR | 24110022 | 7233006600 | PE150 | 97.62 | 44.81 |
| 28 | SC205 SAM | 24029085 | 7208725500 | PE150 | 97.52 | 44.87 |
| 29 | W14 L | 24011323 | 7203396900 | PE150 | 97.6 | 44.61 |
| 30 | W14 L night | 24007216 | 7202164800 | PE150 | 97.7 | 44.01 |
| 31 | W14 XSR | 24031313 | 7209393900 | PE150 | 97.69 | 44.15 |
| 32 | W14 PSR | 24102720 | 7230816000 | PE150 | 97.56 | 44.23 |
| 33 | W14 FP | 24070632 | 7221189600 | PE150 | 97.74 | 43.92 |
| 34 | W14 L，DAP120 | 25483611 | 7645083300 | PE150 | 97.63 | 43.85 |
| 35 | GLAyn L，DAP120 | 23184105 | 6955231500 | PE150 | 97.81 | 43.96 |
| 36 | GLAb L，DAP120 | 22712028 | 6813608400 | PE150 | 97.93 | 45.53 |
| 37 | NZ199 L，DAP120 | 26702381 | 8010714300 | PE150 | 98.1 | 47.13 |
| 38 | SC8 L，DAP120 | 27265453 | 8179635900 | PE150 | 97.53 | 45.95 |
| 39 | SC5 L，DAP120 | 27371816 | 8211544800 | PE150 | 97.19 | 45.44 |
| 40 | BAXI L，DAP120 | 22820474 | 6846142200 | PE150 | 97.26 | 45.53 |
| 41 | FUXUAN L，DAP120 | 23243140 | 6972942000 | PE150 | 97.37 | 43.47 |
| 42 | IB05 L，DAP120 | 22751029 | 6825308700 | PE150 | 97.65 | 45.52 |
| 43 | MIANBAO L，DAP120 | 22963493 | 6889047900 | PE150 | 98.03 | 43.86 |
| 44 | WENCHAN Red L，DAP120 | 23114331 | 6934299300 | PE150 | 97.52 | 44.02 |
| 45 | SC9 L，DAP120 | 23136238 | 6940871400 | PE150 | 96.98 | 43.05 |
| 46 | HB60 L，DAP120 | 24810581 | 7443174300 | PE150 | 97.1 | 45.65 |
| 47 | SC205 L，DAP120 | 23096981 | 6929094300 | PE150 | 97.48 | 43.4 |
| 48 | GR4 L，DAP120 | 27744112 | 8323233600 | PE150 | 96.79 | 46.8 |
| 49 | SC6068 L，DAP120 | 23708337 | 7112501100 | PE150 | 97.82 | 47.85 |
| 50 | TAIYIN L，DAP120 | 23099357 | 6929807100 | PE150 | 97.59 | 44.93 |
| 51 | BC006 L，DAP120 | 27451902 | 8235570600 | PE150 | 97.44 | 44.7 |
| 52 | BC008 L，DAP120 | 22927812 | 6878343600 | PE150 | 97.63 | 45.73 |
| 53 | IB03 L，DAP120 | 23588696 | 7076608800 | PE150 | 98.42 | 45.72 |
| 54 | ZM9781 L，DAP120 | 22708381 | 6812514300 | PE150 | 97.61 | 46.39 |
| 55 | SC205_mesophyll, C1, DAP45 | 25245073 | 7573521900 | PE150 | 97.31 | 47.4 |
| 56 | SC205_leaf_vein,C2,DAP45 | 24565569 | 7369670700 | PE150 | 98.42 | 47.52 |
| 57 | SC205_phloem_fibrous_root, C4, DAP45 | 23920975 | 7176292500 | PE150 | 97.33 | 45.35 |
| 58 | SC205_xylem_fibrous_root,C5, DAP45 | 29251465 | 8775439500 | PE150 | 98.21 | 46.65 |
| 59 | KU50_mesophyll, C6, DAP45 | 21267283 | 6380184900 | PE150 | 98.11 | 44.47 |
| 60 | KU50_leaf_vein, C7, DAP45 | 25735790 | 7720737000 | PE150 | 98.36 | 43.36 |
| 61 | KU50_phloem_fibrous_root, C9, DAP45 | 25939034 | 7781710200 | PE150 | 98.76 | 43.46 |
| 62 | KU50_xylem_fibrous_root, C10, DAP45 | 24758704 | 7427611200 | PE150 | 97.64 | 43.67 |
| 63 | SC205_mesophyll, C11, DAP53 | 23708885 | 7112665500 | PE150 | 97.98 | 46.76 |
| 64 | SC205_leaf_vein, C12, DAP53 | 26557634 | 7967290200 | PE150 | 97.34 | 46.03 |
| 65 | SC205_phloem_storage_root, C16, DAP53 | 26159495 | 7847848500 | PE150 | 97.65 | 46.74 |
| 66 | SC205_xylem_storage_root, C17, DAP53 | 26472643 | 7941792900 | PE150 | 97.86 | 45.06 |
| 67 | KU50_mesophyll, C18, DAP53 | 29684837 | 8905451100 | PE150 | 97.34 | 44.93 |
| 68 | KU50_leaf_vein_20190524, C19, DAP53 | 20547954 | 6164386200 | PE150 | 97.56 | 44.05 |
| 69 | KU50_phloem_storage_root, C23, DAP53 | 22776391 | 6832917300 | PE150 | 98.61 | 43.53 |
| 70 | KU50_xylem_storage_root, C24, DAP53 | 24152707 | 7245812100 | PE150 | 97.57 | 44.64 |
| 71 | SC205_mesophyll,20190528, C25, DAP58 | 23724370 | 7117311000 | PE150 | 97.54 | 47.95 |
| 72 | SC205_leaf_vein, C26, DAP58 | 28772709 | 8631812700 | PE150 | 98.22 | 47.29 |
| 73 | SC205_phloem_storage_root, C30, DAP58 | 23196247 | 6958874100 | PE150 | 97.86 | 45.69 |
| 74 | SC205_xylem_storage_root, C31, DAP58 | 32269496 | 9680848800 | PE150 | 97.85 | 44.54 |
| 75 | KU50_mesophyll, C32, DAP58 | 36105465 | 10831639500 | PE150 | 97.43 | 44.2 |
| 76 | KU50_leaf_vein, C33, DAP58 | 34373038 | 10311911400 | PE150 | 97.32 | 43.74 |
| 77 | KU50_phloem_storage_root C37, DAP58 | 22032896 | 6609868800 | PE150 | 96.89 | 42.18 |
| 78 | KU50_xylem_storage_root, C38,DAP58 | 21811044 | 6543313200 | PE150 | 97.41 | 43.29 |
| 79 | SC205_mesophyll_20190531, C39, DAP61 | 25381690 | 7614507000 | PE150 | 98.22 | 47.17 |
| 80 | SC205_leaf_vein,C40,DAP61 | 33830575 | 10149172500 | PE150 | 96.21 | 46.49 |
| 81 | SC205_phloem_storage_root, C44, DAP61 | 23794395 | 7138318500 | PE150 | 97.46 | 44.93 |
| 82 | SC205_xylem_storage_root, C45, DAP61 | 24358721 | 7307616300 | PE150 | 97.58 | 46.54 |
| 83 | KU50_mesophyll, C46, DAP61 | 29718690 | 8915607000 | PE150 | 98.13 | 43.83 |
| 84 | KU50_leaf_vein, C47, DAP61 | 24182318 | 7254695400 | PE150 | 97.84 | 42.93 |
| 85 | KU50_phloem_storage_root, C51, DAP61 | 24115235 | 7234570500 | PE150 | 97.21 | 43.05 |
| 86 | KU50_xylem_storage_root, C52, DAP61 | 23321890 | 6996567000 | PE150 | 98.16 | 43.87 |
| 87 | SC205_mesophyll, 20190606, C53, DAP67 | 27230315 | 8169094500 | PE150 | 97.68 | 47.06 |
| 88 | SC205_leaf_vein, C54, DAP67 | 23173106 | 6951931800 | PE150 | 96.42 | 46.72 |
| 89 | SC205_phloem_storage_root, C58, DAP67 | 21844553 | 6553365900 | PE150 | 97.76 | 45.94 |
| 90 | SC205_xylem_storage_root, C59, DAP67 | 21953523 | 6586056900 | PE150 | 97.51 | 47.83 |
| 91 | KU50_mesophyll, C60, DAP67 | 26733663 | 8020098900 | PE150 | 96.11 | 44.01 |
| 92 | KU50_leaf_vein, C61, DAP67 | 24454788 | 7336436400 | PE150 | 98.14 | 43.29 |
| 93 | KU50_phloem_storage_root, C65, DAP67 | 23162503 | 6948750900 | PE150 | 97.45 | 43.38 |
| 94 | KU50_xylem_storage_root, C66, DAP67 | 27322941 | 8196882300 | PE150 | 97.92 | 43.07 |
| 95 | SC205_mesophyll_20190618,C67, DAP79 | 26228535 | 7868560500 | PE150 | 97.63 | 47.58 |
| 96 | SC205_leaf_vein, C68, DAP79 | 26771905 | 8031571500 | PE150 | 97.84 | 47.04 |
| 97 | SC205_phloem_storage_root, C72, DAP79 | 27406883 | 8222064900 | PE150 | 97.64 | 44.92 |
| 98 | SC205_xylem_storage_root, C73,DAP79 | 25627589 | 7688276700 | PE150 | 98.52 | 46.07 |
| 99 | KU50_mesophyll, C74, DAP79 | 23565693 | 7069707900 | PE150 | 97.65 | 44.9 |
| 100 | KU50_leaf_vein, C75, DAP79 | 22064817 | 6619445100 | PE150 | 97.03 | 42.57 |
| 101 | KU50_phloem_storage_root_20190618,C79, DAP79 | 22899131 | 6869739300 | PE150 | 98.67 | 43.29 |
| 102 | KU50_xylem_storage_root_20190618, C80, DAP79 | 18955917 | 5686775100 | PE150 | 97.09 | 43.68 |
| 103 | W14_mesophyll_20190618,C81,DAP79 | 24563084 | 7368925200 | PE150 | 96.5 | 43.29 |
| 104 | W14_leaf_vein_20190618,C82, DAP79 | 28090595 | 8427178500 | PE150 | 97.61 | 43.99 |
| 105 | W14_phloem_fibrous_root_20190618,C84, DAP79 | 22643322 | 6792996600 | PE150 | 97.08 | 43.76 |
| 106 | W14_xylem_fibrous_root_20190618, C85, DAP79 | 24337571 | 7301271300 | PE150 | 98.02 | 43.68 |
| 107 | SC205_mesophyll_20190706, C93, DAP97 | 25760329 | 7728098700 | PE150 | 97.13 | 47.83 |
| 108 | SC205_leaf_vein_20190706, C94, DAP97 | 27288655 | 8186596500 | PE150 | 97.17 | 47.55 |
| 109 | SC205_phloem_storage_root_20190706, C98, DAP97 | 26363955 | 7909186500 | PE150 | 98.06 | 42.49 |
| 110 | SC205_xylem_storage_root_20190706,C99, DAP97 | 24264019 | 7279205700 | PE150 | 97.87 | 43.28 |
| 111 | KU50_mesophyll_20190706, C100, DAP97 | 22799421 | 6839826300 | PE150 | 97.53 | 44.72 |
| 112 | KU50_leaf_vein_20190706,C101, DAP97 | 26718348 | 8015504400 | PE150 | 96.33 | 42.86 |
| 113 | KU50_phloem_storage_root_20190706, C105, DAP97 | 23699502 | 7109850600 | PE150 | 97.98 | 42.64 |
| 114 | KU50_xylem_storage_root_20190706, C106, DAP97 | 29230948 | 8769284400 | PE150 | 97.83 | 42.9 |
| 115 | W14_mesophyll_20190706, C107, DAP97 | 25539819 | 7661945700 | PE150 | 97.81 | 46.47 |
| 116 | W14_leaf_vein_20190706, C108, DAP97 | 24632662 | 7389798600 | PE150 | 98.03 | 45.36 |
| 117 | W14_phloem_storage_root_20190706, C112, DAP97 | 23267639 | 6980291700 | PE150 | 96.87 | 43.25 |
| 118 | W14_xylem_storage_root_20190706, C113, DAP97 | 20286563 | 6085968900 | PE150 | 87.42 | 43.66 |

#### Table S11 Cyanoside content in leaves and storage roots of wild and cultivated cassava clones (2020-2021)

| Cassava clones | Storage root（μg/g） | E | Leave（μg/g） | E | SRw/SRc | Lw/Lc |
| --- | --- | --- | --- | --- | --- | --- |
| W14 | 196.75 | 4.40 | 1326.25 | 19.87 | 2.02 | 1.00 |
| FLA A4047 | 397.89 | 1.17 | 754.37 | 5.34 | 1.00 | 1.76 |
| KU50 | 281.79 | 4.31 | 811.18 | 13.23 | 1.41 | 1.63 |
| NZ199 | 85.35 | 1.38 | 872.81 | 2.38 | 4.66 | 1.52 |
| SC205 | 99.78 | 1.49 | 513.77 | 9.72 | 3.99 | 2.58 |
| SC8 | 86.01 | 2.46 | 584.51 | 12.42 | 4.63 | 2.27 |
| SC16 | 79.86 | 0.87 | 379.37 | 4.92 | 4.98 | 3.50 |
| SC9 | 70.89 | 0.87 | 650.72 | 15.00 | 5.61 | 2.04 |
| Average |  |  |  |  |  |  |

#### Table S12 Accessions with high and low cyanide glycoside in storage root

| **Accessions with low cyanide** | | **Accessions with high cyanide** | |
| --- | --- | --- | --- |
| Name | Cyanide glycoside in storage root（μg/g) | Name | Cyanide glycoside in storage root (μg/g) |
| 18-9 | 20.23 | 44-7 | 188.5 |
| 39-2 | 51.87 | 32-5 | 237.9 |
| 20-6 | 57.72 | 57-6 | 208.65 |
| 58-7 | 45.63 | F297 | 226.72 |
| 46-14 | 76.44 | 47-11 | 316.16 |
| F039 | 47.83 | 03-11 | 292.89 |
| SC16 | 29.74 | 16-14 | 218.27 |
| F173 | 43.16 | 50-2 | 233.09 |
| ITBB01 | 67.08 | 05-3 | 234.26 |
| F128 | 51.87 | 14-19 | 157.17 |
| SC5 | 57.90 | 49-20 | 258.18 |
| 46-12 | 49.82 | 01-14 | 298.35 |
| SC205 | 99.78 | KU50 | 281.79 |
| SC9 | 70.89 | A4047 | 397.89 |
| SC8 | 86.01 | W14 | 196.75 |

#### Table S13 the genes involved in biosynthesis, transportation and decomposition of cyanide

| Tracking_id | Gene name | Pathway |
| --- | --- | --- |
| Manes.07G027100 | Linamarase | Cyanide decomposition |
| **Manes.13G092100** | ***MeHNL10-like*** |  |
| Manes.13G092600 | *MeHNL4-like* |  |
| Manes.13G094600 | *MeHNL24-like* |  |
| Manes.05G033000 | *B-CASa* |  |
| Manes.01G255400 | *B-CASb* |  |
| Manes.11G164700 | *NIT4.1* |  |
| Manes.04G000700 | *NIT4.2* |  |
| Manes.12G133500 | *CYP79D2* | Cyanide biosynthesis |
| Manes.13G094200 | *CYP79D1* |  |
| Manes.12G132900 | CYP71E7 |  |
| Manes.12G132800 | CYP71E11 |  |
| **Manes.12G132300** | **UGT85K4** |  |
| Manes.12G133000 | *UGT85K5* |  |
| **Manes.15G180400** | **CGTR1** | **Cyanide transportation** |
| Manes.17G021100 | CGTR2 |  |
| Manes.17G124600 | CGTR |  |
| **Manes.15G176100** | ***GTR*** |  |
| **Manes.16G007900** | **MATE1** |  |
| Manes.16G008000 | MATE2 |  |
| Manes.14G074300 | HPP，nitrite transport activity |  |
| Manes.14G073900 | H^+^-ATPase |  |

#### Table S14 list of primers for Q-PCR validation of genes for photosynthesis, flowering and tuberization and cyanide glycoside metabolism

| **Gene name & ID** | **Right primer sequences** | **Left primer sequences** | **Information** |
| --- | --- | --- | --- |
| *MePC*, 05G142000 | ACGGCTTCTTCAGTCCCAAA | CCCCGGAAGCTACAGAGAAA | Photosynthesis related |
| *MeFNR2*，02G105100 | AAGACGGCATTGATTGGTTGG | TCATTACAGGATGCCTCTGC |  |
| *MeFNR3*，18G012800 | GGCTCAAGGGAATGGAGAAGG | TGATGGCTAGTGCCTGCTAAAA |  |
| *psbA,*（D1protein） 5999949 | CTGATGGTATGCCTCTAGGA | CTTCCTCTTGACCGAATCTG |  |
| *psbD,*（D2 protein），6000044 | TTCATAACTGGACGCTGAAC | GCGGTGACCATTGAATAAGT |  |
| *ATP b*，03G013900 | CTCATGGTCACCCTCGACAAA | TCTTGTTAAGCGCTGCCGA |  |
| *NADPH*，14G015400 | CGGTGGCCTGAAGATAACCAA | GCCGACTGCCTCTATGAAAA |  |
| *Cyt b6f*，14G016000 | TGCTGCTTCTGGGTGCTATTT | CCTTGAGTTAGGGTTCGGTCTC |  |
| *MeCSK*，13G022900 | CAGTTGGACCTCTTTCGCAG | ATCTGATGCATTGGCTCCAGG |  |
| *MeFD*，15G029300 | AGGCATTGGACTCTGGACTG | CAGTGCATACCCACGTTCCA |  |
| *MeLHCB2.1*，01G175700 | AAGATTGGTTCCTTCGGCGG | AACCAGCAGTGTCCCATCCA |  |
| *MePPDK*，03G188300 | GTCGGCGATGAGAGGAATGT | TTGTGCTCGGATTCTGCCTT |  |
| *MePEPC*，02G091300 | CGCACAGATGAGATCCGAAG | CATCACGATCACCACCCATC |  |
| *MeNADP-Me*，11G034000 | GCGTCTTGATGGAGAACACA | AGTCCTTTGTTGTAGCGTGG |  |
| *MeRuBPCo* 10G027600 | ACTCAAGAGAAGGACCCCCA | GGGCAGCTTGACCGTAGAAT |  |
| *MeRuBPCo*1 05G137400 | TACCACCCGCAAAACCAACT | AGCGACCATCGTAGTACCCT |  |
| *MeFT2*(SP6A) 13G000800 | GGGGATGATTTGAGGACCTTCT | GCCGTGGACTCTCATAGCAT | Flowering and development of storage root |
| *MeFT1* 12G001600 | CATGTCACTAATGGCTGTGAGC | GATTTGGGTCACTGGGGCT |  |
| *MeBEL5* 09G045600 | TAAGGAGCAAGAGCGGAATGG | TGCTACAGGGGAAGTTGAAGC |  |
| *MeTFL2* 13G011900 | ACATTTGGAAGGGAGGTGGT | GCAGCTACAGGAAGACCAAGG |  |
| *MeCOL1*  06G028200 | AGAGTCATAGCACGGTGACA | CCCTTCTGGAACCACTCCTA |  |
| *MeHNL10-like*, 13G092100 | GCTGCTGATAGATACGTTGAC | GCCCAGCTTCATTGTTGTAA | Cyanide metabolism related |
| *MeCYP79D2*, 12G133500 | GCAGCGAACATTTACAGACG | CCTCCCACAAAGACGTTCAT |  |
| *MeCYP71E7*  *12G132900* | GCAGCGAACATTTACAGACG | ATCGATTCCTCCCACAAAGAC |  |
| *MeUTG85K4*, 12G132300 | AATCTCAACCCAGAAGCCG | GAAATCAGGAAGACCGTCGAT |  |
| *MeGTR*, 15G176100 | ATGCTCAGCCTCACAATCTG | CGTTGTAAGAGCCAAATGCC |  |
| *MeCGTR1*, 15G180400 | TGTGCTGTCTTCTTCATCGG | GTTCAGTGTGAGAGAGCCTG |  |
| *MeMATE1*, 16G007900 | TGAGAGTGAGCAACGAGCTA | GAATGAGGGTGACAGCCAAG |  |
